# Dimerization of MilM is essential for catalyzing the pyridoxal-5’-phosphate (PLP)-dependent Cγ-hydroxylation of L-arginine during mildiomycin biosynthesis

**DOI:** 10.64898/2026.02.25.707955

**Authors:** Simita Das, Yashwanth Naik, Udita Mishra, Mahima Ganguly, Badri Nath Dubey, Suvamay Jana, Nilkamal Mahanta

## Abstract

MilM from the mildiomycin biosynthetic pathway is a PLP-dependent enzyme, previously annotated as an aminotransferase, but has recently been demonstrated as L-arginine oxidase cum *C*-hydroxylase. Here, we report detailed biochemical, biophysical, structural modeling, and molecular dynamics simulation-based investigations of MilM from *Streptoverticillium rimofaciens* B-98891 to elucidate the mechanisms of substrate binding, catalysis, and the role of the active site residues involved in these processes. Our experimental findings confirmed that MilM functions as a stable homodimer, requiring the PLP cofactor and molecular oxygen to transform the L-arginine substrate into 5-guanidino-4-hydroxy-2-oxovaleric acid and 5-guanidino-2-oxovaleric acid through the intermediacy of a possible superoxide radical anion species, while generating H_2_O_2_ and NH_3_ as reaction by-products. Our labeling studies also established that the hydroxyl group in the product is obtained from the solvent, water. The structure-based three-dimensional modeling and simulation of MilM coupled with site-directed mutagenesis further confirmed that, in addition to the catalytic residues Lys232 and His31, active site residues from both the protomers are crucial for stabilization of the PLP cofactor (Ser92, Phe116, Asn164, Asp195, Lys240 from chain A and Tyr89 from chain B) and the substrate (Thr14, Glu17, Asn118, and Arg364 from chain A and Thr259, Ser260 from chain B). Moreover, the molecular dynamics simulation uncovered a dimer-mediated alternating lid mechanism in which large-scale, concerted motions of the dimer interface helices reciprocally expose and occlude the two active sites. This see-saw-like dynamics controls the substrate entry and product release through transiently formed tunnels, while preserving a catalytically protective environment, a critical phenomenon previously left unnoticed in similar PLP-dependent oxidases/hydroxylases. Overall, these findings provide new insights into the substrate/cofactor stabilization and the catalytic mechanism of MilM, a recent member of an emerging family of remarkable PLP-dependent oxidases and help us decode a key puzzle in the mildiomycin biosynthetic pathway.

## INTRODUCTION

Nucleoside natural products, owing to their distinctive chemical scaffolds and unique biological properties, have found broad applications in medicine and agriculture as antiviral,^1,2^ antifungal,^3,4^ anticancer,^5^ and antibacterial agents.^6,7^ Several N-nucleosides have been clinically applied to treat viral infections, including Hepatitis C virus (HCV) and human immunodeficiency virus (HIV).^1,5^ Mildiomycin, **1** (**Figure 1A**), is a potent broad-spectrum antibiotic isolated from *Streptoverticillium rimofaciens* ZJU5119 (B-98891), which selectively targets fungal protein synthesis.^8–12^ It belongs to the peptidyl N-nucleoside family, including blasticidin S (from *Streptomyces griseochromogenes*), arginomycin (from *Streptomyces arginensis*), cytomycin (from *Streptomyces* sp. HKI-0052), and blasticidin P10 (from *Theonella swinhoei*), which exhibit broad inhibitory activity against both prokaryotes and eukaryotes.^13,14^ Among them, mildiomycin is particularly notable for its strong fungicidal properties. It is used commercially in Japan and Europe as an agricultural fungicide due to its low toxicity to mammals and marine life.^12^ It primarily suppresses powdery mildew, a devastating fungal disease affecting staple crops such as rice, wheat, barley, millet, vegetables, and ornamentals, which poses a major threat to global agricultural economy.^15–20^ Additionally, it is effective against rice blast fungi, another destructive pathogen of cereal grains. Hence, mildiomycin serves as a bio-fungicide in agriculture and horticulture, offering a safer alternative to harmful chemical pesticides.^15–21^ Beyond its agricultural use, it inhibits other microorganisms including *Mycobacterium phlei* and *Rhodotorula rubra*, while showing minimal inhibition of DNA or RNA synthesis in mammalian and plant systems, thereby conferring selectivity with negligible host toxicity.^8–12^ In terms of mode of action, mildiomycin acts by blocking the peptidyl transferase region of the large ribosomal subunit, thereby halting protein biosynthesis.^8–12^

**Figure 1:**
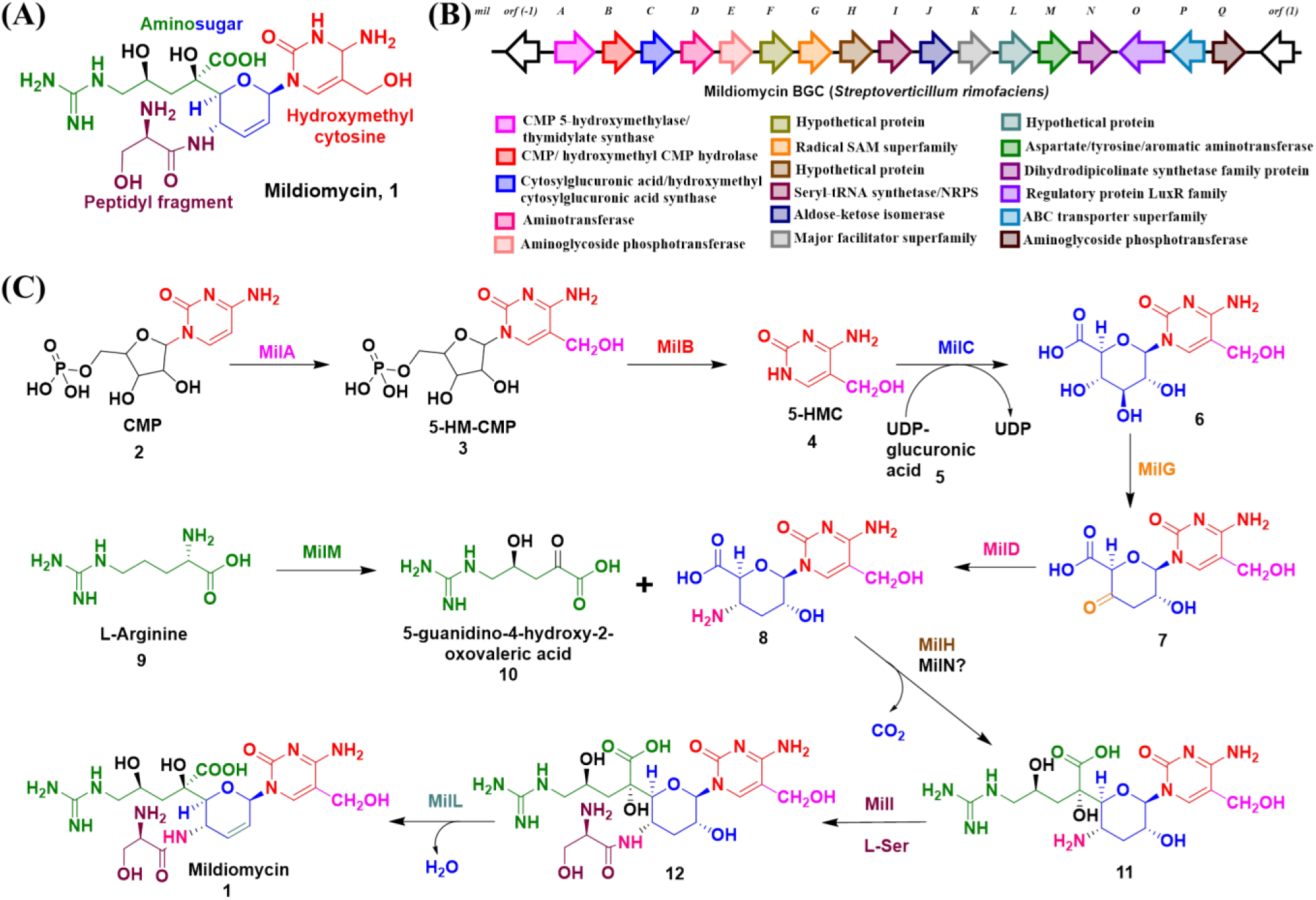
Structure and biosynthetic pathway of mildiomycin in *Streptoverticillium rimofaciens*. **(A)** Chemical structure of mildiomycin, **1**, highlighting its four moieties: hydroxymethyl cytosine (red), dihydropyranose sugar ring (blue), modified arginine side chain (green), and an appended L-serine (purple). **(B)** Organization of the mildiomycin biosynthetic gene cluster (*milA*-*milQ*) in *S. rimofaciens*, with color-coded annotations for predicted enzyme functions. **(C)** Proposed biosynthetic pathway starting from CMP, **2**. MilA-MilD assembles the nucleoside core (**3-8**), while MilM converts L-Arg (**9**) to the hydroxylated ketoacid (**10)**. Subsequent steps are likely to involve MilH/MilN, MilI, and MilL enzymes to generate the final product, mildiomycin (**1**), through intermediates **11** and **12**.

Structurally, this family is unified by a nucleoside core comprising a cytosine base connected to an unsaturated aminohexuronic acid via a glycosidic linkage, which is further diversified by various modifications.^9,13,14^ Mildiomycin, **1** (**Figure 1A**) exhibits an unusual structure composed of 5-hydroxymethylcytosine (5-HMC), a 4-aminoacyl-4-deoxyhexopyranose uronic acid, a 5-guanidino-2,4-dihydroxyvalerate side chain, and an appended L-serine moiety.^8–12^ The biosynthetic gene cluster (BGC) responsible for its production has been found in *S. rimofaciens* (**Figure 1B**), and a preliminary pathway that initiates from cytidine 5’-monophosphate (CMP) has been proposed (**Figure 1C**) based on *in vivo* genetic knockouts, characterization of blocked intermediates, *in vitro* studies, and bioinformatics.^22–26^ However, only a few early enzymes have been partially characterized biochemically,^22–29^ while detailed mechanistic and structural investigation of several key enzymatic steps and optimization of the biosynthetic pathway remains to be achieved.

According to the current working hypothesis, the biosynthetic pathway (**Figure 1C**) begins with hydroxymethylation of CMP (**2**) by MilA, a homolog of thymidylate synthase, to yield 5-hydroxymethyl CMP (5-HM-CMP, **3**). Recent work on MilA confirmed that it is dependent on N5, N10-methylenetetrahydrofolate and uses only CMP as a substrate, but no mechanistic studies have been reported.^26,27^ Next, MilB^24,25^, a nucleosidase, was postulated to cleave the C-N bond in **3** to form 5-HMC (**4**). *In vitro* biochemical and structural characterization of MilB has been reported.^24,25^ In the following step, MilC which shares homology to cytosyl glucuronic acid synthase, couples UDP-glucuronic acid (**5**) with **4** to form the nucleoside sugar skeleton, **6.**^23^ MilC was characterized biochemically, however detailed mechanistic investigation is yet to be conducted.^23^ Thereafter, MilG (homolog of BlsE from blasticidin S biosynthesis),^30–32^ a putative radical S-adenosyl methionine (SAM) enzyme, is proposed to perform a remarkable 4’-oxidation/3’-dehydration of **6** to form **7**. Thus far, no *in-vitro* studies on MilG have been reported. Then, MilD, an uncharacterized PLP-dependent aminotransferase (ATase), is postulated to convert the C4’-oxo group to C4’-amino group to form 3’,4’-dideoxy-4’-aminoglucuronic acid containing nucleoside, **8**. Interestingly, **1** has an arginine side chain appended to 5’ carbon of the sugar fragment. MilM, which was previously hypothesized to be an ATase,^13^ but recently been elucidated to be a hydroxylase,^28^ catalyzes the formation of 5-guanidino-4-hydroxy-2-oxovaleric acid (**10**) directly from L-arginine (L-Arg, **9**), thus eliminating the need for a separate hydroxylation catalyst (such as MilJ, a putative oxidase proposed earlier^23^) in this pathway. Thereafter, the coupling of **8** to **10** to form **11** remains enigmatic and could involve MilH and MilN enzymes. MilH may catalyze a decarboxylative, intermolecular aldol-type reaction to attach **8** to **10,** and MilN, which is similar to 4-hydroxy-tetrahydrodipicolinate synthases, could also be involved in this reaction.^13^ Next, MilI which contains an N-terminal region that resembles with the T domains of NRPS and a C-terminal region which resembles with the seryl-tRNA synthetases is likely to be involved in amide bond formation by attaching L-serine (L-Ser) to **11** to generate **12**. Finally, MilL is postulated to perform the 2’-O dehydration of **12** to form mildiomycin, **1**.^13^

MilM represents a fold type I PLP-dependent oxidase identified in a nucleoside natural product biosynthetic pathway that catalyzes the direct conversion of L-Arg (**9**) into 5-guanidino-4-hydroxy-2-oxovaleric acid (**10**). MilM shows 37% identity with MppP from *Streptomyces wadayamensis* (annotated as a PLP-dependent ATase), which was reported to convert L-Arg to **10** during the biosynthesis of unrelated metabolites, such as L-enduracididine, a non-proteinogenic amino acid found in a number of bioactive peptides.^33,34^ Another PLP-dependent hydroxylase enzyme, RohP from *Streptomyces cattleya* (60% identity with MilM) involved in azomycin biosynthesis, also catalyzes the conversion of L-Arg to form the hydroxylated ketoacid (**10**).^35,36^ These PLP-dependent fold type I enzymes employ O_2_ as a co-substrate to oxidize organic substrates^37,38^. Very recently, Edwards *et al.* reported that MilM from *S. rimofaciens* functions as a similar PLP-dependent oxidase/hydroxylase.^28^ Through enzyme assays, biochemical, and stoichiometric analyses, they showed MilM converts L-Arg (**9**) to **10**, with a 1:1 stoichiometry of O_2_ to H_2_O_2_. Isotopic labeling (50% H_2_^18^O) confirmed the hydroxyl oxygen originates from water, and Ultraviolet-Visible (UV-Vis) spectroscopic studies revealed distinct quinonoid intermediates, providing preliminary mechanistic evidence for an O_2_-dependent PLP-mediated hydroxylation pathway.

O_2_-dependent PLP oxidases/hydroxylases represent a highly unusual and sparsely characterized enzyme class. To date, only three functionally validated members performing *C*-hydroxylation reaction on L-Arg have been reported: MppP,^33,34^ RohP,^35^ and MilM.^28^ Despite recent advances, several key mechanistic questions remain unresolved, including whether the catalysis proceeds through a superoxide radical anion intermediate or a hydroperoxy intermediate for the critical C-H activation step remains to be established. Furthermore, particularly in MilM, the identification of active site residues which bind substrate/PLP cofactor as well as the residues which participate in catalysis is yet to be elucidated. Moreover, although homologous enzymes are typically tetramer^33,34^ or dimer,^35^ the functional necessity of dimerization/oligomerization and the role of each protomer in cofactor coordination, substrate recruitment, and product release have not been established for this class of enzymes.

Herein, we report an integrated biochemical, mutational, computational, and molecular dynamics simulation-based investigation that defines the structural and mechanistic basis of MilM from *S. rimofaciens* B-98891. Our independent *in vitro* biochemical assays demonstrate O_2_-dependence of MilM with hydrogen peroxide and ammonia as by-products, validating it as a bonafide PLP-dependent oxidase. Inhibition of the enzymatic reaction by superoxide dismutase further supports the intermediacy of a possible superoxide radical anion species during catalytic turnover. Experimental studies supported by structural modeling and molecular dynamics (MD) simulation established that MilM exists as a functional dimer. Site-directed mutagenesis (SDM) of active site residues from both protomers, coupled with kinetic and substrate-scope analysis, identifies the key determinants of PLP binding, substrate recognition, and product formation in MilM. Finally, MD simulations reveal that MilM employs a potential dynamic lid-opening and closing mechanism to control active-site access, demonstrating how the dimer architecture choreographs substrate’s entry and product’s release without the dissociation of the monomeric subunits. Taken together, these results establish MilM as a dimer-dependent PLP oxidase cum hydroxylase that utilizes a plausible substrate-gated conformational switching process to achieve a catalytically competent active site and use reactive oxygen chemistry to catalyze an intriguing regiospecific *C*-hydroxylation of L-Arg. We believe our studies will strengthen the mechanistic framework for this emerging class of PLP-dependent enzymes and broaden our understanding of nucleoside biosynthesis for further optimization of the enzymatic machinery and engineering of the pathways.

## MATERIALS AND METHODS

### Materials

Reagents used in molecular biology procedures, nickel nitrilotriacetic acid (Ni-NTA) resin and superoxide dismutase (SOD) and were obtained from New England Biolabs (NEB), Ipswich, MA (USA). *S. rimofaciens* strain B-98891 (ATCC 31120) was purchased from the American Type Culture Collection (ATCC) through HiMedia Laboratories Pvt. Ltd. (India). *Escherichia coli* DH5α and BL21 (DE3) strains were obtained from the NCIM culture collection, Pune (India), and used to maintain the plasmid and to overproduce protein, respectively. Oligonucleotides were purchased from Eurofins Genomic India Pvt. Ltd., oligonucleotide synthesis department, Bangalore (India). All plasmid inserts and mutants were sequenced at Eurofins Genomic India Pvt. Ltd., sequencing department, Bangalore (India). Isopropyl β-D-1-thiogalactopyranoside (IPTG), α-ketoglutarate (α-KG), dansyl chloride (DNS-Cl) and H_2_^18^O were purchased from BLD Pharmatech Pvt. Ltd. (India), and D_2_O was purchased from Tokyo Chemical Industry (TCI) Co., Ltd., Japan. Luria-Bertani (LB) broth, L-Arg, D-Arg, L-alanine (L-Ala), L-lysine (L-Lys), L-glutamate (L-Glu), L-glutamine (L-Gln), nicotinamide adenine dinucleotide (NADH), H_2_O_2_, N-(2-Hydroxyethyl)-Piperazine-N’-(2-ethanesulfonic acid) (HEPES), and other reagents were obtained from Sisco Research Laboratories (SRL) Pvt. Ltd., India. Bovine serum albumin (BSA), catalase, Horseradish peroxidase (HRP), 2, 2’-azinobis-[3-ethylbenzothiazoline-6-sulfonic acid] (ABTS), formic acid (FA), and Glutamate dehydrogenase (GDH) were purchased from Sigma-Aldrich, USA. Ultra centrifugal filters (30 kDa, Amicon) were obtained from Merck-Millipore, USA, and nanosep centrifugal filters (30 kDa) were purchased from Pall Life Sciences, USA.

### Genomic DNA Isolation

Genomic DNA from the mildiomycin producing *S. rimofaciens* B-98891 strain was isolated using the DNeasy UltraClean Microbial Kit (Qiagen, USA). The concentration of genomic DNA was found to be 990 ng/µL (**Figure S1**).

### Cloning of the *milM* gene

Molecular biology experiments were performed using standard techniques.^39,40^ For polymerase chain reaction (PCR) amplification of *milM*, genomic DNA from *S. rimofaciens* was used as the template. Q5 high-fidelity DNA polymerase (NEB) was used for gene amplification with specific forward and reverse primers designed with BamHI and NotI restriction sites (**Table S1**). The product containing the desired gene was purified and appropriate restriction enzymes were used for digestion, re-purified, and then ligated using T4 DNA ligase (NEB) into purified pET28 vector which was digested in the same way. Ligated DNA was transformed into competent *E. coli* DH5α cells, selected on LB agar plates containing 50 µg/mL kanamycin (SRL) and confirmed by DNA sequencing.

### Construction of MilM variants

SDM was carried out using the QuikChange protocol with Q5 high-fidelity DNA polymerase for creating the MilM variants: T14A, E17A, H31A, Y89A, S92A, F116A, N118A, N164A, D195A, K232A, K240A, K232A-K240A, T259P-S260P, and R364A. pET28*-milM-*wild type (WT) plasmid was used as the template for SDM using the QuikChange method with the corresponding pairs of primers listed in **Table S1**. The template DNA was cleaved with DpnI (NEB) and transformed into *E. coli* DH5α competent cells. The colonies were grown to extract mutated plasmids, and the newly constructed plasmids were verified for correct point mutations by DNA sequencing.

### Over-expression and purification of MilM-His_6_ tagged protein and its variants

Chemically prepared competent *E. coli* BL21 (DE3) cells^41^ were transformed with the gene constructs of interest and grown for 12-15 hours on LB agar plates containing 50 µg/mL kanamycin at 37 °C. Colonies were inoculated in 20 mL of LB media having 50 µg/mL kanamycin and grown for 16-18 hours at 37 °C. This pre-inoculum was then used to inoculate 2 L of LB media having 50 µg/mL kanamycin and incubated at 37 °C with 200 rpm to OD_600_ of 0.4-0.6, then later cooled at 16 °C for 15 min. Expression of the protein was induced by adding 500 µM IPTG and the culture was incubated for 12-16 hours at 16 °C at 150 rpm. Thereafter, cell pellets were separated from media by centrifugation (4200 rpm for 30 min), washed with phosphate-buffered saline (137 mM NaCl, 2.7 mM KCl, 10 mM Na_2_HPO_4_, and 1.8 mM KH_2_PO_4_), and harvested by centrifugation (4000 rpm for 30 min). The resulting cell pellet was immediately frozen and stored at -80 °C until further use. For protein purification, first harvested cells were resuspended in the lysis buffer [20 mM HEPES pH 7.5, 500 mM NaCl, 10 mM Imidazole, 2.5% (*v/v*) Glycerol, Triton X-100 (*v/v*)]. Cells lysis was achieved by sonication at 4 °C using alternating 30 sec on-30 sec off cycle for a total of 1 hour at 40 % amplitude settings. Cellular debris was suspended by centrifugation at 15000 rpm for 1 hour at 4 °C. Thereafter, the resulting supernatant was filtered through a 0.45 µm syringe filter and added onto a Ni-NTA gravity-flow column (5 mL resin) that had been pre-equilibrated with the lysis buffer. After washing the column with 100 mL of wash buffer [lysis buffer lacking Triton X-100], bound His_6_-tagged protein was eluted with 15 mL elution buffer [20 mM HEPES, 300 mM NaCl, 300 mM imidazole, 2.5% glycerol (*v/v*)]. The eluent was then concentrated with a 30 kDa molecular weight cutoff Amicon Ultra centrifugal filter (Millipore), followed by buffer exchange with storage buffer [50 mM HEPES pH 7.5, 300 mM NaCl, 10% glycerol (*v/v*)]. Final protein concentrations were determined spectrophotometrically at 280 nm using the theoretical extinction coefficient of MilM (38390 M^-1^cm^-1^, calculated using the Expasy ProtParam tool; http://web.expasy.org/protparam/). Protein purity was visually verified by Coomassie dye-stained sodium dodecyl sulfate-polyacrylamide gel electrophoresis (SDS-PAGE) analysis (**Figure S2**).

### Size Exclusion Chromatography (SEC) method

SEC analysis was performed using a Fast Performance Liquid Chromatography (FPLC) instrument (Cytiva AKTA Start Chromatography System, Department of Chemistry, IIT Dharwad). HiPrep 16/60 Sephacryl S-100 High Resolution column (1 x 120 mL) was used for the analysis. This column was equilibrated for 6 hours with buffer 10 mM HEPES (pH 7.5) at a flow rate of 0.5 mL/min before injecting sample. 100 µL of 20 mg/mL purified MilM-His_6_-WT was prepared in the same buffer, passed through a 0.2 µm filter, and injected through a loop into the equilibrated column at a flow rate of 1 mL/min with absorbance detection at 280 nm. BSA, CckA and BolA proteins prepared in similar way were used as standard/known references for our protein and the FPLC raw data were analyzed using Origin software.

### UV-Vis spectroscopic method

The MilM protein was analyzed by Cary 5000 UV-Vis spectrophotometer facility at IIT Dharwad for monitoring PLP-bound protein characteristic peaks. To determine the protein: PLP ratio, UV-Vis spectrum of 100 μM of denatured MilM protein (by heating at 95 °C) in 50 mM HEPES pH 7.5 was recorded, which was later compared with the UV-Vis spectra of different concentrations of standard PLP (10-500 μM). Similarly, the UV-Vis spectroscopic analysis of 50 µM MilM protein with substrate L-Arg (1 mM) in 50 mM HEPES pH 7.5 was also determined to verify the identity of the intermediates formed in the MilM reaction. The raw data were analyzed using Origin software.

### *In vitro* enzymatic assays for MilM

For evaluating MilM activity as an ATase, a reaction mixture (200 µl) comprising of 50 mM HEPES pH 7.5, 1 mM L-Arg, 1 mM α-KG, and 50 µM purified MilM-His_6_ was reacted at room temperature (r.t.) for 2 hours. MilM reaction mixture was quenched by heating for 5 min at 95 °C and then subjected to centrifugation at 13000 rpm for 1 min to separate the precipitated protein. The resulting supernatant solution was used for analysis using various analytical methods.

For evaluating MilM activity as an oxidase, a reaction mixture (200 µl) comprising of 50 mM HEPES pH 7.5, 1 mM L-Arg, and 50 µM purified MilM-His_6_ (with or without 1 mg of catalase) was incubated at r.t. for 2 hours. MilM reaction mixture was quenched by heating for 5 min at 95 °C and then centrifuged at 13000 rpm for 1 min to separate the precipitated protein. The resulting supernatant solution was then analyzed using various analytical methods. MilM reaction under anaerobic conditions was performed using a glovebox (Coy Laboratories, USA) to elucidate the role of molecular oxygen in the reaction.

### Dansyl chloride (DNS-Cl) derivatization method

A reaction mixture of 100 µl composed of 50 mM HEPES pH 7.5, 1mM L-Arg, and purified 50 μM MilM-His was incubated at r.t. for 3 hours. The protein was removed by heating for 5 min at 95 °C and centrifuged at 13000 rpm for 1 min to separate the precipitated proteins. The resulting supernatant was made basic (pH around 9-10) by adding NaOH, then added 10 mM of freshly prepared DNS-Cl in acetonitrile (ACN) and kept for 1 hour at 37 °C in dark.^42^ The reaction was then quenched with 0.1 M HCl and further used for HPLC analysis. Similar method for derivatization of standard 1mM L-Arg with DNS-Cl was also performed for the control assay.

### Reverse Phase High-performance Liquid Chromatography (HPLC) method

HPLC analysis (Shimadzu Phenomenex HPLC system at the Department of Chemistry, IIT Dharwad) was performed to detect the PLP and derivatized products of the MilM reaction. The MilM reaction mixture was passed through a 0.2 µm filter, and 15 µL of the filtrate was injected into shim-pack solar C18-column (dimension of 4.6 mm I.D × 250 mm, particle size 5 µm) for HPLC analysis. Linear gradient LC elution was performed at a flow rate of 1 mL/min with absorbance detection at 254 nm with oven temperature maintained at 35 °C. Solvent A is HPLC grade water in 0.1 % FA; solvent B is HPLC grade ACN in 0.1 % FA. The gradient method used is as follows: 0 min: 20% B; 2 min: 20% B; 10 min: 60% B; 15 min: 80% B; 18 min: 80% B; 25 min: 60% B; 30 min: 20% B. HPLC raw data were analyzed using Origin software.

### Liquid Chromatography-Mass Spectrometry (LC-MS) method

High-resolution electrospray ionization mass spectrometry (HR-ESI-MS, Waters Xevo G3 QTof LC-MS system) analyses were performed at the mass spectrometry facility, Department of Chemistry, IIT Dharwad. Sample preparation was performed as mentioned in the HPLC method above and 1 µL sample was injected through a C18-column (dimension of 2.1 x 50 mm, 1.7 µm particle size) with oven temperature maintained at 40 °C. The LC elution was performed at a 0.2 mL/min flow rate. Solvent A is LC-MS grade water in 0.1 % FA; solvent B is LC-MS grade methanol in 0.1 % FA; and solvent C is 5 mM LC-MS grade ammonium acetate buffer (0.2 mL/min flow rate). The gradient method used is as follows: 0 min: 100% C; 1 min: 100% C; 2 min: 48% A, 12% B, 40% C; 3 min: 50% A, 20% B, 30% C; 4 min: 30% A, 10% B, 60% C; 6 min: 100% C. Mass spectrometric data acquisition was conducted in both positive and negative ESI modes. Ion source parameters were as follows: temperature 100 °C, cone gas flow rate 50 L/hr, and capillary voltage 1000 V for negative mode and 2500 V for positive mode. Data were collected over a mass range of *m/z* 140-195. Raw data was analyzed using MassLynx software (Waters).

### Nuclear Magnetic Resonance (NMR) Spectroscopic method

For NMR analysis of the MilM assay, first 2 mM L-Arg was added in 50 mM sodium phosphate buffer (pH 7.5) in 100% D_2_O (total of 700 µl) in NMR tube and ^1^H-NMR spectrum was recorded using a Jeol ECZL 400 MHz NMR instrument at the Sophisticated Central Instrumentation Facility (SCIF) of IIT Dharwad. Then, 100 µM of purified MilM was added in the same NMR tube and recorded the ^1^H-NMR after 3 hours. The NMR Free Induction Decay (FID) data was analyzed using JEOL delta v6.0 NMR software.

### *In vitro* MilM assay in the presence of SOD

MilM assays were performed in the presence of SOD to examine its effect on the enzyme-catalyzed oxidation of L-Arg. Reaction mixtures (200 µL) having50 mM HEPES buffer (pH 7.5), 1 mM L-Arg, and 50 µM purified MilM-WT were supplemented with varying concentrations of SOD (0.5, 1.25, 2.5, 5.0, 7.5, and 10 mg/mL). The reactions were incubated at r.t. for 2 hours and subsequently quenched by heating at 95 °C for 5 min. Precipitated proteins were removed by centrifugation at 13,000 rpm for 1 min, and the resulting supernatants were subjected to LC-MS analysis to detect MilM reaction products.

For quantification of residual L-Arg in each reaction, aliquots from the above mixtures were derivatized with 10 mM DNS-Cl for 1 hour at 37 °C in the dark and analyzed by HPLC, monitoring the DNS-Arg peak at retention time (Rt = 3.5 min), which corresponds to unreacted L-Arg. To determine the concentration of remaining L-Arg (or DNS-Arg), a calibration curve was generated by derivatizing standard L-Arg solutions (10, 25, 50, 100, 250, 500, 750 μM) under identical conditions and plotting the resulting peak areas against concentration. This standard curve was used to calculate the residual L-Arg concentrations in the MilM reactions performed with different amounts of SOD (**Table S2**).

### MilM and ABTS-HRP coupled assay

Reaction mixtures of 100 µl composed of 50 mM HEPES pH 7.5, purified 10 µM MilM-His, 10 mM ABTS and 10 ng HRP were incubated at r.t. on a 96-well plate having different concentrations of L-Arg (10 - 1000 µM). This reaction was used to detect the by-product, hydrogen peroxide (H_2_O_2_), produced during the MilM reaction, where HRP oxidizes ABTS in the presence of H_2_O_2_, producing a bluish-colored product (ABTS•⁺).^35,43^ For comparison, similar reactions were conducted with standard H_2_O_2_ of different concentrations (10 - 1000 µM), 10 mM ABTS and 10 ng HRP (without MilM-His and L-Arg).

### UV-Vis analysis of MilM and GDH coupled assay

To detect ammonia (NH_3_) as a byproduct of MilM catalyzed transformation, a reaction mixture (400 µl) composed of 50 mM HEPES pH 7.5, 1 mM L-Arg, purified 50 μM MilM-His, 3.5 mM α-KG and 250 μM NADH was incubated at r.t. for 20 minutes prior to the addition of 100 ng GDH. The reaction mixture was analyzed by Cary 5000 UV-Vis spectrophotometer facility at SCIF, IIT Dharwad to record the decrease in the NADH absorbance at 340 nm for an interval of 2 minutes for time duration of about 15 minutes. The raw data were analyzed using Origin software.

### Steady state kinetic assay

The by-product H_2_O_2_, produced during the MilM reaction, was used by HRP to oxidize ABTS, producing a bluish-colored product (ABTS•⁺).^35,43^ The rate of ABTS•⁺ formation was monitored spectrophotometrically at 418 nm using a Synergy HTX multi-mode reader (Agilent Technologies) at the Department of Chemistry, IIT Dharwad. Reactions were conducted in a 100 µL mixture containing 10, 25, 50, 100, 250, 500, 750 µM L-Arg, 1 µM MilM (MilM-WT and MilM-variants), 1 mM ABTS, 10 ng HRP in 50 mM HEPES pH 7.5. The reaction was initiated by adding the MilM enzyme followed by monitoring the formation of ABTS•⁺ spectroscopically at 418 nm for a time of 30 minutes with 1 minute interval. Absorbance values were compared against a standard calibration curve for H_2_O_2_, and the calculated concentrations of ABTS•⁺ (equivalent to H_2_O_2_ formed in the MilM reaction) were plotted against time for each substrate concentration. Initial reaction rates were determined from the slopes of these plots and fitted to the Michaelis-Menten equation using Kaleida graph 4.0 to generate the steady-state kinetic parameters (*k*_cat_, *K*_m_, and *k*_cat_/*K*_m_) for the WT and its variants. All the assays were performed in triplicate to ensure reproducibility.

### Substrate scope of MilM

For evaluating MilM activity as an oxidase for different substrates (D-Arg, L-Lys, L-Glu, L-Gln, and L-Ala), reaction mixtures of 200 µl comprising of 50 mM HEPES pH 7.5, 1 mM substrate, and 50 µM purified MilM-His_6_ were incubated at r.t. for 2 hours. MilM reaction mixtures with different substrate analogs were quenched by heating for 5 min at 95 °C and then centrifuged at 13000 rpm for 1 min to separate the precipitated protein. The resulting supernatant solutions were analyzed for any product formation using the LC-MS method as mentioned earlier.

### Three-dimensional (3D) modeling of MilM and PLP-L-Arg complexes

The 3D structure of MilM (UniProt: H9BDX2) was modeled as a dimeric assembly using AlphaFold-Multimer v3.^44,45^ The highest-confidence-ranked model was selected for all subsequent analyses. To model the L-Arg-PLP and D-Arg-PLP complexes, the predicted MilM dimer was structurally aligned to the experimentally determined L-Arg-PLP-bound RohP complex (PDB ID: 6C3C) and the D-Arg-PLP-bound MppP complex (PDB ID: 5DJ3) in PyMOL (https://pymol.org/2/), respectively. Following superposition, L-Arg-PLP and catalytically relevant water molecules, including conserved interface-associated waters, were transferred from the RohP structure to the corresponding binding pocket of the MilM model. The resulting complexes were slightly adjusted manually in PyMOL and Coot^46^ to relieve steric clashes while preserving the original ligand/substrate geometry.

### Dimer interface analysis by PISA

Dimer interface energetics and assembly stability were evaluated using the Protein Interfaces, Surfaces, and Assemblies (PISA) server.^47,48^ The AlphaFold-Multimer dimer (PDB format) was submitted to PISA (https://www.ebi.ac.uk/pdbe/pisa/), and interface properties including solvent-accessible surface area (SASA) burial, solvation free energy gain (ΔG) upon dimer formation, hydrogen bonding, and salt bridge networks were calculated using default parameters. Residues exhibiting >1 Å^2^ loss in SASA upon complex formation were classified as interface contributors. Interface architecture and residues were visualized in PyMOL, using chain-specific coloring and surface representations for figure preparation (**Table S3**).

### Dimer interface Mapping with DIMPLOT

To generate schematic representations of inter subunit interactions, the MilM dimer model was analyzed using DIMPLOT module of the LigPlot+ suite.^49^ DIMPLOT identified hydrogen bonds (donor-acceptor distance ≤ 3.5 Å) and hydrophobic contacts arising from close Van der Waals surface complementarity between opposing protomers. The resulting two-dimensional (2D) interaction diagrams were exported for illustration.

### Ligand Interaction Analysis with LigPlot+

Interactions between MilM and L-Arg-PLP were further characterized using LigPlot+.^49^ Hydrogen bond donors/acceptors and hydrophobic contacts within the catalytic pocket were mapped automatically, and annotated interaction networks were visualized in both 2D plots and 3D structural overlays.

### MD simulation

#### General

To further assess the stability of the modeled dimer complex, we carried out atomistic MD simulations for two separate systems: (1) the substrate L-Arg-PLP-bound MilM dimer, and (2) the substrate L-Arg-PLP-bound MilM monomer. For the dimer system, we considered two L-Arg-PLP molecules, each bound to the active site of the respective MilM monomer. For the monomer system, only one L-Arg-PLP bound MilM monomer unit was considered. The CHARMM-GUI webserver was employed to prepare the inputs for the MD simulation^50–52^ utilizing the structural model derived from AlphaFold, and subsequently AMBER software was used for structure minimization, heating, density equilibration, and production MD simulation.^53^

#### Protocol for system preparation

The CHARMM-GUI’s “Ligand Reader & Modeler” module was first utilized to parameterize the substrate L-Arg-PLP.^54^ Subsequently, the CHARMM-GUI’s “Solution Builder” module was used to construct the MilM-L-Arg-PLP complex (dimer and monomer separately), solvate it in a cubic box of water, and neutralize the system’s charge by adding Na^+^, and Cl^-^ counter ions to achieve a salt concentration of 0.15 M.^50^ The protonation state of the protein’s titratable amino acids were determined at pH 7.4 separately using PDB2PQR webserver,^55–58^ and the same was assigned during system preparation in the “Solution Builder” module. In this manner, the MD simulation inputs for the dimer and monomer complexes were prepared with the dimer system size of 111 Å x 111 Å x 111 Å containing ∼128,000 atoms, and the monomer system size of 85 Å x 85Å x 85 Å containing ∼57,000 atoms, respectively. Also, during system preparation, the protein was defined by the CHARMM36m force field,^51,59^ water and counter ions were described by the TIP3P force field,^60,61^ and the L-Arg-PLP substrate was described by the CGenFF force field.^62^ The identical force field parameters were maintained for subsequent structure minimization and MD simulation steps in AMBER.

#### MD simulation protocol

The solvated systems obtained from CHARMM-GUI were first minimized in AMBER for 5000 steps with the first 2500 steps using the steepest descent approach and the remaining steps using the conjugate gradient method. After minimization, the systems were heated at 298 K in the NVT ensemble over 1 ns using the Langevin thermostat.^63^ At this point, three independent 298 K simulations were started from different random number seeds for both dimer and monomer systems. These six simulations were then density-equilibrated for 1 ns in the NPT ensemble in AMBER using the Langevin thermostat and the Monte Carlo barostat at 298 K temperature and at 1 atm pressure.^64^ The equilibrated systems were then simulated for 150 ns (production MD simulation) in AMBER in the NPT ensemble, employing the same temperature, pressure, thermostat, and barostat as utilized in the preceding density equilibration phase. A uniform time step of 2 fs and a non-bonded cutoff distance of 10 Å were also utilized consistently during the heating, density equilibration, and production simulation stages. To further increase the computational efficiency, all of the hydrogen distances were kept fixed using the SHAKE algorithm during simulation.^65^

## RESULTS AND DISCUSSIONS

### MilM Protein Exists as Dimer

To perform the biochemical and biophysical characterization of MilM, we purified the protein using affinity chromatography followed by gel filtration. To gain insight into the oligomeric state of MilM, we carried out analytical gel filtration. The gel filtration profile showed that MilM eluted as a single peak at an elution volume (EV) of 38.8 mL (**Figure 2A**). To estimate the oligomeric state of MilM, we also loaded known proteins such as BSA, CckA and BolA, whose molecular weights and oligomeric states have been previously established. On comparison with the SEC-FPLC elution profiles of known proteins under identical conditions, it was observed that MilM elutes after the CckA dimer protein^66,67^ (∼146.1 kDa, EV of 35.7 mL) but elutes earlier than the BSA standard (∼66.5 kDa, EV of 42.6 mL) and BolA dimer protein^68^ (∼21.3 kDa, EV of 59.1 mL) (**Figure 2A**). These results suggest that the purified MilM protein may possibly exist as a dimer (expected molecular weight from the protein sequence is ∼44.5 kDa as calculated using the Expasy ProtParam tool). Native-PAGE analysis further confirmed that MilM exists as a dimer, migrating at an apparent molecular mass of ∼90 kDa (**Figure 2B**), whereas under denaturing SDS-PAGE conditions it appears as a monomer of ∼44.5 kDa (**Figure 2C**). Collectively, these results demonstrate that MilM exists as a stable homodimer in solution, akin to structurally characterized RohP.^35^

**Figure 2:**
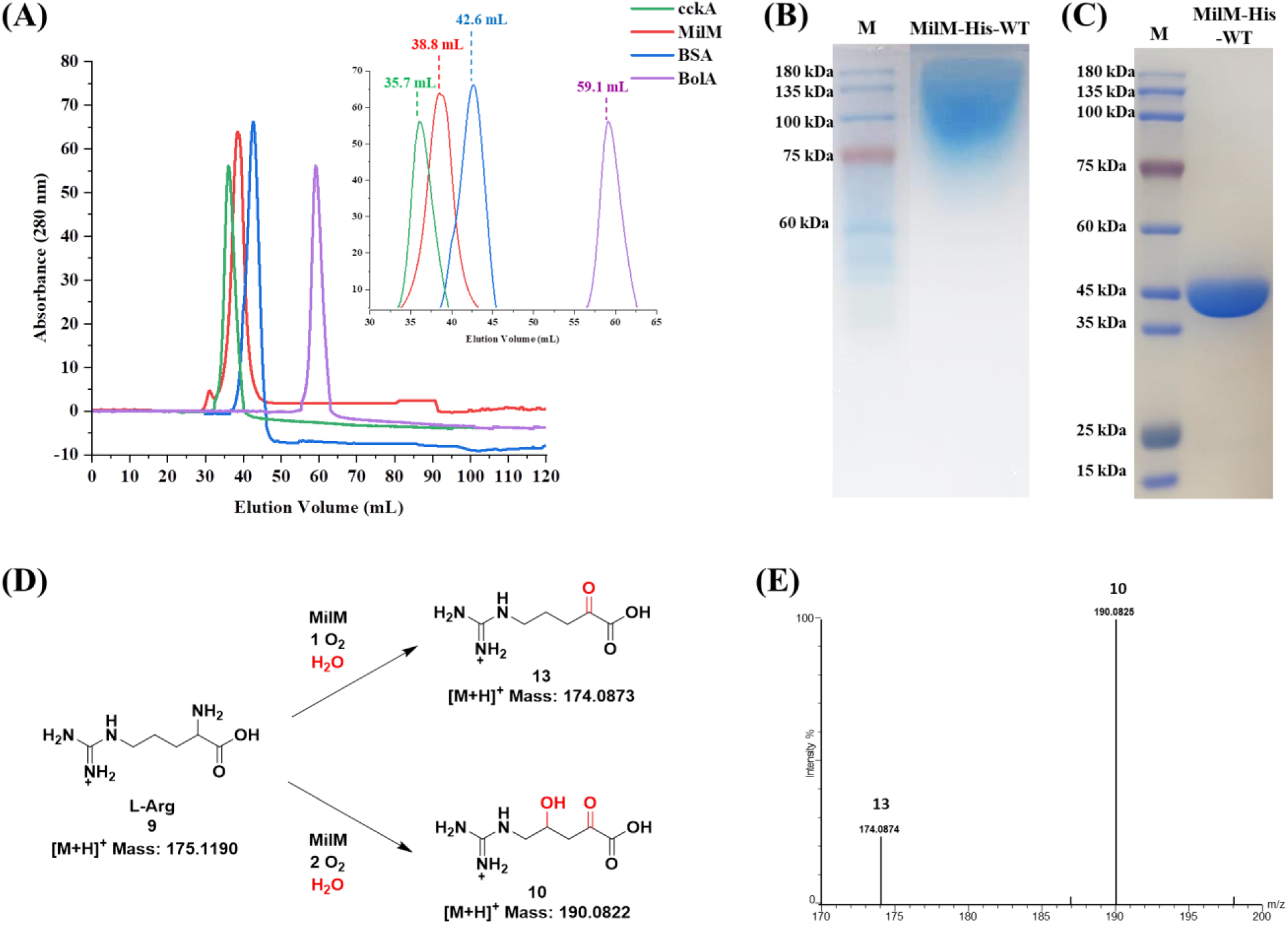
Oligomeric state of MilM and its catalytic products derived from L-Arg. **(A)** Analytical SEC-FPLC analysis of MilM revealed a major peak eluting at an elution volume (EV) of 38.8 mL, corresponding to a dimeric form when compared to CckA (∼146.1 kDa, EV of 35.7 mL); BSA standard (∼66 kDa, EV of 42.6 mL) and BolA (∼21.3 kDa, EV of 59.1 mL). **(B)** Consistently, the native-PAGE analysis showed as-isolated MilM-His_6_ migrates as a single band corresponding to a dimeric species (∼90 kDa). **(C)** SDS-PAGE gel image of denatured MilM-His_6_ indicated it as a single band corresponding to its monomeric form (∼44.5 kDa) **(D)** MilM catalyzes the oxidation of L-Arg using molecular oxygen and water to yield either 5-guanidino-2-oxovaleric acid (**13**) or 5-guanidino-4-hydroxy-2-oxovaleric acid (**10**). **(E)** In the presence of catalase, the MilM reaction shows masses (*m/z*) corresponding to compounds **10** and **13**.

### PLP is the Physiological Cofactor of MilM

Purified MilM (as-isolated) was obtained as a yellow-colored protein (**Figure S3A**), indicating the presence of a tightly bound PLP cofactor. PLP-dependent enzymes typically form a covalent Schiff base (internal aldimine) between the aldehyde group of PLP and a conserved Lys residue within the active site.^38,69^ Consistent with the formation of the internal aldimine, UV-Vis spectroscopic analysis of the purified enzyme exhibited a characteristic absorption maximum at ∼420 nm (**Figure S3B**). Denaturation of MilM by heating the protein at 95 °C released the bound cofactor as indicated by a shift in absorption to ∼392 nm in the supernatant, corresponding to the free PLP cofactor (**Figure S3B**). HPLC analysis further confirmed this assignment, as the supernatant obtained from the denatured protein sample displayed a peak co-eluting with authentic PLP standard at a retention time (Rt) of 10.8 min (**Figure S3C**). From UV-Vis spectra, the PLP: protein ratio was found to be 1:1, suggesting that 100% of MilM is cofactor bound (**Figure S3D**). The incorporation of PLP without supplementation during heterologous expression in *E. coli* also suggested a strong intrinsic affinity of MilM for the cofactor, a feature that may contribute to its catalytic efficiency and stability.

PLP-dependent oxidases typically form an internal aldimine through a conserved Lys residue, and sequence alignment of MilM with structurally characterized homologs revealed the presence of such a conserved Lys at position 232 (Lys232, **Figure S4**). To evaluate whether Lys232 mediates covalent attachment of PLP, we mutated it to Ala. To our surprise, the purified MilM-K232A variant retained its characteristic yellow coloration (**Figure S5A**), and UV-Vis analysis showed the characteristic absorbance at 420 nm corresponding to a PLP bound internal aldimine (**Figure S5B**). These results indicated that PLP remained covalently bound even in the absence of Lys232 and suggested the involvement of an alternative nucleophilic Lys residue. Inspection of the sequence alignment revealed a second conserved residue, Lys240, positioned near Lys232 and similarly conserved among related oxidases (**Figure S4**). We hypothesized that in the absence of Lys232, PLP may adopt a compensatory binding mode in which Lys240 acts as the Schiff-base forming residue. To test this possibility, we constructed the double variant (MilM-K232A-K240A) by substituting both Lys residues with Ala. In contrast to the single K232A variant, the purified double variant was completely colorless, and no characteristic absorbance corresponding to internal aldimine or free PLP was detected (**Figure S5B**), indicating the loss of the covalently bound cofactor. These results support a model in which Lys240 can serve as an auxiliary PLP-binding residue only when the primary catalytic Lys232 is absent. We hypothesize that Lys232 is the principal residue responsible for formation of the catalytically competent internal aldimine species in MilM based on the sequence conservation of this Lys residue in the structurally characterized homologous proteins, MppP (Lys221)^33^ and RohP (Lys235)^35^ (**Figure S4**). However, further structural studies on MilM will be required to confirm this hypothesis. Nonetheless, this rescue mechanism highlights unexpected plasticity of the PLP binding mode in MilM active site but reinforces Lys232 as the physiologically relevant catalytic Lys in MilM.

### MilM is a PLP-Dependent Oxidase, Not an Aminotransferase

MilM was previously proposed to be a PLP-dependent ATase based on bioinformatics prediction and as per genetic knock-out studies.^13,23^ To elucidate its biochemical function and further confirm its role in mildiomycin biosynthesis, we have independently tested the activity of MilM with L-Arg as substrate and α-KG, the latter being a common co-substrate for ATases.^42,70,71^ Reaction mixtures were analyzed by HPLC, LC-MS, and NMR spectroscopy. L-Arg was derivatized with DNS-Cl to generate DNS-Arg (**Figure S6A**), which appeared as a distinct peak at Rt 3.5 min in our HPLC analysis (**Figure S6B**). This observation was also confirmed in LC-MS in positive ion mode where DNS-Arg showed a mass at *m/z* 408.1710 (**Figure S6C**). In contrast, when DNS-Cl was added after completion of the MilM reaction (containing L-Arg and α-KG), no peak corresponding to DNS-Arg was observed, indicating complete consumption of the L-Arg substrate (**Figure S6B**). Interestingly, a similar chromatographic profile was obtained when MilM reaction was performed in the absence of α-KG (**Figure S6B**), suggesting that L-Arg is likely to be the sole substrate of MilM and that the enzyme does not require α-KG, thereby ruling out a canonical ATase activity. This was also confirmed by the absence of the DNS-Glu product in the MilM reaction, which was expected for an ATase.^42^ LC-MS analysis of the MilM reaction products in positive ion mode, showed masses of *m/z* 174.0861 and *m/z* 190.0826 which correspond to compounds **13** (oxo-product) and **10** (hydroxylated product), respectively (**Figure S7A**). Importantly, no mass corresponding to the expected by-product, glutamate was detected, and identical products were observed in the absence of α-KG in the LC-MS analysis, further supporting the assignment of MilM as an oxidase/hydroxylase rather than an ATase.^28^ In addition, we also observed additional compounds with masses of *m/z* 146.0893 (**14**) and *m/z* 162.0874 (**15**). We propose that these were formed due to the non-enzymatic reaction of **13** and **10** (**Figures S7B** and **S7C**), respectively, with H_2_O_2_ released during the catalytic cycle as observed in case of RohP.^35^ To confirm that **14** and **15** were formed because of a non-enzymatic pathway and not due to MilM mediated catalysis, we performed the MilM reaction in the presence of catalase,^35^ which is known to decompose H_2_O_2_. LC-MS analysis of this reaction mixture demonstrated masses corresponding to only two products **13** and **10** at *m/z* 174.0874 and *m/z* 190.0825, respectively, in a ratio of ∼1:3 (**Figures 2D**, **2E**, and **S7D**). Finally, ^1^H-NMR analysis of the MilM assay further confirmed the identities of products **10** and **13** (**Figure S8**). From these data, it can be postulated that the *Cγ*-hydroxylated Arg (**10**) is the actual product of MilM, consistent with the efficient biosynthetic incorporation of this hydroxylated L-Arg residue into the mildiomycin structure^23^ (**Figure 1**), while the oxo product (**13**) observed under the *in vitro* assay conditions is possibly an off-pathway side product and does not probably have any biosynthetic significance in this pathway. Collectively, these biochemical and spectroscopic data demonstrate that MilM functions as a PLP-dependent oxidase cum hydroxylase, rather than an ATase, corroborating the recent findings.^28^

### Mechanistic Investigation of the MilM-Catalyzed Oxidation cum Hydroxylation

#### Proposed mechanism

A limited subset of PLP-dependent enzymes uses molecular oxygen as a co-substrate during catalysis.^69^ Based on MilM’s similarity with MppP^32^ from non-proteinogenic amino acid L-enduracididine biosynthesis (37% identity with MilM) and RohP^35^ from azomycin biosynthesis (60% identity with MilM), a mechanism for PLP-dependent oxidase MilM is proposed (**Figure 3**). This proceeds through a multi-step process involving radical intermediates and sequential oxidation, with two distinct quinonoid intermediates playing key roles in the mechanism. We propose that initially MilM forms an internal aldimine **17**, with PLP (**16**) likely through Lys232 (from our sequence and mutational analysis, **Figures S4** and **S5**), which is further converted to external aldimine (**18**) in presence of the substrate (L-Arg, **9**). Deprotonation of this complex at the Cα position (possibly using free Lys232 as the base) generates the first quinonoid intermediate (Q1, **19**), a highly stabilized, electron-rich species. Upon exposure to molecular oxygen, electron rich intermediate Q1 (**19**) possibly participates in single-electron transfer to O_2_, which has a biradical triplet ground state, thus generating the superoxide radical anion intermediate and a substrate-centered radical (**20**). This reactive superoxide radical anion may abstract a hydrogen atom from the β-carbon,^72^ forming an α, β-unsaturated imine (**21**) and a hydroperoxide anion (HO_2_^-^). The steps involving the formation of superoxide radical anion and hydrogen abstraction at the Cβ position (**Figure 3**) could potentially occur through alternative pathways as well, such as the involvement of a hydroperoxy-PLP adduct at the C4’ position.^28,35^ If the active site Lys232 attacks the PLP’s C4’ at this point, it would result in the formation of the proposed intermediate (**22**) and the generation of one equivalent of H_2_O_2_. Intermediate **22**, upon hydrolysis, would release NH_3_ and generate the oxo-product (**13**).

**Figure 3:**
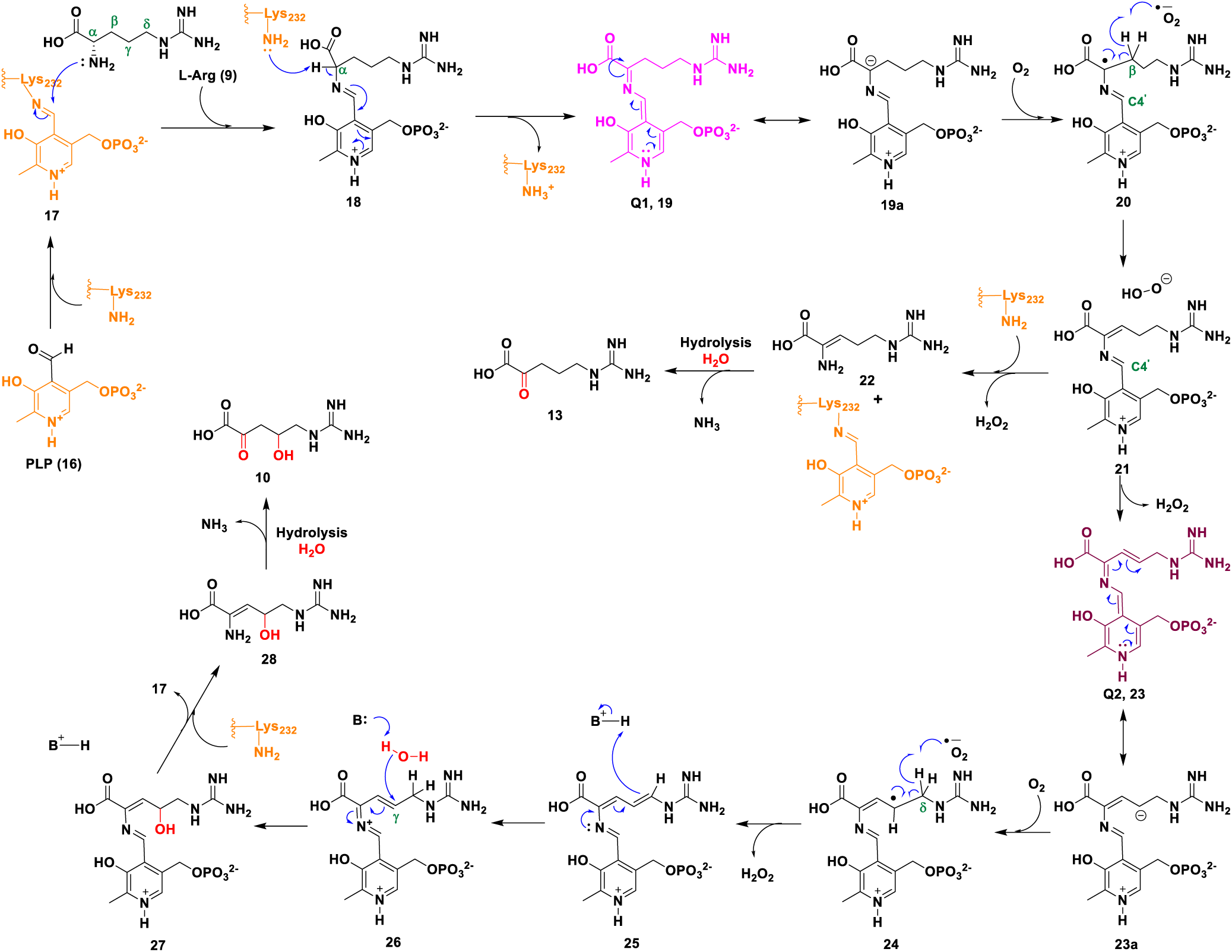
Proposed mechanism for the PLP-dependent oxidase cum hydroxylase, MilM. MilM catalyzes the PLP-dependent oxidation of L-Arg (**9**) through sequential steps involving two quinonoid intermediates (Q1 and Q2). Reaction with molecular oxygen generates possible radical intermediates, leading either to oxo-product (**13**) or to further hydroxylation. Based on our sequence alignment analysis, we hypothesize a key active site base (His31) mediates the regiospecific hydration of the didehydro intermediate (**25**), yielding the hydroxylated product 5-guanidino-4-hydroxy-2-oxovaleric acid (**10**). Lys232 cycles between forming the internal and external aldimine in the catalytic cycle of MilM, thus regenerating the essential cofactor.

In case Lys232 does not attack **21** prematurely, the elimination of the first equivalent of H_2_O_2_ will allow the system to form a highly conjugated second quinonoid intermediate (Q2, **23**). This species then reacts with a second O_2_ molecule, generating superoxide radical anion and a second radical intermediate (**24**). Abstraction of a hydrogen atom from Cδ of **24** could produce a diene intermediate (**25**), along with the release of another molecule of H_2_O_2_. The next stage involves hydration of the PLP-tethered didehydroarginine intermediate (**25**) for generating the hydroxylated product. First, we propose that **25** is transformed into intermediate **26** upon re-protonation at Cδ, possibly with the help of an active site protonated residue. Thereafter, the free base (likely residue is His31 from our sequence alignment, **Figure S4**) may activate a water molecule for nucleophilic attack at Cγ of **26**, installing the hydroxyl group and yielding the PLP-tethered hydroxylated intermediate (**27**). Completion of the catalytic cycle occurs when Lys232 attacks the PLP’s C4’, regenerating the internal aldimine (**17**) and releasing the hydroxylated enamine (**28**). The latter is subsequently hydrolyzed to yield the final product, 5-guanidino-4-hydroxy-2-oxovaleric acid (**10**), with concomitant release of ammonia by-product. Here, we report several biochemical experiments on MilM to support our mechanistic proposal.

#### UV-Vis spectroscopic analysis confirms formation of the proposed quinonoid intermediates

As discussed earlier, Lys232 is implicated as an important catalytic residue of MilM from our sequence and mutational analysis. To validate the functional role of Lys232 in internal aldimine formation with PLP, we examined the activity of the MilM-K232A variant using L-Arg as substrate. LC-MS analysis of the reaction mixture showed only unreacted L-Arg, with no detectable product, indicating complete loss of enzymatic activity in the K232A variant (**Figure S5C**). This result confirms that Lys232 is the primary catalytic Lys required to form the internal aldimine with PLP and to correctly position the cofactor for subsequent conversion to the external aldimine with L-Arg, thereby enabling progression of the catalytic cycle. Additionally, as expected, the MilM-K232A-K240A double variant was also found to be catalytically inactive (**Figure S5D**).

To further investigate the catalytic mechanism of MilM, we performed time-course UV-Vis spectroscopic analysis of the MilM reaction with L-Arg as substrate (**Figure S9**). Upon formation of the external aldimine (**18**), the absorption maximum corresponding to the internal aldimine (**17**, 420 nm) exhibited a slight red shift to 424 nm. Over time, two additional peaks characteristic of quinonoid intermediates appeared: Q1 (**19**) at 516 nm (pink in color) and Q2 at 565 nm. Notably, only Q1 accumulated over time, whereas Q2 (**22**) disappeared. This accumulation of Q1 suggests that molecular oxygen is required to drive the reaction forward, consistent with its proposed role in converting Q1 to advanced reaction intermediates (**Figure 3**). Indeed, vigorous shaking of the reaction mixture, which enhances oxygen dissolution, resulted in the rapid disappearance of the Q1 peak (along with the disappearance of the associated pink color), confirming its rapid reaction with O_2_. Interestingly, Q1 reappeared after a short interval, indicating the regeneration of the catalytic cycle. Similar trend was observed by reported PLP-dependent oxidases,^31,33^ as well as the recent MilM enzyme studies.^28^ To further validate the requirement of oxygen, MilM reaction was carried out under strictly anaerobic conditions inside a glove box. Under these conditions, the reaction mixture developed a pink color that persisted even after vigorous shaking. UV-Vis spectroscopic analysis of this reaction mixture showed characteristic Q1 absorption band (**Figure S10A**). Thereafter, MilM was removed by ultrafiltration, and the reaction mixture was analyzed by LC-MS which showed only unreacted L-Arg (*m/z* 175.1182) and no product-related masses were observed (**Figure S10B**). These results clearly demonstrate that O_2_ is indispensable for reaction progression beyond the Q1 intermediate, thus establishing its critical role in the MilM catalytic cycle.

#### MilM reaction is inhibited in the presence of SOD

In our proposed catalytic mechanism for MilM (**Figure 3**), quinonoid intermediates (Q1/Q2) are postulated to undergo single-electron transfer reactions with molecular oxygen to generate superoxide radical anion, which is likely to initiate hydrogen atom abstraction from the substrate to continue subsequent steps of oxidation. A similar mechanism of single-electron transfer was proposed for MppP,^34^ however, no evidence for the intermediacy of the superoxide radical anion has been reported thus far. To test this proposal, MilM reactions were performed in the presence of increasing concentrations of SOD, followed by LC-MS analysis of the reaction mixtures to evaluate the effect of SOD on product formation (**Figure S11A**). SOD is known to quench the reactive superoxide radical anion, converting it to molecular oxygen and hydrogen peroxide (**Figure S11B**).^72,73^ The effectiveness of SOD to quench superoxide radical anion generated during the MilM-catalyzed reaction (**Figure 3**) was further validated by quantifying the residual L-Arg in the MilM assay mixtures. Unreacted L-Arg was derivatized with DNS-Cl, and its concentration was determined using a standard calibration curve prepared with DNS-Arg (equivalent to L-Arg concentration, **Figure S12**). Interestingly, our experiments illustrated that increasing SOD levels led to a marked decrease in the product formation and a concomitant accumulation of the unreacted substrate, L-Arg in MilM reaction mixtures (**Figure 4A** and **Table S2**). We could completely quench the MilM reaction at a higher concentration of SOD (10 mg/ml, **Figure S11A**). This is likely due to the sequestering of reactive superoxide radical anion by SOD, thus preventing it from performing the critical hydrogen abstraction step in MilM catalysis (**Figure 3**). Hence, our results strongly support the possible involvement of the superoxide radical anion as a key intermediate in MilM catalysis, thereby linking the observed O_2_ dependence with the mechanistic role of reactive oxygen species in driving this remarkable transformation.

**Figure 4:**
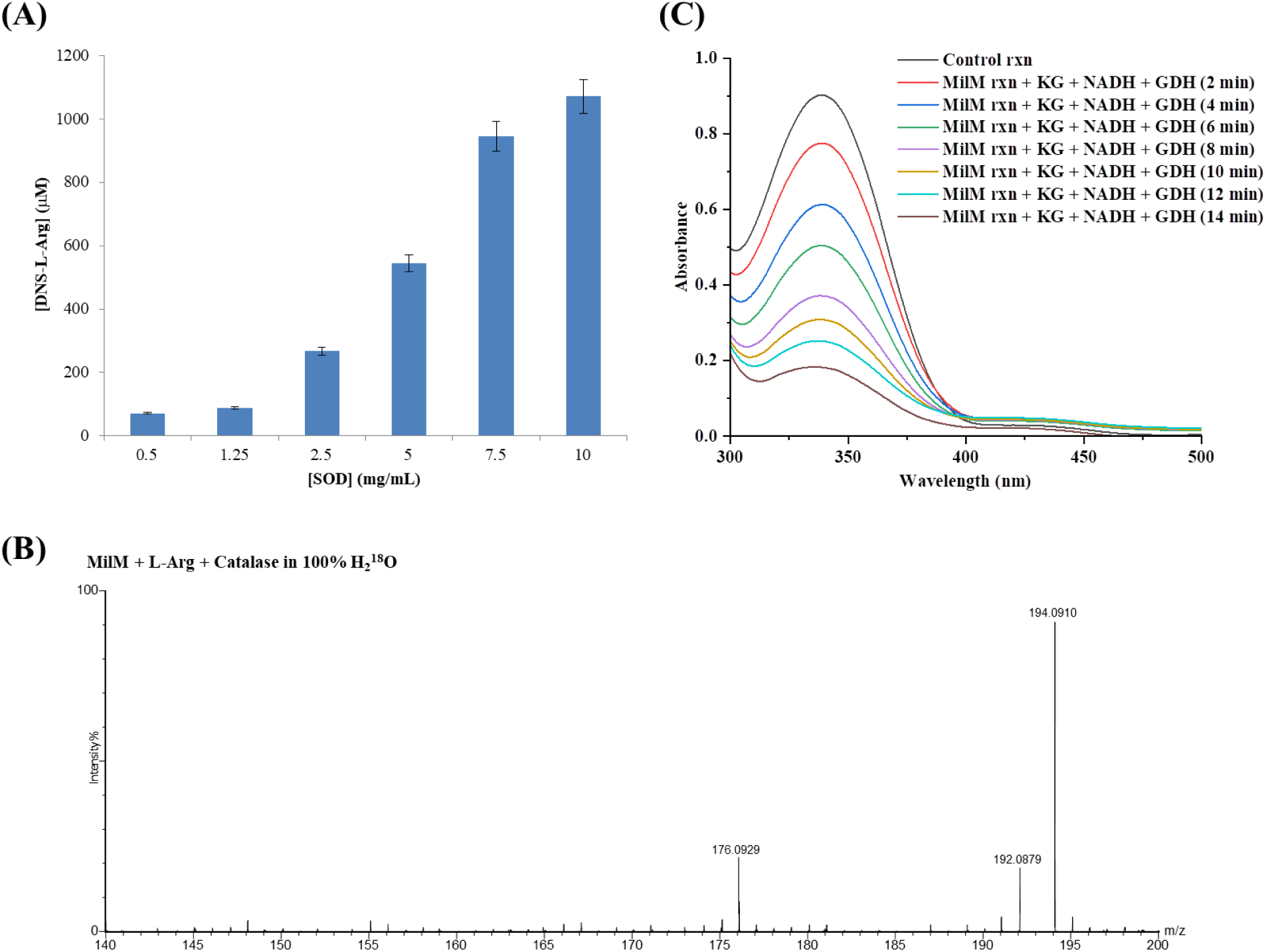
Mechanistic investigation of the MilM catalyzed reaction. **(A)** Quantification of residual L-Arg (as DNS-Arg derivative) in MilM assays containing increasing concentrations of SOD, showing a concentration-dependent suppression of MilM-catalyzed L-Arg oxidation due to possible quenching of the superoxide radical anion by SOD. DNS-Arg levels were determined from a standard calibration curve (**Figure S12C**). **(B)** LC-MS spectrum of the MilM reaction with L-Arg and catalase in 100% H_2_^18^O, confirming incorporation of solvent-derived ^18^O into the products, **10** (*m/z* 194.0910, calc. 194.0907) and **13** (*m/z* 176.0929, calc. 176.0929). **(C)** UV-Vis spectral changes for NADH at 340 nm observed over time in MilM reactions supplemented with α-KG, NADH, and GDH, indicating NH_3_ is a byproduct in MilM catalyzed reaction.

#### Water is the sole source of the oxo/hydroxyl group in MilM reaction products, not molecular oxygen

To determine the source of the oxygen atom incorporated into MilM reaction products (**10** and **13**, **Figure 2D**), isotope-labeling experiments were carried out using 100% H_2_^18^O as solvent in the presence of catalase. LC-MS analysis of the MilM reaction mixture revealed a +4 Da shift for product **10** (*m/z* 194.0910) and a +2 Da shift for product **13** (*m/z* 176.0929) (**Figure 4B**). These results confirm that the oxygen atom in products **10** and **13** in MilM reaction is derived from water, rather than from molecular O_2_, as expected from our proposed mechanism (**Figure 3**). This experiment further establishes that MilM functions as an oxidase/hydroxylase rather than an oxygenase^28^ (an oxygenase incorporates oxygen atom from molecular O_2_ into a reaction product),^74^ which is in line with observations made for other PLP-dependent oxidases MppP and RohP.^28,34,35^

#### Confirmation of His31 as the catalytic base

To investigate the proposed catalytic role of a general base for water activation needed for the hydroxylation step, we evaluated the pH dependence of MilM activity. As per our proposal, for hydroxylation, at first the general base must be in a protonated state to facilitate protonation of the intermediate **25** (at Cδ, **Figure 3**) followed by deprotonation of water by the free base for attack at Cγ of the intermediate **26** (**Figure 3**). To elucidate the role of pH, we assayed the MilM reaction over a pH range of 5.5 - 9.0. LC-MS analysis of the reaction mixtures revealed a decrease in formation of hydroxylated product (**10**) at higher pH (**Figure S13**). This trend indicates that the protonated form of the catalytic base is required for protonation of Cδ prior to water activation by the deprotonated base for a productive turnover, which is consistent with our proposal (**Figure 3**). Furthermore, sequence alignment of MilM with homologous enzymes (**Figure S4**), together with previously reported mutational data for the RohP-H34A variant,^35^ supports the assignment of His31 in MilM as the key residue fulfilling this catalytic function. To probe its function in MilM catalysis, H31A variant was created. The purified variant lacked the characteristic yellow color of the PLP cofactor and was also found to be catalytically inactive (**Figure S14A**). When MilM-H31A was reconstituted with external PLP during purification and tested for activity, the reaction mixture detected only a low level (∼10%) of the oxo product (**13**), with no formation of the hydroxylated product, **10** (**Figure S14B**). This behavior contrasts with RohP-H34A variant, which retains full activity for the oxo product formation.^35^ These data suggest that His31 in MilM is required not only for water activation in the hydroxylation step but also for possible stabilization of the L-Arg-PLP complex in the active site. In sum, from our studies it is evident that His31 is a key residue for binding and catalysis, however, additional protein crystallographic investigation will be needed to fully delineate the role of His31 in MilM.

#### Hydrogen peroxide and ammonia are released as by-products

The formation of H_2_O_2_ as a by-product in the MilM-catalyzed reaction (**Figure 3**) was validated using both LC-MS analysis in the presence of catalase and ABTS-HRP assay. As already discussed above, the LC-MS analysis of MilM assay performed in the presence of catalase, showed the disappearance of the non-enzymatic peaks at *m/z* 146.0923 (**14**) and *m/z* at 162.0874 (**15**). This indicates that H_2_O_2_ generated during the catalytic cycle reacts non-enzymatically with products (**10** and **13**) to produce off-pathway compounds **15** and **14**, respectively (**Figure S7**). To further confirm the generation of H_2_O_2_ as a by-product in the MilM reaction, ABTS-HRP coupled assay was performed. H_2_O_2_ generated during the enzymatic reaction oxidized ABTS in the presence of HRP to produce a bluish-colored ABTS•⁺ radical, having a strong absorbance at 418 nm (**Figures S15A** and **S15B**). Together, these results establish that MilM operates as an oxidase, utilizing molecular oxygen as an electron acceptor and producing H_2_O_2_ as a by-product during the oxidative conversion of L-Arg.

Finally, the release of ammonia as a by-product was confirmed using a coupled reaction of MilM with GDH in the presence of α-KG and NADH. We postulate that GDH can utilize NH_3_ released in the MilM reaction to convert α-KG to glutamate, with concomitant oxidation of NADH to NAD^+^, a transformation that can be monitored spectrophotometrically by observing the NADH absorbance change at 340 nm.^34,75^ As anticipated, UV-Vis spectroscopic analysis of this coupled reaction resulted in the time-dependent decrease of absorbance at 340 nm, consistent with NADH oxidation (**Figure 4C**). A control reaction without MilM did not show this change in absorbance at 340 nm. This indicates that NH_3_, released during the MilM reaction, is utilized in the GDH-catalyzed conversion of α-KG to L-Glu (**Figure S15C**), which in agreement with our mechanistic proposal (**Figure 3**).

### Structural Modeling Suggests MilM Forms a Functional Dimer

Since our native-PAGE and SEC analyses demonstrated that MilM predominantly exists as a dimer in solution, we embarked on modeling MilM as a dimer using AlphaFold-Multimer v3.^44,45^ Indeed, the dimeric model was predicted with a high confidence score (plDDT > 90) (**Figure 5A**). Next, we analyzed the dimer interface by the PISA server (**Table S3**).^47,48^ To understand the structural basis of MilM dimerization, the AlphaFold-dimer model of MilM was analyzed. This analysis revealed a stable dimeric assembly characterized by a well-defined interface between the two monomer chains A and B (**Figure 5B**) of the dimer. The PISA server estimated a solvation free energy gain ( Δ G) of -17.3 kcal/mol upon dimer formation, indicating that the association is thermodynamically favorable. The buried surface area at the interface was approximately ∼20409 Å^2^, consistent with stable protein-protein interactions observed in functional dimers (**Figure 5B**). Key interacting residues in the dimer interface are shown in **Table S3**, which undergo a significant reduction in solvent-accessible surface area upon dimerization.

**Figure 5:**
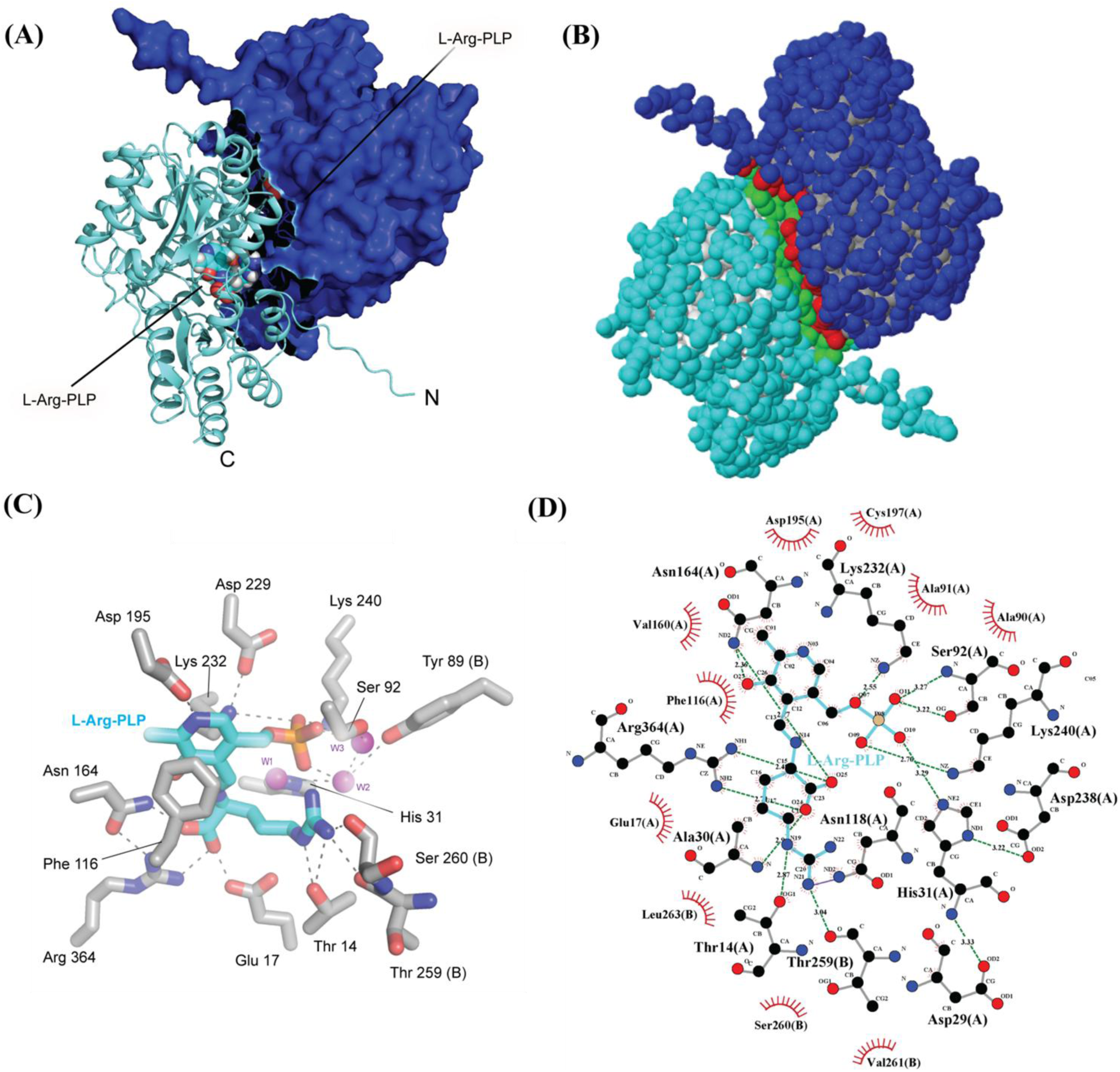
Alphafold model structure of MilM in complex with L-Arg-PLP. **(A)** Alphafold structure of MilM in dimer form bound with substrate L-Arg and cofactor PLP (L-Arg-PLP). Chain A is shown as cyan in the cartoon, and chain B as blue in the surface representation. **(B)** Dimer interface obtained by PISA server shows the interacting residues at the dimer interface highlighted in green for chain A and red in chain B. **(C)** A close-up view of the L-Arg-PLP binding pocket of MilM. Active-site residues are shown as grey sticks, while L-Arg and PLP are depicted in cyan. Coordinating water molecules are shown as violet spheres. Hydrogen bonds are indicated by grey dashed lines. **(D)** LigPlot analysis of the MilM-L-Arg-PLP complex model showing residues (grey) from chains A and B involved in interactions with PLP and L-Arg (cyan). Hydrogen bonds are indicated by olive green dashed lines, while maroon semicircular arcs represent hydrophobic interactions.

To further delineate the specific inter-subunit contacts within the MilM dimer, the modeled structure was analyzed using DIMPLOT (LigPlot+).^49^ The DIMPLOT interaction diagram revealed a well-organized network of hydrogen bonds and hydrophobic interactions stabilizing the interface between the two protomers (**Figure S16**). Several intermolecular hydrogen bonds were observed, primarily involving side chains of polar and charged residues including Gln36, Asp238, Thr14, Gln15, Asp12, Gln237, Asn264, Ser92, Asn125, Ser266, Trp51, Glu55, Asn256, Thr259, Val261, Ser260 of chain A and Glu55, Asn264, Thr259, Asn256, Ser266, Trp51, Asp238, Val261, Ser260, Gln237, Gln36, Gln15, Asp12, Thr14, Ser92, Asn125 of chain B, respectively (left to right, **Figure S16** and **Table S3**). In addition, clusters of non-polar residues form tight hydrophobic contacts that substantially contribute to interface packing and stabilization, including extensive interactions among residues on helices α1, α2, α3, α6, α7, α13, and α14 of each monomer (**Figures S17A** and **S17B**). Consistent with our structural and biophysical data, sequence alignment of MilM homologs further supports the evolutionary conservation of its oligomeric architecture. BLAST analysis^76^ identified eleven closely related homologs from *Streptomyces* species and other actinomycetes, and their alignment revealed that the key residues forming the MilM dimer interface are highly conserved across all sequences examined (**Figure S18**). The conserved interface residues shown in yellow align consistently across homologs, underscoring the conservation of polar contacts essential for interface packing and stability. Such strong conservation suggests that dimerization is not unique to MilM but represents a common and functionally important feature within these subgroups of PLP-dependent oxidases. These findings further established the physiological relevance of the MilM dimer observed in our structural modeling analysis. Together, our results confirm that the dimeric assembly predicted by AlphaFold-Multimer is structurally stable and represents the biologically relevant form of MilM in solution, consistent with our native-PAGE and SEC data demonstrating dimerization.

### Structural modeling of MilM with L-Arg-PLP reveals a conserved PLP-binding pocket

To explore MilM’s catalytic mechanism and cofactor-binding environment, we modeled MilM in complex with L-Arg-PLP using superimposition on RohP (PDB ID: 6C3C) (**Figure S19A**).^35^ Superposition of MilM with RohP [root mean square deviation (RMSD) 0.48Å for all Cα atoms] revealed strong structural similarity and a well-conserved PLP-binding pocket, implying that MilM adopts the canonical PLP-binding motif and potentially shares the catalytic mechanism of these enzymes. Using the Q1-bound (or L-Arg-PLP-bound) RohP complex as a template, the L-Arg-PLP adduct, and catalytic water molecules were positioned into the MilM active site (**Figures 5C** and **S19B**). The resulting complex displayed a geometrically favorable orientation of the external aldimine, consistent with catalytically competent conformations observed in other PLP-dependent enzymes such as RohP.^35^

The MilM-L-Arg-PLP complex model revealed that the phosphate group of PLP could be stabilized by a conserved phosphate-binding pocket composed of Ser92, Asp195, and Lys240, which form a network of hydrogen bonds and electrostatic interactions (**Figures 5C** and **S19B**). Phe116 provides hydrophobic stacking, while Asn164 contributes additional stabilization through hydrogen bonding to the PLP ring system. The substrate L-Arg interacts extensively with key catalytic residues Thr14, Glu17, His31, Asn118 and Arg364, which suitably stabilize its α-amino and guanidinium groups for productive catalysis (**Figures 5C** and **5D**). The active site showed a critical role for His31, which is strategically positioned to activate a water molecule (W1, **Figure 5C**) for nucleophilic attack on the C4 (Cγ) carbon of L-Arg (**Figure 7A**), thereby facilitating the hydroxylation reaction (**Figure 3**). Sequence alignment shows that this histidine is strictly conserved among PLP-dependent arginine oxidases (**Figure S4**), underscoring its mechanistic importance. Consistent with this, mutagenesis of the corresponding residue in RohP (H34A) abolishes hydroxylation activity, yielding only the oxo-product.^35^ Akin to this, our biochemical studies of the H31A variant of MilM showed no formation of the hydroxylated product (**Figure S14**). Taken together, these observations led us to propose that His31 in MilM serves as the catalytic base, deprotonating water and directing the reaction toward hydroxylation. Apart from chain A, several residues essential for PLP and substrate stabilization are located on chain B of the MilM dimer such as the hydroxyl group of Tyr89, through water forms hydrogen bonds and Val261 and Leu263 form hydrophobic interaction with PLP, thereby reinforcing cofactor anchoring across the dimer interface (**Figures 5C** and **5D**). A similar inter-subunit contribution has been reported in MppP, where PLP engages in a water-mediated hydrogen-bonding with Tyr88 and the main-chain amide of Leu252 from chain B of the dimer.^34^ This finding emphasizes the importance of dimerization, in which residues from chain B form the phosphate-binding pocket, thereby stabilizing the cofactor and ensuring proper catalytic geometry. Likewise, the carbonyl groups of amide backbones of Thr259 and Ser260 from chain B form hydrogen bonds with the guanidinium group of L-Arg in chain A, anchoring the substrate within the catalytic pocket of MilM as per our structural modeling analysis (**Figures 5C** and **5D**). These interfacial contributions demonstrate that a fully functional active site of MilM can only assemble upon dimerization. In the monomeric state, the geometry of PLP binding and substrate positioning would be compromised, leading to reduced or abolished catalytic activity. This structural observation supports our experimental findings from native-PAGE and SEC data, and confirms MilM’s dimeric nature in solution.

### MD simulation confirms MilM dimer forms the stable enzyme-substrate complex

To further elucidate the importance of dimerization on protein stability and substrate binding, we conducted atomistic MD simulations of the L-Arg-PLP-bound MilM dimer, and the L-Arg-PLP-bound MilM monomer systems. Overall, we carried out three independent MD production simulations for both the dimer and the monomer systems, each 150 ns long, yielding cumulative production duration of 450 ns for each. The first 25 ns of each run was excluded for equilibration and the remaining 125 ns from each run, amounting to a total of 375 ns for both dimer and monomer systems, were used for visualization and analysis.

We first visualized the equilibrated 375 ns simulation trajectories in VMD,^77^ and discovered that both substrates in chain A and chain B of the dimer consistently occupied the active site pocket over the whole simulation duration, a condition not applicable to the monomer simulation (**Figure 6A**). In two of the three-monomer simulations, we noticed that the substrate L-Arg-PLP departed from the active site and entered the bulk aqueous phase (**Figure 6A**). To further support this finding, we carried out several analyses from the equilibrated MD simulation trajectories using the CPPTRAJ module of AMBER.^78^ We computed the substrate solvation, the root mean square fluctuation (RMSF) of the substrate and the protein backbone (**Figures 6B**, **6C**, and **6D** respectively), as well as the non-bonded interaction energy between the protein and the substrate (**Figure 6E**).

**Figure 6:**
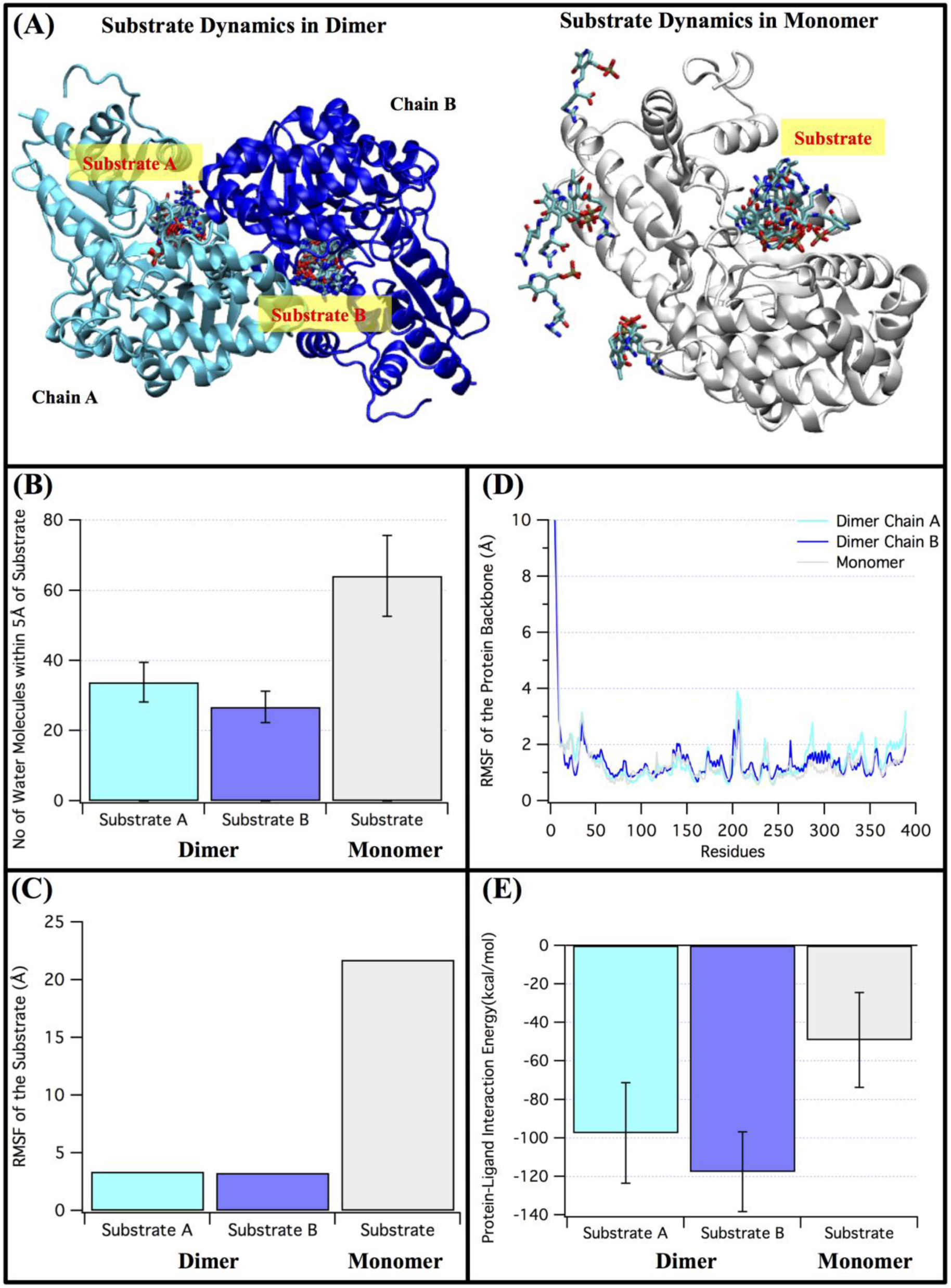
Comparison of substrate and protein dynamics between MilM dimer and monomer. **(A)** The time evolution of the substrate L-Arg-PLP’s movement with respect to the MilM active site. **(B)** The substrate solvation, quantified by the average number of water molecules within 5 Å of the substrate of interest. **(C)** and **(D)** represent the RMSF of the substrate, and the RMSF of the protein backbone atoms, respectively. **(E)** The average non-bonded interaction energy between the substrate and the protein. Here, 375 ns equilibrated trajectories of dimer and monomer simulations were utilized for comparative analysis. **(A)** was generated using VMD software package, whereas **(B)** to **(E)** were prepared with data analysis and visualization tool, IgorPro (https://www.wavemetrics.com/products/igorpro).

In order to assess the substrate solvation, we first counted the amount of water molecules within 5 Å of the substrate in each snapshot of the MD simulation, which was then time averaged over the entire 375 ns simulation duration. The standard deviation of the averaged water molecule counts was also calculated and presented as an error bar in **Figure 6B**. We independently quantified the water molecules for the substrate bound to chain A and the substrate bound to chain B in the dimer and compared those findings with the monomer simulations. Since, in monomer simulations, the substrate came out, it was expected to see higher ligand solvation in monomer than the dimer. Our analysis also demonstrated that the number of water molecules near the substrate was significantly higher, ∼64 water molecules on average, as opposed to substrate in chain A (∼34 water molecules) and chain B (∼27 water molecules) of the dimer (**Figure 6B**). This observation shows that the substrate in the monomer was more accessible to the water molecules, implying its presence in the bulk water during majority of the simulation, as illustrated in **Figure 6A**.

To further examine the mobility of the substrate and the protein, we computed the RMSF from the simulation trajectories. The higher RMSF usually indicates greater mobility of the substrate/residue of interest over the simulation time. To measure the RMSF, we first computed the time-averaged position of the substrate as well as each residue of the protein. The square root of the variance from this time-averaged position was then determined over the course of the 375 ns equilibrated simulation trajectories, physically corresponding to the average fluctuation (RMSF) from the time-averaged position. We measured the RMSF of substrate in chain A and chain B of the dimer as well as the substrate of the monomer (**Figure 6C**). We also measured the RMSF of the protein backbone atoms of the monomer as well as chain A and chain B of the dimer (**Figure 6D**). The substrate RMSF shows that the substrate exhibited significant mobility for the monomer (20 Å), in contrast to the substrates in chain and chain B of the dimer (∼3 Å), suggesting that the substrate was less stable in the active site of the monomer and predominantly moved freely in the bulk water (**Figures 6A** and **6B**). Conversely, the mobility of the substrates in chains A and B of the dimer was significantly constrained and predominantly localized within the hypothesized binding pocket, in line with the observations made in **Figure 6A**. To investigate whether the increased substrate mobility in the monomer resulted from protein instability, we compared the RMSF of the protein backbone of chains A and B of the dimer with that of the monomer (**Figure 6D**). In all instances, the RMSF of the protein backbone was notably comparable, with the initial nine residues of the N-terminal exhibiting significantly elevated RMSF (> 7 Å) in comparison to the remaining amino acids. The elevated RMSF of the N-terminal was anticipated due to its lack of secondary structure and its exposure to bulk water. In the remaining amino acid residues, the backbone RMSF typically stayed close to 2 Å, with the exception of the residues between 200 and 215. Overall, we observed no substantial divergence in the RMSF of the protein backbone between the monomer and dimer, indicating that the elevated RMSF of the substrate in the monomer simulation (**Figure 6B**) was evidently attributable to the absence of binding interactions between the substrate and the other monomer unit.

To further strengthen our hypothesis, we calculated the non-bonded interaction energies between the substrate and the protein in both the monomer and dimer configurations (**Figure 6E**). We also computed the non-bonded interaction energy on a per residue basis to identify the amino acids that are critical for substrate binding, as detailed in the Supporting Information (**Table S4**). Here, the non-bonded interaction energy is defined as the sum of electrostatic and Van der Waals interactions calculated between the substrate and the protein and averaged over 375 ns of equilibrated simulation trajectories. Our results clearly showed that the average interaction energy between the substrate and the MilM monomer was markedly less favorable than that of the substrate-dimer interaction, both for the substrate in chain A or substrate in chain B with the entire dimer (**Figure 6E**). The per residue interaction (**Table S4**) further revealed that many amino acids, such as Thr259, Ser260 from chain B of the dimer were interacting with the substrate in chain A, exhibiting a favorable energy lower than -1 kcal/mol. Similarly, these amino acids (Thr259, Ser260) from chain A of the dimer were found to interact with the substrate located in chain B. The absence of these interactions in the monomer simulation clearly caused the ligand to dissociate from the active site to the water, as highlighted in **Figure 6A**. Overall, our comparative analysis of the MilM dimer and monomer simulations revealed that the dimer configuration though does not contribute substantially to the protein stability (**Figure 6D**), it is highly critical for substrate binding. Hence, our 3D structural modeling and MD simulation studies clearly establish that dimerization is indispensable for stabilizing the enzyme-substrate-cofactor binding pocket for effective catalysis in MilM, a key feature that was not previously addressed for this class of PLP-dependent oxidases.

### Structure-guided mutagenesis reveals critical active side residues for MilM

In order to validate the proposed PLP binding residues of MilM as per our computational modeling (**Figure 5C**), we created Ala variants these residues (S92A, F116A, N164A, D195A, and K240A) using SDM, followed by their over-expression and purification. The activities of these variants were evaluated under similar conditions to that of MilM-WT (in the presence of catalase) using LC-MS analysis (**Figure S20**). None of these variants were found to be catalytically active, suggesting their importance in binding or activating PLP for catalysis, consistent with our modeling (**Figure 5C**) and MD simulation analysis (**Table S4**), where the per residue interaction with the substrate (L-Arg-PLP) exhibited a favorable energy lower than -1 kcal/mol.

Similarly, we also evaluated the activities of MilM variants which are postulated to interact with substrate L-Arg from our modeling studies: T14A, E17A, N118A and R364A (**Figure S21**). From our LC-MS analysis, it was found that R364A showed no activity at all while T14A, E17A and N118A showed formation of only the oxo product (**13**). No hydroxylated product (**10**) was observed in case of these variants, indicating that these residues are essential for the formation of the hydroxylated product (**10**). As per our model structure, Glu17 interacts with the carboxylic group of L-Arg (**Figure 7B**) and is critical for correctly positioning L-Arg in the active site for nucleophilic attack of a water molecule activated by His31 (W1, **Figure 7A**) at C4 (or Cγ) atom of L-Arg, thus enabling hydroxylation. MD simulation also revealed that the interaction of His31 with L-Arg-PLP was less than -2 kcal/mol (energetically favorable) as shown in **Table S4**, indicating its key role in cofactor/substrate binding and catalysis (**Figure S14**). In addition, Thr14 and Asn118 may hold the L-Arg guanidine group (tail) in position for an efficient hydroxylation reaction. In MilM variants where these residues were substituted with Ala, L-Arg still possibly binds in the active site, albeit loosely, hence allowing the formation of only the oxo product (**13**).

**Figure 7:**
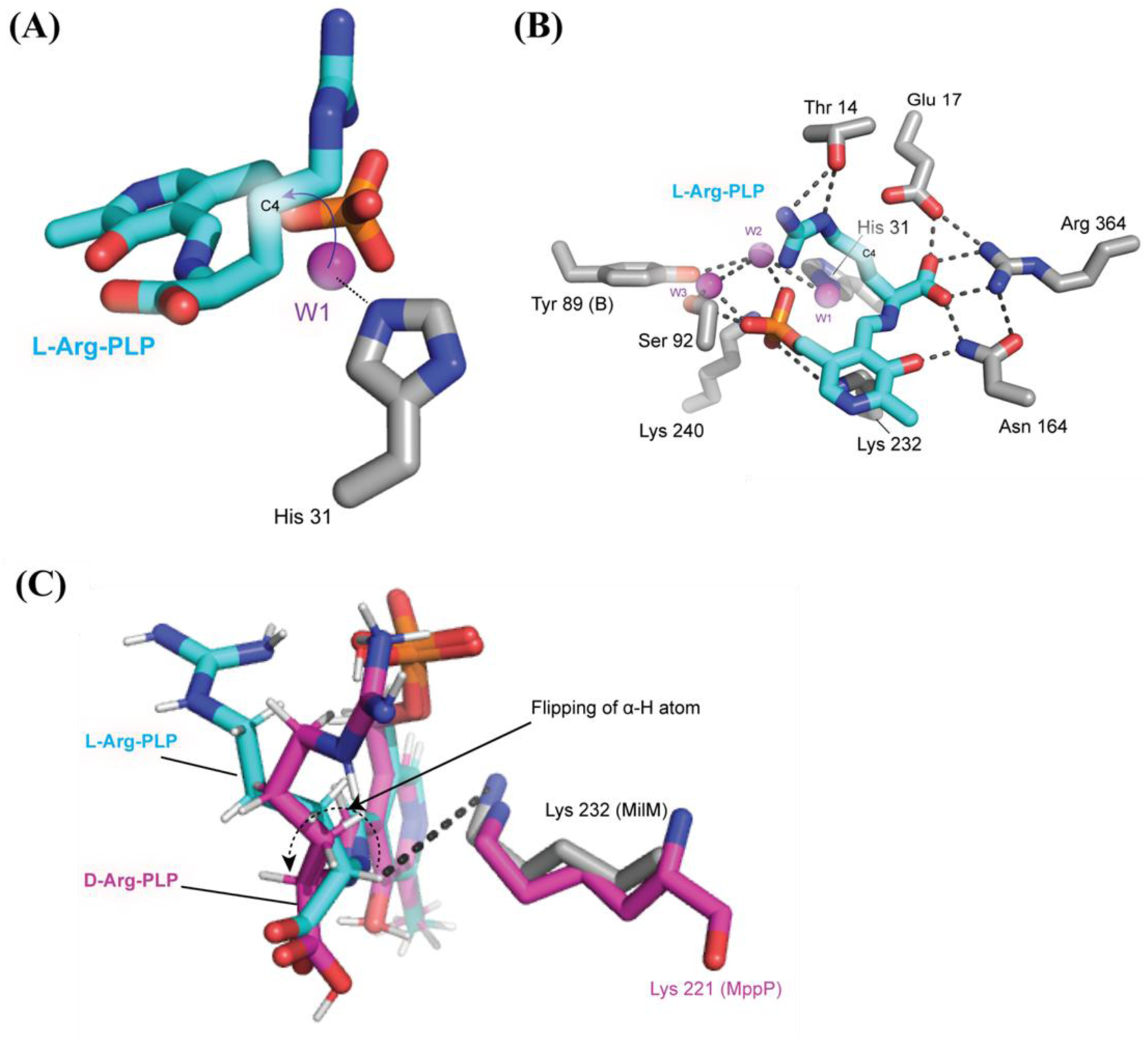
Structural model of the MilM active site showing crucial determinants of substrate positioning required for *Cγ*-hydroxylation. **(A)** Positioning of His31 relative to a modeled water molecule (W1, at a distance of ∼2.4 Å), based on the RohP quinonoid I structure, facilitating the nucleophilic attack of W1 at C4 (Cγ) of L-Arg. **(B)** In the MilM-L-Arg-PLP complex, Glu17 stabilizes Arg364 and Asn164, thus forming a network of hydrogen bonds with the carboxyl group of L-Arg which suitably positions the latter for nucleophilic attack of W1 at C4 during hydroxylation. **(C)** Superposition of MilM-bound L-Arg (cyan) and MppP-bound D-Arg (PDB ID: 5DJ3; purple) reveals outward flipping of the α-H in the D-Arg complex, leading to misalignment relative to Lys232 for hydrogen abstraction, and formation of a stalled, non-productive external aldimine (Figure 3).

Our 3D model structure of MilM homodimer (**Figure 5C**) and sequence alignment also showed some of the conserved residues from the other monomer interacting with either PLP or L-Arg (**Figure S4**). Among these, it was found that Tyr89 (**Figures 5C** and **7B**) interacts with the phosphate group of PLP via a water molecule (W2), while amide backbone of Thr259 and Ser260 engages directly with the L-Arg substrate (**Figure 5C**). To evaluate the importance of these residues in coordinating the substrate/cofactor, we created Tyr89 to Ala variant and Thr259/Ser260 to Pro double variant of MilM. Both variants, Y89A and T259P-S260P, showed no product formation when reacted with L-Arg in LC-MS analysis performed under conditions similar to those of WT-MilM (**Figure S22**). MD simulation studies also showed interactions between these two residues from chain A and L-Arg-PLP in chain B, and vice versa, with energies less than -1 kcal/mol (**Table S4**). This indicates that these residues from one monomer are also important for stabilizing the PLP or L-Arg in the other monomer, highlighting the importance of the dimeric form of MilM for efficient substrate binding and thus catalysis.

### Steady-state kinetics analysis confirms the role of critical active side residues

To verify these findings and to gain further insights into the enzyme-substrate interactions, steady-state kinetic analyses were performed for MilM-WT and the variants MilM-T14A, MilM-E17A and MilM-N118A, using the ABTS-HRP assay.^35,43^ To calculate the kinetic parameters, a H_2_O_2_ calibration curve (**Figure S23A**) was generated (see Materials and Methods for details). The MilM-WT exhibited *k*_cat_/*K*_m_ of 4.8 × 10^3^ s^-1^M^-1^, *k*_cat_ of 3.69 ± 0.04 min^-1^ and *K*_m_ of 12.75 ± 2.09 µM (**Figure S23B** and **Table 1**), values which are consistent with previously reported kinetic parameters of PLP-dependent arginine oxidases,^34,35^ indicating conserved catalytic behavior.

**Table 1:** Steady-state kinetic parameters for MilM variants with L-Arg as the substrate.

| Variants | $k_{\text{cat}}$ ($\text{min}^{-1}$) | $K_{\text{m}}$ ($\mu\text{M}$) | $k_{\text{cat}}/K_{\text{m}}$ ($\text{s}^{-1}\text{M}^{-1}$) | Products |
| --- | --- | --- | --- | --- |
| MilM-WT | $3.70 \pm 0.04$ | $12.75 \pm 2.09$ | $4.83 \times 10^3$ | Hydroxy (10) & Oxo (13) |
| MilM-T14A | $0.13 \pm 0.01$ | $114.77 \pm 25.35$ | $0.49 \times 10^2$ | Oxo only (13) |
| MilM-E17A | $1.33 \pm 0.02$ | $7.17 \pm 0.69$ | $9.26 \times 10^3$ | Oxo only (13) |
| MilM-N118A | $3.47 \pm 0.17$ | $13.27 \pm 3.64$ | $4.36 \times 10^3$ | Oxo only (13) |

The steady state kinetic studies of the variants revealed distinct functional roles for Thr14 and Glu17. MilM-T14A variant displayed severely diminished activity; with ∼98-fold lower turnover compared to MilM-WT, and generated only the oxo product, **13** (**Figure S23C** and **Table 1**). In contrast, MilM-E17A variant showed enhanced activity, with ∼1.9-fold higher turnover of the oxo product than MilM-WT (**Figure S23D** and **Table 1**). These results parallel earlier studies on MppP variants (MppP-T12A and MppP-E15A)^34^, which demonstrated similar effects, underscoring the functional conservation across this class of enzymes. MilM-N118A variant, which produced only the oxo product (**13**) was found to exhibit ∼1.1-fold lower turnover when compared with MilM-WT (**Figure S23E** and **Table 1**), suggesting that its activity is almost similar to that of the WT but mutation of Asn118 to Ala abolishes the hydroxylation activity of the enzyme. LC-MS analysis of the MilM variants that showed the product (**13**) distribution also corroborated these kinetic results (**Figure S21**). Structural modeling further supported these observations, revealing that Thr14, Glu17, and Asn118 are positioned to anchor L-Arg in a catalytically competent orientation, ensuring C4 hydroxylation by a water molecule activated through His31 (**Figure 7A**). Disruption of these interactions either destabilizes substrate positioning (T14A) or alters active-site dynamics (E17A, N118A), leading to impaired catalysis. Furthermore, these residues help maintain the geometry required for Q1 oxidation and subsequent hydroxylation, providing a structural explanation for the kinetic and product-specific trends observed. Collectively, these data highlight the critical roles of Thr14, Glu17 and Asn118 in governing the hydroxylation efficiency of MilM (**Figure 3**).

### Stringent substrate specificity and stereochemical requirement of MilM

To understand the substrate promiscuity of MilM, enzymatic assays were performed with other amino acids such as D-Arg, L-Lys, L-Glu, L-Gln and L-Ala as potential substrates. LC-MS analysis revealed no hydroxylated or oxo-products were formed from these amino acid substrates, suggesting that MilM has a strict substrate preference towards only L-Arg. D-Arg (the enantiomer of L-Arg) also failed to undergo any transformation (**Figure S24**), underscoring the strict stereo-chemical requirement for L-Arg recognition. Structural studies on MppP demonstrated previously that although D-Arg is capable of forming an external aldimine with PLP, the orientation of its α-proton is directed toward bulk solvent rather than toward the catalytic Lys residue (Lys 221 in MppP), leaving the intermediate catalytically inert.^33^ Productive catalysis requires the α-proton of L-Arg to be precisely oriented toward the catalytic Lys (Lys232 in case of MilM) and the associated catalytic base network to enable subsequent oxidation (**Figure 3**).

Superimposition of the crystal structure of MppP-D-Arg complex^33^ on MilM-L-Arg model structure (RMSD 1.0 Å for all Cα atoms) shows that the α-proton of D-Arg is oriented away from Lys232, disrupting this geometry and resulting in a stalled, non-productive external aldimine complex (**Figures 7C** and **S25A**). Thus, the inactivity of D-Arg in MilM reflects a conserved stereo-chemical constraint across this enzyme family, where catalysis depends on finely tuned stereo-electronic alignment of L-Arg within the PLP pocket. Such strict selectivity is consistent with the proposed active-site architecture of MilM, suggesting that precise interactions within the active site are essential for substrate binding and catalysis. Deviations in side-chain length (L-Lys or L-Gln), charge distribution (L-Glu), or stereochemistry (D-Arg) likely disrupt these critical contacts, thereby preventing catalysis. Together, the biochemical assays and structural modeling insights provide a coherent explanation for the observed substrate specificity of MilM.

Furthermore, to predict the probable configuration of the hydroxyl group in the hydroxylated product (**10**) in the MilM reaction, the crystal structure of RohP complexed with 4-hydroxy-2-ketoarginine (PDB ID: 6C3A)^35^ was superimposed with the MilM-L-Arg-PLP structural model (**Figure S25B**). From our model, it was evident that the hydroxyl group of the product complexed in the RohP active site oriented towards the water molecule (W1) in the MilM-L-Arg-PLP complex, suggesting that the nucleophilic attack from W1 at C4 (or Cγ) of the L-Arg complexed intermediate in MilM catalytic cycle (**Figure 3**) possibly occurs from the same face of the double bond as that of RohP. Thus, based on the homology of MilM with RohP,^35^ the stereochemistry of the hydroxyl group in **10** is likely to be of *S*-configuration,^28^ but further crystal structure studies on MilM complexed with the hydroxylated product will be required to fully delineate this stereochemical assignment.

### MD simulation exhibits a lid opening and closing mechanism for MilM dimer protein

Our MD simulation analysis also raised an important question, particularly in conjunction with the substrate entry and the product exit mechanism. If the dimerization is critical for substrate binding, how the substrate entry will occur, especially considering that the substrate exit was spontaneous in monomer simulation as the active site was greatly solvent exposed? Conversely, the substrate may encounter less obstacles (thermodynamically advantageous) when accessing the active site if the protein exists in a monomeric form, as opposed to the dimer configuration, where both active sites are obscured by the opposing monomer. Consequently, it was essential to ascertain the likely routes for substrate entry/product exit in the dimer; for which, we conducted tunnel analysis of the dimer and monomer simulation trajectories utilizing the CAVER 3.0 software package.^79,80^ Caver employs a geometric criteria to discern accessible cavities in the protein and group them to ascertain the likely tunnels connecting the active site to the solvent-exposed protein surface. For our case, we used the following parameters to identify the tunnels: a probe radius of 1.8 Å, a shell depth of 4 Å, a shell radius of 3 Å, and a clustering threshold of 2 Å. Also, the starting point for tunnel analysis was defined as the protein’s active site, defined as the center of mass of the proposed L-Arg-PLP binding residues, including Thr14, Glu17, His31, Ser92, Asn118, Asp195, Lys232, Lys240, and Arg364. Here, the probe radius used was slightly larger than the standard water probe (1.4 Å) to discern tunnels pertinent solely to substrate entry and product exit. We performed this analysis independently for the dimer’s chain A and chain B, as well as the monomer. Subsequently, three top-scoring tunnels for both the dimer and the monomer were identified and reported, along with their tunnel characteristics (**Figure 8A**). We could clearly see that the monomer tunnels had an average length of ∼4 Å and a radius of ∼3 Å, whereas the dimer tunnels were significantly longer (>10 Å) and had a smaller radius of ∼2 Å. This observation clearly demonstrates that the substrate entry and product exit are more thermodynamically favorable in the monomer than in the dimer. Nonetheless, substrate entry and product exit are still possible in the dimer, albeit with a far larger energy penalty than in the monomer. We hypothesize that because substrate binding is more thermodynamically beneficial in dimer than in the monomer (**Figures 6A** and **6E**), it is likely to overcome the substrate’s initial negative tunnel entry.

**Figure 8:**
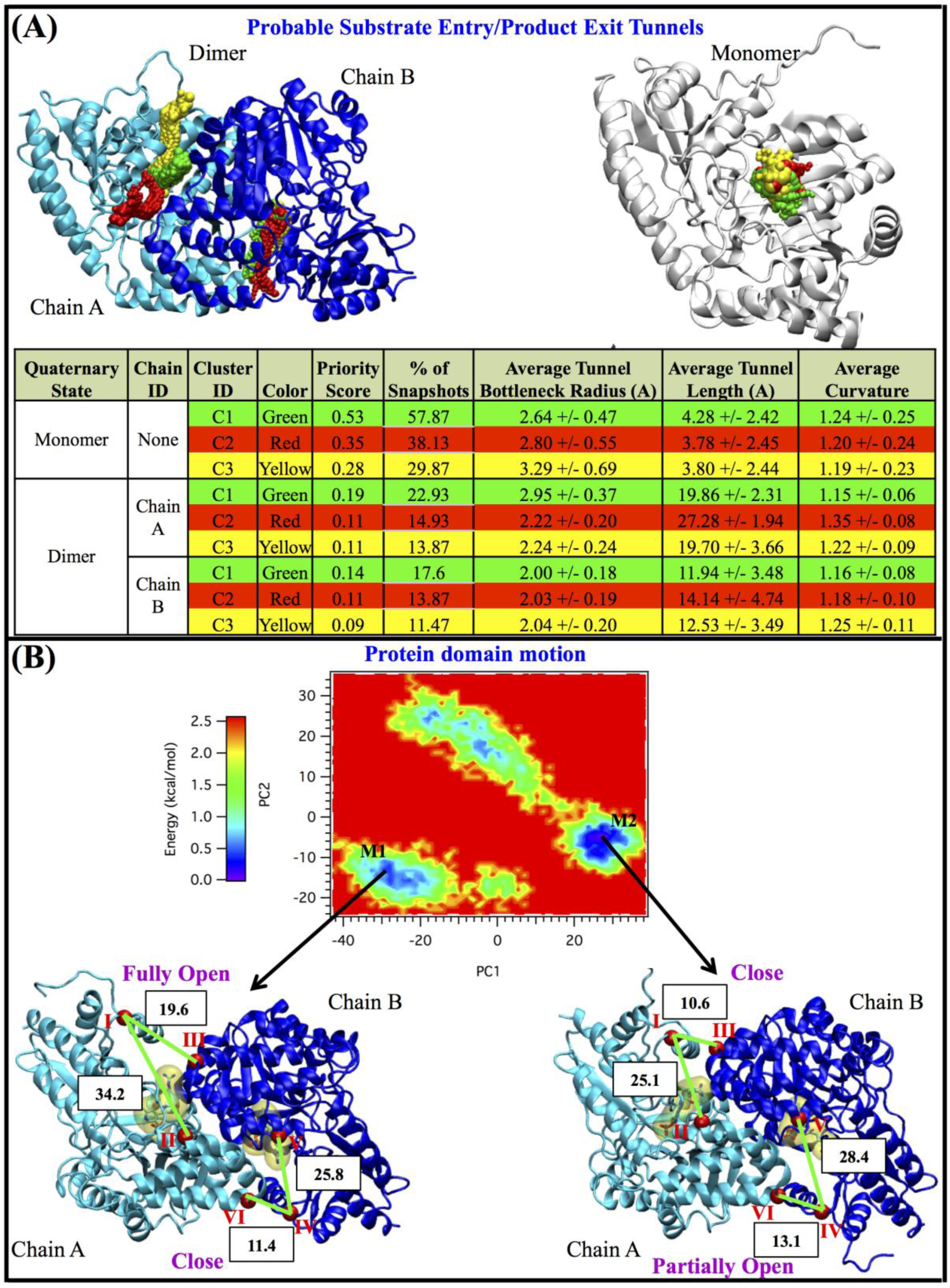
Substrate entry product exit mechanism in MilM, revealed through tunnel identification and principal component analysis. **(A)** Top scoring tunnels in both of the chains of the dimer and the monomer, denoted by green, red, and yellow beads. Here, the top scoring tunnels were determined according to the “Priority Score” values produced by the Caver software. The detailed characteristics of the tunnels including the tunnel average bottleneck radius, average tunnel length, tunnel curvature are also provided. **(B)** The free energy landscape obtained by projecting the equilibrated MD trajectory of dimer protein Cα onto principal components: PC1 and PC2. Representative dimer structures corresponding to two energy minima, M1 and M2, on the energy landscape are also provided. The distances (in Å unit) between Ser23 (I) of chain A and Gln41 (II) of chain A, Ser23 (I) of chain A and Arg58 (III) of chain B, Ser23 (IV) of chain B and Gln41 (V) of chain B, and Ser23 (IV) of chain B and Arg58 (VI) of chain A are also highlighted in black text with white rectangle background to emphasize the predominant motion associated with the lid opening and closing. Here, “Fully Open” lid conformation corresponds to a distance of ∼34 Å between Ser23 (I) and Gln41 (II) of chain A as well as a distance of ∼19 Å between Ser23 (I) of chain A and Arg58 (III) of chain B. “Partially Open” lid conformation corresponds to a distance of ∼28 Å between Ser23 (IV) and Gln41 (V) of chain B as well as a distance of ∼13 Å between Ser23 (IV) of chain B and Arg58 (VI) of chain A. “Close” conformation corresponds to a distance of ∼25 Å between Ser23 (I) and Gln41 (II) of chain A or Ser23 (IV) and Gln41 (V) of chain B as well as a distance of ∼11 Å between Ser23 (I) of chain A and Arg58 (III) of chain B or between Ser23 (IV) of chain B and Arg58 (VI) of chain A. Here, selection of these residues was based on visualization of the projected simulation trajectories in PC1 and PC2.

We also carried out the principal component analysis (PCA) to identify the major domain motions of the MilM dimer that may energetically favor the substrate entry/ product exit. For PCA, we first calculated the RMSD of the Cα atom positions of both of the chains of the dimer, excluding the first nine residues for which RMSF were significantly higher (**Figure 6D**). Subsequently, these Cα RMSD were projected into the resulting principal components (PCs). The first two principal components with the highest eigenvalues (PC1 and PC2) together accounted for over ∼67% of the total eigenvector variance, thus, expected to capture the major motions of the protein backbone (**Figure S26**). We independently visualized those projected trajectories corresponding to each of the two PCs (**Supplementary Movies 1** and **2**), and subsequently constructed a two-dimensional free energy landscape in PC1-PC2 coordinates, which revealed two distinct energy minima, M1 and M2 (**Figure 8B**). The representative MD snapshots corresponding to those minima were further compared based on the distances between Ser23 (I) and Gln41 (II) of chain A, Ser23 (I) of chain A and Arg58 (III) of chain B, as well as Ser23 (IV) and Gln41 (V) of chain B, and Ser23 (IV) and Arg58 (VI) of chain A (**Figure 8B**) to discern any significant conformational change that may arise during the MD simulations. The comparisons of these distances indeed demonstrated a substantial change, especially between Ser23 (I) and Gln41 (II) of chain A, as well as between Ser23 (I) of chain A and Arg58 (III) of chain B, where the distances greatly varied between the two minima, M1 and M2, with a change in distances of ∼9 Å, indicating the spontaneous opening (at M1 minima) and closing (at M2 minima) of the active site of chain A of the dimer. This observation also aligns with our tunnel analysis findings where we could identify three top scoring tunnels distinctly (**Figure 8A**), implying wider substrate entry/ product exit path for chain A active site. In contrast, the changes in distances were moderate in case of Ser23 (IV) and Gln41 (V) of chain B and Ser23 (IV) of chain B and Arg58 (VI) of chain A with a change in distances of ∼2.5 Å (**Figure 8B**). The tunnel analysis also showed that the top scoring tunnels were more clustered towards one another, representing comparatively narrower substrate entry/product exit path for chain B active site. Although we anticipated an opening (at M2 minima) and closing (at M1 minima) of the chain B active site comparable in magnitude to that of the chain A active site (2.5 Å vs 9 Å), we observed only a partial opening at M2 minima, which may be attributed to inadequate sampling despite conducting three independent simulations, each lasting 150 ns. Nevertheless, in both of these PCs, it was clearly evident that chain A and chain B of the dimer can approach each other and then recede, functioning like a lid that opens and closes to facilitate substrate entry and product release (**Figure 8B**; **Supplementary Movies 1** and **2**).

Interestingly, our PCA analysis revealed that when the active site of chain A is open, the active site of chain B remains closed at M1 minima; conversely, when the active site of chain B partially opens, the active site of chain A remains closed at M2 minima (**Figure 8B**). To determine if there is any correlated motion between chain A and chain B of the dimer that governs this alternating opening and closing of the active sites of both the chains, we calculated the cross-correlation matrix of the RMSD of protein Cα atoms, obtained from the MD trajectory (**Figure S27**). We observed a significant correlated motion between chain A and chain B of the dimer, particularly at the alpha helices near the dimer interface involving residues from 40 to 60 (helices α2-α4), and 265 to 280 (helix α14) of chain A exhibiting a positive correlated motion with residues from 40 to 65 (helices α2-α4) and 265 to 280 (helix α14) of chain B, respectively. Interestingly, these are the interface helices that demonstrated substantial domain movement during the opening and closing of the lid, as observed in our PCA analysis (**Figure 8B**; **Supplementary Movies 1** and **2**). We observed that when the interface helices containing Gln41 (II and V) and Arg58 (III and VI) from both chains A and B move away from Ser23 (I) of chain A, they approach Ser23 (IV) of chain B, leading to the opening of the active site of chain A and the closure of the active site of chain B. In a similar manner, if they are away from Ser23 (IV) of chain B, they are nearer to Ser23 (I) of chain A, resulting in the opening of the active site of chain B and the closure of the active site of chain A. Overall, our combined PCA and cross-correlation analysis indicate that the concerted movement of these interface helices governs lid opening and closing, thereby potentially lowering the energy barrier for substrate entry and product release.

### Lid Dynamics Rather than Global Oligomeric Transitions Governs Substrate Entry and Product Release in MilM Dimer

Since MilM exists as a dimer, for cofactor binding, substrate entry, and product release, generally we can postulate a monomer-dimer equilibrium mechanism, where MilM in monomer form first associates with another monomer unit using PLP cofactor and allows entry of the substrate, thus constituting the active dimer form for further catalysis. Thereafter, once the reaction is complete, the product is released from the active site on dissociation of the dimer. But, as evident from our 3D modeling, mutational, and MD simulation studies, the catalytic active site in MilM is positioned right at the dimer interface, and residues from both protomers coordinate PLP and L-Arg substrate, thereby indicating that the dimer represents the minimal functional unit in MilM. Dissociation into monomers would disrupt the proper positioning of the substrate-cofactor complex required for catalysis. Additionally, the extensive buried interfaces and energetic penalties associated with subunit separation render the monomer-dimer equilibrium hypothesis thermodynamically unfavorable and mechanistically unlikely. Rather, the preservation of the dimer throughout substrate entry and catalysis appears to be an evolutionarily conserved hallmark of PLP-dependent ATases or oxidases.^81^

PLP-dependent fold type I enzymes display dynamic open-closed conformational changes that are essential for their catalytic function.^82–85^ In the open state, the active site is accessible and structurally flexible, facilitating cofactor/substrate binding, whereas ligand binding induces movement of the small domain to close the active site, excluding bulk solvent.^82,83^ This closure stabilizes reactive intermediates and fine-tunes the active site geometry, thereby enhancing catalytic efficiency and substrate specificity.^82–84^ These enzymes are known to undergo substantial conformational rearrangements near the active site,^81^ particularly within flexible lid or loop regions that regulate solvent accessibility and catalytic alignment. Structural comparisons of apo and holo ATase forms have demonstrated that PLP binding induces ordering of previously disordered active-site elements, effectively completing the catalytic pocket and reducing nonspecific reactivity.^81^ This disorder-to-order transition has been directly visualized in ω-transaminases from *Chromobacterium violaceum*^83^ and *Vibrio fluvialis*,^86^ and a β-transaminase from *Mesorhizobium* sp.,^87,88^ where > 60 residues become structured only in the PLP-bound state. These observations also establish holo enzyme stabilization as a conserved feature of fold type I ATases.^81^

In line with these reports, a mechanistic model summarizing the substrate entry and product release cycle of the MilM dimer, derived from our MD simulations, tunnel analysis, and PCA/cross-correlation findings has been illustrated in **Figure 9**. The dimer consists of two protomers (chain A in cyan and chain B in blue), each harboring a PLP-dependent active site. Simulations revealed that MilM adopts a dynamic alternating lid mechanism, in which the interface helices undergo coordinated opening and closing motions that regulate access to each active site. We propose this occurs majorly in five stages in a cyclic manner as follows. Our mechanistic model starts with the holo state (**I**) with PLP bound as internal aldimines with Lys232 in the two active site pockets of the MilM dimer (chain A and chain B). We assume holo MilM likely adopts a closed conformation that occludes the active sites from solvent. Our structural model shows that PLP bound MilM is compact in nature (**Figure S28**). This architecture is likely to minimize nonspecific reactions (such as non-specific hydrolysis of the internal aldimine linkage) and preserves the reactive PLP cofactor in a protected environment while the enzyme awaits the substrate’s entry. However, it is possible that the enzyme may exist in a dynamic equilibrium between open and closed conformations in solution state as observed in our MD simulation experiments.

**Figure 9:**
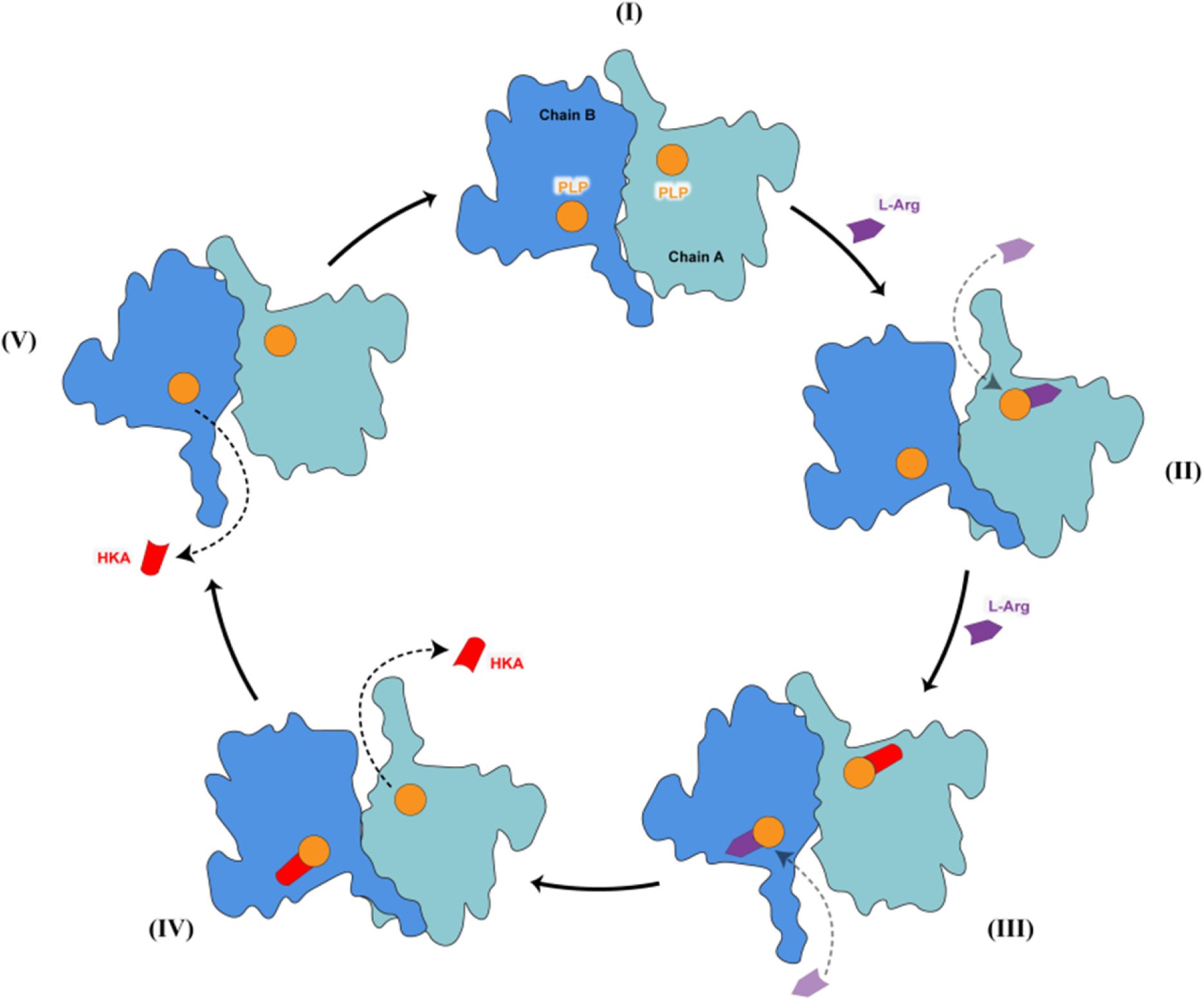
Proposed model of dynamic lid gating mechanism controlling substrate access and product release in the MilM dimer. (**I**) Closed-lid dimeric state with PLP cofactors bound in both chains A and B active site pockets of MilM. (**II**) Entry of L-Arg (substrate) in chain A from top open tunnel and external aldimine formation. (**III**) Another substrate entry in chain B from the bottom open tunnel forms the external aldimine in chain B. Meanwhile, in chain A closed state, catalysis proceeds in the active site, forming the hydroxylated product, **10**. (**IV**) Product formed in chain A exits via re-opening of the top tunnel with concomitant catalysis and product formation in chain B closed state. (**V**) Similarly, the product formed in chain B exits via re-opening of the bottom tunnel and the enzyme returns to the resting closed state (**I**). Chain A and B of MilM are represented in cyan and blue, respectively; PLP (cofactor) is represented as an orange circle; L-Arg (substrate) is represented as a purple arrow; and the hydroxylated product, **10** (also known as 4-hydroxy-2-ketoarginine, HKA) is represented as a red cylinder.

In the next step, displacement of the interface helices generates an open conformation (top tunnel), in which the active site of chain A becomes transiently accessible to the substrate. On the other hand, the chain B active site remains in the closed state and remains inaccessible to the substrate. In this state, L-Arg enters through the wider tunnels identified in our simulations and binds to PLP to form the external aldimine in chain A (**II**). Then, the top tunnel closes, and simultaneous opening occurs at the opposite end of the dimer (bottom tunnel) thus permitting a second L-Arg to access chain B active site pocket and form the external aldimine there as well (**III**). During this stage, chain A active site pocket (top tunnel) remains in the closed state, where catalytic progression enables product formation. Subsequently, the hydroxylated product, **10** (also known as 4-hydroxy-2-ketoarginine, HKA) is released from the active site of chain A through re-opening of the top tunnel in state **IV** (**Figure 9**). At the same time, the bottom tunnel closes, facilitating a similar catalytic event in chain B active site pocket. Finally, the product is released from the active site of chain B through re-opening of the bottom tunnel in state **V,** followed by reinstating the enzyme to the closed state (**I**). In these open γonformations (**IV** and **V**), the dimer may support simultaneous substrate entry and product exit, thereby maintaining catalytic throughput as long as the L-Arg substrate is available in the solution.

The global motions underlying these transitions align with the dominant PCs of the MD trajectory. This model is consistent with the PCA-derived free energy minima (M1 and M2), which captured mutually exclusive opening of the two active sites, and with the strong positive correlations observed between interface helices of chains A and B. Taken together, these findings support a concerted, see-saw-like lid mechanism for MilM that ensures only one active site is open at a time. This alternating-lid architecture provides a structural rationale for how the dimer maintains high substrate affinity while still permitting substrate entry and product exit through otherwise deeply buried active sites. However, further protein crystallographic or cryo-electron microscopic studies aimed at capturing MilM in multiple conformational states will be necessary to validate and refine this proposed dynamic model of substrate binding and catalysis in this class of PLP-dependent oxidases.

## CONCLUSIONS

In this study, we report detailed biochemical, structural modeling, mutational, and MD simulation-based experiments to elucidate the substrate binding and catalytic mechanism of MilM, a PLP-dependent oxidase from the mildiomycin biosynthetic pathway. MilM, contrary to the earlier annotation as an ATase, catalyzes the oxidation of L-Arg to produce 5-guanidino-4-hydroxy-2-oxovaleric acid (**10**) as the predominant product. We believe this class of enzymes has evolved mainly to perform *C*-hydroxylation chemistry rather than amino group transfer and the oxo product (**13**) is probably an artifact of the *in vitro* reaction and a biosynthetically insignificant side product. Active site architecture and conservation of certain critical residues possibly drive this change in reactivity. Catalytic His residue (His31) proposed to be responsible for water activation and hydroxylation step is conserved among hydroxylases and not in ATases,^35^ may be one of the key residues. However, the critical factors that are responsible for enabling this ATase enzyme fold to perform this fascinating PLP mediated C-H activation reaction remain to be investigated further.

Spectroscopic, biochemical, and isotope-labeling studies confirmed that MilM employs molecular oxygen and water in its catalytic cycle, which proceeds through two PLP-based quinonoid intermediates and a potential superoxide radical anion intermediate, critical for the C-H activation step. In addition, the hydroxyl group in the product was shown to be derived from water. These experiments establish MilM as the 3^rd^ reported member of this emerging class of PLP-dependent oxidase rather than a transaminase or an oxygenase. Steady-state kinetic analysis further revealed that MilM displays catalytic efficiency comparable to known L-Arg oxidases MppP^34^ and RohP,^35^ while mutational studies identified Thr14, Glu17, Asn118 and Arg364 as key residues for substrate positioning and hydroxylation chemistry. A central advance of this work is the demonstration that dimerization is essential for MilM function, a key feature which was not properly addressed earlier in PLP-dependent L-Arg oxidases. Our modeling analysis identified the critical residues which are involved in stabilizing the MilM dimer interface. Furthermore, SEC/native-PAGE experiments supported by 3D structure-based modeling analysis, MD simulation, and SDM, revealed that several residues required for PLP and substrate stabilization in chain A [including Tyr89 (B), Thr259 (B), and Ser260 (B)] are contributed by the opposing chain B. Specifically, the MD simulation revealed a spontaneous escape of the substrate from the monomer’s active site when the protein was not simulated as a dimer. These observations altogether clearly establish that a catalytically competent state of the enzyme is achieved only in the dimeric state. These interfacial interactions are indispensable for anchoring both the phosphate group of PLP and the guanidinium moiety of L-Arg, highlighting a unique structural dependence on dimerization for cofactor and substrate positioning. To the best of our knowledge, this is the first demonstration that dimerization is required for efficient substrate/cofactor coordination and catalysis in PLP-dependent L-Arg oxidases cum hydroxylases. Furthermore, MD simulation data supports a substrate-activated lid-gating model as the primary substrate entry and product exit mechanism in MilM, with the dimer maintained as a stable and essential catalytic unit throughout turnover. Together, our independent findings redefine the functional role of MilM as a *C*-hydroxylase in mildiomycin biosynthesis, as opposed to earlier prediction as an ATase, thus reinforcing the recent report by Edwards et al.,^28^ apart from providing clarity on its substrate specificity and mechanistic underpinnings. Beyond MilM, this work suggests that dimerization-dependent active site assembly may be a broader feature among PLP-dependent oxidases, a notion that merits further biochemical, theoretical, and structural investigations. Future crystallographic studies of MilM will be invaluable to elucidate the structural determinants of this dimerization-dependent mechanism fully, and to explore its implications for engineering such oxidases with tailored substrate scopes for prospective combinatorial biosynthesis and chemo-enzymatic applications.

## ASSOCIATED CONTENT

## Supporting Information

**The supporting information is available free of charge at** https://pubs.acs.org/doi

Includes additional experimental details, primers used for plasmid construction and site-directed mutagenesis, SDS-PAGE gels for proteins and mutants, sequence alignment for proteins, UV-Vis chromatograms, HPLC, LC-MS, and NMR data, Michaelis-Menten curves, computationally modeled structures, and MD simulation movies.

Supporting information document (One, file type, i.e., PDF)

MD simulation movies (Two, file type, i.e., MP4)

## Accession Codes

The UniProt and NCBI accession codes for MilM protein from *S. rimofaciens* are H9BDX2 and AFD20753, respectively.

## AUTHOR INFORMATION

## Authors

Complete contact information is available at: https://pubs.acs.org/

## Author Contributions

The manuscript was written through contributions of all authors. All authors have given approval to the final version of the manuscript.

## Notes

The authors declare no competing financial interest.

## ACKNOWLEDGMENT

This work was supported in part by the research grants from the Council of Scientific and Industrial Research, Govt. of India (02/0419/21/EMR-II to N.M.), Anusandhan National Research Foundation-Core Research Grant, Govt. of India (CRG-2022-004059 to N.M.), Indian Council of Medical Research-Extramural research grant (2021-15271 to N.M.), Ministry of Education’s Scheme for Transformational and Advanced Research in Sciences (STARS-2/2023-0657 to N.M.), and IIT Dharwad-Seed Research Grant (IITDH/R&D/6.81/SGNF/2023-24/169 to SJ). SJ also thanks IIT Dharwad High Performance Computing (HPC) facility for providing the computational hours. BND acknowledges the Ramalingaswami Re-entry Fellowship Grant (IITDH/R&D/BD/2025–26/269) from the Department of Biotechnology, Government of India.

## ABBREVIATIONS

PLP: pyridoxal-5’-phosphate
ATase: aminotransferase
SOD: superoxide dismutase
GDH: glutamate dehydrogenase
MD: molecular dynamics.

