## Supplementary information document for "Dimerization of MilM is essential for catalyzing the pyridoxal-5’-phosphate (PLP)-dependent Cγ-hydroxylation of L-arginine during mildiomycin biosynthesis"

**Table of Contents:**

Table S1: Oligonucleotide primers used in this study ............................................................S3

Table S2: Amount of the residual substrate L-Arg in the MilM reaction in the presence of different SOD concentrations using DNS-Cl derivatization assay ..........................................S4

Table S3: PISA server derived residues in dimer interface of MilM protein ..........................S5

Table S4: MilM Dimer and L-Arg-PLP interactions ..............................................................S7

Figure S1: Agarose gel electrophoresis analysis of isolated genomic DNA ...........................S15

Figure S2: SDS-PAGE analysis of the proteins used in this study ..........................................S16

Figure S3: Spectroscopic and chromatographic characterization of PLP cofactor in MilM ....S17

Figure S4: Sequence alignment of MilM with homologous enzymes.....................................S18

Figure S5: Spectroscopic and analytical characterization of MilM lysine mutants.................S19

Figure S6: DNS-Cl derivatization of L-Arg and analysis of MilM reaction mixtures .............S20

Figure S7: LCMS analysis of MilM-catalyzed reaction with L-Arg as substrate ...................S21

Figure S8: $^{1}$H-NMR analysis of the MilM assay .....................................................................S22

Figure S9: UV-Vis spectroscopic investigation of the MilM reaction showing the time-dependent formation of quinonoid intermediates during the MilM-catalyzed oxidation of L-Arg ..........................................................................................................................................S23

Figure S10: MilM assay with L-Arg under anaerobic conditions ...........................................S24

Figure S11: MilM reaction in presence of SOD ....................................................................S25

Figure S12: Quantification of residual L-Arg in the MilM reaction in the presence of SOD using DNS-Cl derivatization ..........................................................................................S26

Figure S13: pH-dependent product formation catalyzed by MilM..........................................S27

Figure S14: LCMS analysis of MilM-His31Ala variant...........................................................S28

Figure S15: Detection of $H_2O_2$ and $NH_3$ as by-products in MilM-catalyzed reaction ............S29

Figure S16: DimPlot analysis of residues stabilizing the MilM dimer interface......................S30

Figure S17: Topology of MilM...............................................................................................S31

Figure S18: Sequence alignment of MilM homologs showing conservation of residues at the dimer interface ........................................................................................................................S32

Figure S19: Structural comparison of MilM...........................................................................S35

Figure S20: LCMS analysis of the reactions of PLP-binding MilM variants with L-Arg substrate ...................................................................................................................................S36

Figure S21: LCMS analysis of the reactions of L-Arg binding MilM variants........................S37

Figure S22: LCMS analysis of the reactions of MilM variants from chain B binding with L-Arg-PLP in chain A .........................................................................................................................S38

Figure S23: Steady-state kinetic analysis of MilM-WT and its variants .................................S39

Figure S24: LCMS analysis of the MilM assay using D-Arg as the substrate .........................S40

Figure S25: Superimposition of MilM model structure with MppP and RohP crystal structures. ................................................................................................................................S41

Figure S26: The characteristics of the major principal components........................................S42

Figure S27: Residue cross-correlation map based on RMSD of protein Cα atoms.................S43

Figure S28: 3D model structure of holo MilM enzyme...........................................................S44

Supporting references ..............................................................................................................S45

**Table S1: Oligonucleotide primers used in this study.** For all sequences provided 5' to 3', "fwd" indicates a forward primer; "rev" indicates reverse primer; underlined are the BamHI and NotI restriction sites and lower-case letters indicate the *altered codon* for each variant (G. DNA = genomic DNA).

| Primer name | Oligonucleotide Sequence | Template Used |
|---|---|---|
| MilM_BamHI_fwd | AAAGGATCCATGTCCGACACTCTCGC | G. DNA |
| MilM_NotI_rev | AAAGCGGCCGC TCACGCATAGCGGGC | G. DNA |
| MilM_T14A_fwd | CCCTCGACCTGgccCAGCACGAGATAGCGGCCCTGCGCTCC | MilM-WT |
| MilM_T14A_rev | ATCTCGTGCTGggcCAGGTCGAGGGGACGGTTGTGCGCGAG | MilM-WT |
| MilM_E17A_fwd | TGACCCAGCACgcgATAGCGGCCCTGCGCTCCGAGCACAAT | MilM-WT |
| MilM_E17A_rev | AGGGCCGCTATcgcGTGCTGGGTCAGGTCGAGGGGACGGTT | MilM-WT |
| MilM_H31A_fwd | TCGCGGACGCGgccACGCACCAGTACCAGTCGCCGGCCCAG | MilM-WT |
| MilM_H31A_rev | TACTGGTGCGTggcCGCGTCCGCGAGATTGTGCTCGGAGCG | MilM-WT |
| MilM_Y89A_fwd | CGCTGCTCACCgccGCCGCCTCCATCTCCACGATGATCGCC | MilM-WT |
| MilM_Y89A_rev | ATGGAGGCGGCggcGGTGAGCAGCGTGCGGTCCAGGCCGAT | MilM-WT |
| MilM_S92A_fwd | CCTACGCCGCCgccATCTCCACGATGATCGCCGGGATGTTC | MilM-WT |
| MilM_S92A_rev | ATCGTGGAGATggcGGCGGCGTAGGTGAGCAGCGTGCGGTC | MilM-WT |
| MilM_F116A_fwd | TCGAGCCCTGCgccGACAACCTCCCCGACCTGCTCGTCAAT | MilM-WT |
| MilM_F116A_rev | GGGAGGTTGTCggcGCAGGGCTCGACCAGCGTCACCCGCGC | MilM-WT |
| MilM_N118A_fwd | CCTGCTTCGACgccCTCCCCGACCTGCTCGTCAATCTGGGC | MilM-WT |
| MilM_N118A_rev | AGGTCGGGGAGggcGTCGAAGCAGGGCTCGACCAGCGTCAC CCGC | MilM-WT |
| MilM_N164A_fwd | TCGACCCCAACgccCCGACTGGCCATAGCCTGTTCGCCGAC | MilM-WT |
| MilM_N164A_rev | TGGCCAGTCGGggcGTTGGGGTCGACGAGAAAAAGCGCCTC | MilM-WT |
| MilM_D195A_fwd | TCCTCGTCCTCgccCTGTGCTTCGCGGCCTTCGCCCTCGGC | MilM-WT |
| MilM_D195A_rev | GCGAAGCACAGggcGAGGACGAGGACCGTGCCGCGCTCGCG | MilM-WT |
| MilM_K232A_fwd | GGACACCGGCgcgACCTGGCCCGTCCAGGACGCCAAATGC | MilM-WT |
| MilM_K232A_rev | ACGGGCCAGGTcgcGCCGGTGTCCTCCATGGCGATGTAGGT | MilM-WT |
| MilM_K240A_fwd | TCCAGGACGCCgcaTGCGCCCTGCTCACCACCAGCGCCGAC | MilM-WT |
| MilM_K240A_rev | AGCAGGGCGCAtgcGGCGTCCTGGACGGGCCAGGTCTTGCC | MilM-WT |
| MilM_K232A_K240A_fwd | TCCAGGACGCCgcaTGCGCCCTGCTCACCACCAGCGCCGAC | MilM-K232A |
| MilM_K232A_K240A_rev | AGCAGGGCGCAtgcGGCGTCCTGGACGGGCCAGGT | MilM-K232A |
| MilM_T259P_S260P_fwd | ACAACCTCCACccccccGTCCTGCTGAACGTCTCGCCCTTC | MilM-WT |
| MilM_T259P_S260P _rev | TTCAGCAGGACggggggGTGGAGGTTGTACACGGCGGGGTA | MilM-WT |
| MilM_R364A_fwd | AGCGCTACGTCgcgGTGGCGCTGGCGCGTGATCCCGGGGAG | MilM-WT |
| MilM_R364A_rev | GCCAGCGCCACcgcGACGTAGCGCTCGCCGCGGCTCGGCTC | MilM-WT |

**Table S2: Amount of the residual substrate L-Arg in the MilM reaction in the presence of different SOD concentrations using DNS-Cl derivatization assay.**

| [L-Arg] (μM) | [SOD] (mg/mL) | [DNS-Arg] (μM) | Estimated % of remaining L-Arg substrate |
|---|---|---|---|
| 1000 | 0.50 | 72 | 7 |
| 1000 | 1.25 | 89 | 8 |
| 1000 | 2.50 | 268 | 25 |
| 1000 | 5.00 | 546 | 51 |
| 1000 | 7.50 | 947 | 88 |
| 1000 | 10.0 | 1000 | 100 |

**Table S3: PISA server derived residues in dimer interface of MilM protein.**
(Source: PISA Server, https://www.ebi.ac.uk/pdbe/pisa).[1]

| Chain A | | Distance [Å] | Chain B | |
|---|---|---|---|---|
| Residue No. | Amino Acid | | Residue No. | Amino Acid |
| 259 | THR [OG1] | 2.57 | 12 | ASP [OD2] |
| 237 | GLN [NE2] | 3.07 | 51 | TRP [O] |
| 36 | GLN [NE2] | 2.77 | 55 | GLU [OE1] |
| 104 | ARG [NH1] | 3.62 | 125 | ASN [OD1] |
| 266 | SER [OG] | 2.57 | 237 | GLN [O] |
| 55 | GLU [N] | 3.11 | 237 | GLN [OE1] |
| 264 | ASN [ND2] | 3.08 | 238 | ASP [OD2] |
| 15 | GLN [NE2] | 3.07 | 256 | ASN [OD1] |
| 118 | ASN [ND2] | 3.47 | 259 | THR [O] |
| 15 | GLN [NE2] | 3.38 | 259 | THR [OG1] |
| 125 | ASN [ND2] | 2.92 | 260 | SER [OG] |
| 92 | SER [OG] | 2.57 | 261 | VAL [O] |
| 12 | ASP [OD2] | 2.57 | 259 | THR [OG1] |
| 51 | TRP [O] | 3.07 | 237 | GLN [NE2] |
| 55 | GLU [OE1] | 2.77 | 36 | GLN [NE2] |
| 125 | ASN [OD1] | 3.84 | 104 | ARG [NH1] |
| 237 | GLN [O] | 2.58 | 266 | SER [OG] |
| 237 | GLN [OE1] | 3.10 | 55 | GLU [N] |
| 238 | ASP [OD2] | 3.05 | 264 | ASN [ND2] |
| 256 | ASN [OD1] | 3.12 | 15 | GLN [NE2] |
| 259 | THR [O] | 3.39 | 118 | ASN [ND2] |
| 259 | THR [OG1] | 3.40 | 15 | GLN [NE2] |
| 260 | SER [OG] | 2.93 | 125 | ASN [ND2] |
| 261 | VAL [O] | 2.56 | 92 | SER [OG] |

Standard atom nomenclature in Protein Data Bank (with reference to **Table S3**)

| Atom Code | Typical Residues | Description |
|---|---|---|
| N | All amino acids | Backbone nitrogen (amide N-H) |
| O | All amino acids | Carbonyl oxygen of peptide bond |
| OG | Serine | Side-chain hydroxyl oxygen |
| OG1 | Threonine | Hydroxyl oxygen on the side chain’s first branch |
| OD1 / OD2 | Aspartate, Asparagine | Side-chain carboxyl or amide oxygens |
| OE1 | Glutamate, Glutamine | Side-chain carboxyl or amide oxygens |
| NE2 | Histidine | Ring nitrogens in imidazole |
| ND2 | Asparagine | Delta-2 nitrogen atom |
| NH1 | Arginine | Terminal guanidinium group nitrogens |

**Table S4: MilM Dimer and L-Arg-PLP interactions.** The total non-bonded interaction energy (TNBIE) between the residues of chains A and B of the dimer, with the ligand (L-Arg-PLP) bound to chain A and B, computed from MD simulations. Here, the "TNBIE" value is the combination of electrostatic and van der Waals interactions, with more negative values indicating stronger interactions between the residue and the substrate, positive values denoting unfavorable interactions, and zero implying no non-bonded interactions.

**Interaction between residues of chain A and the L-Arg-PLP bound to chain A:**

| Residue No. | TNBIE (kcal/mol) | Residue No. | TNBIE (kcal/mol) | Residue No. | TNBIE (kcal/mol) | Residue No. | TNBIE (kcal/mol) | Residue No. | TNBIE (kcal/mol) |
|---|---|---|---|---|---|---|---|---|---|
| 1 | -0.01 | 79 | 0.00 | 157 | 0.00 | 235 | -0.02 | 313 | 0.00 |
| 2 | 0.00 | 80 | 0.00 | 158 | 0.28 | 236 | -0.10 | 314 | 0.00 |
| 3 | 0.00 | 81 | 0.00 | 159 | -0.56 | 237 | -0.04 | 315 | 0.00 |
| 4 | 0.00 | 82 | 0.00 | 160 | 0.09 | 238 | 2.11 | 316 | 0.08 |
| 5 | 0.00 | 83 | 0.00 | 161 | -0.34 | 239 | 0.00 | 317 | -0.38 |
| 6 | 0.00 | 84 | 0.00 | 162 | -0.37 | 240 | -7.09 | 318 | -0.16 |
| 7 | 0.00 | 85 | 0.00 | 163 | -0.04 | 241 | 0.05 | 319 | -0.16 |
| 8 | 0.00 | 86 | 0.00 | 164 | -3.95 | 242 | -0.04 | 320 | 0.00 |
| 9 | -0.01 | 87 | 0.00 | 165 | -0.12 | 243 | -0.17 | 321 | -0.02 |
| 10 | 0.00 | 88 | 0.00 | 166 | 0.05 | 244 | 0.05 | 322 | 0.00 |
| 11 | 0.04 | 89 | 0.05 | 167 | -0.22 | 245 | 0.00 | 323 | 0.00 |
| 12 | -0.18 | 90 | -0.61 | 168 | 0.06 | 246 | 0.00 | 324 | 0.00 |
| 13 | -1.04 | 91 | -1.58 | 169 | -0.05 | 247 | 0.00 | 325 | 0.00 |
| 14 | -0.62 | 92 | -1.05 | 170 | 0.00 | 248 | 0.00 | 326 | 0.00 |
| 15 | -0.05 | 93 | -0.16 | 171 | 0.00 | 249 | 0.00 | 327 | 0.00 |
| 16 | -0.16 | 94 | 0.32 | 172 | 0.00 | 250 | 0.00 | 328 | 0.00 |
| 17 | -2.04 | 95 | -1.00 | 173 | 0.00 | 251 | 0.00 | 329 | 0.00 |
| 18 | -0.03 | 96 | -0.11 | 174 | 0.00 | 252 | 0.00 | 330 | 0.00 |
| 19 | 0.00 | 97 | -0.06 | 175 | 0.00 | 253 | 0.00 | 331 | 0.00 |
| 20 | 0.01 | 98 | 0.01 | 176 | 0.00 | 254 | 0.00 | 332 | 0.00 |
| 21 | -0.03 | 99 | 0.02 | 177 | 0.00 | 255 | 0.00 | 333 | 0.00 |
| 22 | 0.01 | 100 | 0.00 | 178 | 0.00 | 256 | 0.00 | 334 | 0.00 |
| 23 | 0.00 | 101 | 0.00 | 179 | 0.00 | 257 | 0.00 | 335 | 0.00 |
| 24 | 0.00 | 102 | 0.00 | 180 | 0.00 | 258 | 0.00 | 336 | 0.00 |
| 25 | 0.00 | 103 | 0.00 | 181 | 0.00 | 259 | 0.00 | 337 | 0.00 |
| 26 | 0.04 | 104 | 0.00 | 182 | 0.00 | 260 | 0.00 | 338 | 0.00 |
| 27 | 0.01 | 105 | 0.00 | 183 | 0.00 | 261 | 0.00 | 339 | 0.00 |
| 28 | 0.27 | 106 | 0.00 | 184 | 0.00 | 262 | 0.00 | 340 | 0.00 |
| 29 | 1.23 | 107 | 0.00 | 185 | 0.00 | 263 | 0.00 | 341 | 0.00 |
| 30 | -1.18 | 108 | 0.00 | 186 | 0.00 | 264 | 0.00 | 342 | 0.00 |
| 31 | -5.55 | 109 | -0.01 | 187 | 0.00 | 265 | 0.00 | 343 | 0.00 |
| 32 | -0.13 | 110 | 0.00 | 188 | 0.00 | 266 | 0.00 | 344 | 0.00 |
| 33 | -0.04 | 111 | 0.00 | 189 | 0.00 | 267 | 0.00 | 345 | 0.00 |
| 34 | 0.00 | 112 | 0.17 | 190 | 0.00 | 268 | 0.00 | 346 | -0.24 |
| 35 | 0.00 | 113 | -0.03 | 191 | 0.00 | 269 | 0.00 | 347 | 0.10 |
| 36 | 0.00 | 114 | -0.13 | 192 | 0.00 | 270 | 0.00 | 348 | 0.01 |
| 37 | 0.00 | 115 | -0.36 | 193 | 0.03 | 271 | 0.00 | 349 | -0.02 |
| 38 | 0.00 | 116 | -5.30 | 194 | 0.13 | 272 | 0.00 | 350 | -0.10 |

| | | | | | | | | | |
|---|---|---|---|---|---|---|---|---|---|
| 39 | 0.00 | 117 | -0.54 | 195 | -36.85 | 273 | 0.00 | 351 | -0.45 |
| 40 | 0.00 | 118 | -4.26 | 196 | 0.21 | 274 | 0.00 | 352 | 0.06 |
| 41 | 0.00 | 119 | -1.15 | 197 | 0.92 | 275 | 0.00 | 353 | -0.15 |
| 42 | 0.00 | 120 | -0.12 | 198 | -2.81 | 276 | 0.00 | 354 | 0.00 |
| 43 | 0.00 | 121 | -0.18 | 199 | 0.09 | 277 | 0.00 | 355 | 0.00 |
| 44 | 0.00 | 122 | 0.09 | 200 | -0.20 | 278 | 0.00 | 356 | 0.00 |
| 45 | 0.00 | 123 | -0.02 | 201 | -0.21 | 279 | 0.00 | 357 | 0.00 |
| 46 | 0.00 | 124 | 0.00 | 202 | -0.01 | 280 | 0.00 | 358 | 0.00 |
| 47 | 0.00 | 125 | 0.08 | 203 | 0.00 | 281 | 0.00 | 359 | 0.00 |
| 48 | 0.00 | 126 | 0.00 | 204 | 0.00 | 282 | 0.00 | 360 | 0.00 |
| 49 | 0.00 | 127 | 0.00 | 205 | 0.00 | 283 | 0.00 | 361 | 0.00 |
| 50 | 0.00 | 128 | 0.00 | 206 | 0.00 | 284 | 0.00 | 362 | 0.00 |
| 51 | 0.00 | 129 | 0.00 | 207 | 0.00 | 285 | 0.00 | 363 | 0.00 |
| 52 | 0.00 | 130 | -0.01 | 208 | 0.00 | 286 | 0.00 | 364 | -4.62 |
| 53 | 0.00 | 131 | 0.00 | 209 | 0.00 | 287 | 0.00 | 365 | 0.00 |
| 54 | 0.00 | 132 | 0.00 | 210 | 0.08 | 288 | 0.00 | 366 | -0.08 |
| 55 | 0.00 | 133 | 0.00 | 211 | 0.00 | 289 | 0.00 | 367 | 0.00 |
| 56 | 0.00 | 134 | 0.00 | 212 | 0.00 | 290 | 0.00 | 368 | -0.05 |
| 57 | 0.00 | 135 | 0.04 | 213 | 0.00 | 291 | 0.00 | 369 | -0.20 |
| 58 | 0.00 | 136 | 0.00 | 214 | 0.01 | 292 | 0.00 | 370 | 0.00 |
| 59 | 0.00 | 137 | 0.00 | 215 | 0.00 | 293 | 0.00 | 371 | 0.00 |
| 60 | 0.00 | 138 | 0.00 | 216 | 0.00 | 294 | 0.00 | 372 | 0.00 |
| 61 | 0.00 | 139 | 0.00 | 217 | 0.00 | 295 | 0.00 | 373 | 0.00 |
| 62 | 0.00 | 140 | 0.00 | 218 | 0.00 | 296 | 0.00 | 374 | 0.00 |
| 63 | 0.00 | 141 | 0.00 | 219 | 0.00 | 297 | 0.00 | 375 | 0.00 |
| 64 | 0.00 | 142 | 0.00 | 220 | 0.00 | 298 | 0.00 | 376 | 0.00 |
| 65 | 0.00 | 143 | 0.00 | 221 | 0.00 | 299 | 0.00 | 377 | 0.00 |
| 66 | 0.00 | 144 | 0.00 | 222 | 0.00 | 300 | 0.00 | 378 | 0.00 |
| 67 | 0.00 | 145 | 0.00 | 223 | 0.00 | 301 | 0.00 | 379 | 0.00 |
| 68 | 0.00 | 146 | 0.00 | 224 | 0.00 | 302 | 0.00 | 380 | 0.00 |
| 69 | 0.00 | 147 | 0.00 | 225 | -0.17 | 303 | 0.00 | 381 | 0.00 |
| 70 | -0.06 | 148 | 0.00 | 226 | 0.37 | 304 | 0.00 | 382 | 0.00 |
| 71 | 0.00 | 149 | 0.00 | 227 | -0.55 | 305 | 0.00 | 383 | 0.00 |
| 72 | 0.00 | 150 | 0.00 | 228 | -0.44 | 306 | 0.00 | 384 | 0.00 |
| 73 | 0.00 | 151 | 0.00 | 229 | 3.15 | 307 | 0.00 | 385 | 0.00 |
| 74 | 0.08 | 152 | 0.00 | 230 | -0.29 | 308 | 0.00 | 386 | 0.00 |
| 75 | 0.00 | 153 | 0.00 | 231 | -0.94 | 309 | 0.00 | 387 | 0.00 |
| 76 | 0.00 | 154 | 0.00 | 232 | -3.74 | 310 | 0.00 | 388 | 0.00 |
| 77 | 0.00 | 155 | 0.00 | 233 | 0.03 | 311 | 0.00 | 389 | 0.00 |
| 78 | 0.00 | 156 | 0.00 | 234 | 0.16 | 312 | 0.00 | | |

**Interaction between residues of chain B and the ligand bound to chain A:**

| Residue No. | TNBIE (kcal/mol) | Residue No. | TNBIE (kcal/mol) | Residue No. | TNBIE (kcal/mol) | Residue No. | TNBIE (kcal/mol) | Residue No. | TNBIE (kcal/mol) |
|---|---|---|---|---|---|---|---|---|---|
| 1 | 0.00 | 79 | 0.00 | 157 | 0.00 | 235 | 0.00 | 313 | 0.00 |
| 2 | 0.00 | 80 | 0.00 | 158 | 0.00 | 236 | 0.00 | 314 | 0.00 |
| 3 | 0.00 | 81 | 0.00 | 159 | 0.00 | 237 | 0.00 | 315 | 0.00 |
| 4 | 0.00 | 82 | 0.00 | 160 | 0.00 | 238 | 0.00 | 316 | 0.00 |
| 5 | 0.00 | 83 | 0.00 | 161 | 0.00 | 239 | 0.00 | 317 | 0.00 |
| 6 | 0.00 | 84 | 0.00 | 162 | 0.00 | 240 | 0.00 | 318 | 0.00 |
| 7 | 0.00 | 85 | 0.00 | 163 | 0.00 | 241 | 0.00 | 319 | 0.00 |
| 8 | 0.00 | 86 | 0.02 | 164 | 0.00 | 242 | 0.00 | 320 | 0.00 |
| 9 | 0.00 | 87 | 0.01 | 165 | 0.00 | 243 | 0.00 | 321 | 0.00 |
| 10 | 0.00 | 88 | 0.00 | 166 | 0.00 | 244 | 0.00 | 322 | 0.00 |
| 11 | 0.00 | 89 | -0.27 | 167 | 0.00 | 245 | 0.00 | 323 | 0.00 |
| 12 | 0.00 | 90 | 0.00 | 168 | 0.00 | 246 | 0.00 | 324 | 0.00 |
| 13 | 0.00 | 91 | 0.00 | 169 | 0.00 | 247 | 0.00 | 325 | 0.00 |
| 14 | 0.00 | 92 | 0.00 | 170 | 0.00 | 248 | 0.00 | 326 | 0.00 |
| 15 | 0.00 | 93 | 0.00 | 171 | 0.00 | 249 | 0.00 | 327 | 0.00 |
| 16 | 0.00 | 94 | 0.00 | 172 | 0.00 | 250 | 0.00 | 328 | 0.00 |
| 17 | 0.00 | 95 | 0.00 | 173 | 0.00 | 251 | -0.01 | 329 | 0.00 |
| 18 | 0.00 | 96 | 0.00 | 174 | 0.00 | 252 | -0.03 | 330 | 0.00 |
| 19 | 0.00 | 97 | 0.05 | 175 | 0.00 | 253 | -0.04 | 331 | 0.00 |
| 20 | 0.00 | 98 | 0.00 | 176 | 0.00 | 254 | -0.08 | 332 | 0.00 |
| 21 | 0.00 | 99 | 0.00 | 177 | 0.00 | 255 | -0.33 | 333 | 0.00 |
| 22 | 0.00 | 100 | 0.03 | 178 | 0.00 | 256 | -0.72 | 334 | 0.00 |
| 23 | 0.00 | 101 | 0.00 | 179 | 0.00 | 257 | -0.43 | 335 | 0.00 |
| 24 | 0.00 | 102 | 0.00 | 180 | 0.00 | 258 | -0.61 | 336 | 0.00 |
| 25 | 0.00 | 103 | 0.00 | 181 | 0.00 | 259 | -3.39 | 337 | 0.00 |
| 26 | 0.00 | 104 | 0.00 | 182 | 0.00 | 260 | -2.01 | 338 | 0.00 |
| 27 | 0.00 | 105 | 0.00 | 183 | 0.00 | 261 | -1.23 | 339 | 0.00 |
| 28 | 0.00 | 106 | 0.00 | 184 | 0.00 | 262 | -2.33 | 340 | 0.00 |
| 29 | 0.00 | 107 | 0.00 | 185 | 0.00 | 263 | -2.81 | 341 | 0.00 |
| 30 | 0.00 | 108 | 0.00 | 186 | 0.00 | 264 | 0.07 | 342 | 0.00 |
| 31 | 0.00 | 109 | 0.00 | 187 | 0.00 | 265 | -0.07 | 343 | 0.00 |
| 32 | 0.00 | 110 | 0.00 | 188 | 0.00 | 266 | -0.01 | 344 | 0.00 |
| 33 | 0.00 | 111 | 0.00 | 189 | 0.00 | 267 | 0.00 | 345 | 0.00 |
| 34 | 0.00 | 112 | 0.00 | 190 | 0.00 | 268 | 0.01 | 346 | 0.00 |
| 35 | 0.00 | 113 | 0.00 | 191 | 0.00 | 269 | 0.00 | 347 | 0.00 |
| 36 | 0.00 | 114 | 0.00 | 192 | 0.00 | 270 | 0.01 | 348 | 0.00 |
| 37 | 0.00 | 115 | 0.00 | 193 | 0.00 | 271 | 0.00 | 349 | 0.00 |
| 38 | 0.00 | 116 | 0.00 | 194 | 0.00 | 272 | 0.00 | 350 | 0.00 |
| 39 | 0.00 | 117 | 0.00 | 195 | 0.00 | 273 | 0.00 | 351 | 0.00 |
| 40 | 0.00 | 118 | 0.00 | 196 | 0.00 | 274 | 0.00 | 352 | 0.00 |
| 41 | 0.00 | 119 | 0.00 | 197 | 0.00 | 275 | 0.00 | 353 | 0.00 |
| 42 | 0.00 | 120 | 0.00 | 198 | 0.00 | 276 | 0.00 | 354 | 0.00 |
| 43 | 0.00 | 121 | 0.00 | 199 | 0.00 | 277 | 0.00 | 355 | 0.00 |
| 44 | 0.00 | 122 | 0.00 | 200 | 0.00 | 278 | 0.00 | 356 | 0.00 |
| 45 | 0.00 | 123 | 0.00 | 201 | 0.00 | 279 | 0.00 | 357 | 0.00 |
| 46 | 0.00 | 124 | 0.00 | 202 | 0.00 | 280 | 0.00 | 358 | 0.00 |
| 47 | 0.00 | 125 | 0.00 | 203 | 0.00 | 281 | 0.00 | 359 | 0.00 |

| | | | | | | | | | |
|---|---|---|---|---|---|---|---|---|---|
| 48 | 0.00 | 126 | 0.00 | 204 | 0.00 | 282 | 0.00 | 360 | 0.00 |
| 49 | 0.00 | 127 | 0.00 | 205 | 0.00 | 283 | 0.00 | 361 | 0.00 |
| 50 | 0.00 | 128 | 0.00 | 206 | 0.00 | 284 | 0.00 | 362 | 0.00 |
| 51 | 0.00 | 129 | 0.00 | 207 | 0.00 | 285 | 0.00 | 363 | 0.00 |
| 52 | 0.00 | 130 | 0.00 | 208 | 0.00 | 286 | 0.00 | 364 | 0.00 |
| 53 | 0.00 | 131 | 0.00 | 209 | 0.00 | 287 | 0.00 | 365 | 0.00 |
| 54 | 0.00 | 132 | 0.00 | 210 | 0.00 | 288 | 0.00 | 366 | 0.00 |
| 55 | 0.00 | 133 | 0.00 | 211 | 0.00 | 289 | 0.00 | 367 | 0.00 |
| 56 | 0.00 | 134 | 0.00 | 212 | 0.00 | 290 | 0.00 | 368 | 0.00 |
| 57 | 0.00 | 135 | 0.00 | 213 | 0.00 | 291 | 0.00 | 369 | 0.00 |
| 58 | 0.00 | 136 | 0.00 | 214 | 0.00 | 292 | 0.00 | 370 | 0.00 |
| 59 | -0.01 | 137 | 0.00 | 215 | 0.00 | 293 | 0.00 | 371 | 0.00 |
| 60 | 0.00 | 138 | 0.00 | 216 | 0.00 | 294 | 0.00 | 372 | 0.00 |
| 61 | 0.00 | 139 | 0.00 | 217 | 0.00 | 295 | 0.00 | 373 | 0.00 |
| 62 | 0.00 | 140 | 0.00 | 218 | 0.00 | 296 | 0.00 | 374 | 0.00 |
| 63 | -0.14 | 141 | 0.00 | 219 | 0.00 | 297 | 0.00 | 375 | 0.00 |
| 64 | 0.00 | 142 | 0.00 | 220 | 0.00 | 298 | 0.00 | 376 | 0.00 |
| 65 | 0.00 | 143 | 0.00 | 221 | 0.00 | 299 | 0.00 | 377 | 0.00 |
| 66 | 0.00 | 144 | 0.00 | 222 | 0.00 | 300 | 0.00 | 378 | 0.00 |
| 67 | 0.00 | 145 | 0.00 | 223 | 0.00 | 301 | 0.00 | 379 | 0.00 |
| 68 | 0.00 | 146 | 0.00 | 224 | 0.00 | 302 | 0.00 | 380 | 0.00 |
| 69 | 0.00 | 147 | 0.00 | 225 | 0.00 | 303 | 0.00 | 381 | 0.00 |
| 70 | 0.00 | 148 | 0.00 | 226 | 0.00 | 304 | 0.00 | 382 | 0.00 |
| 71 | 0.00 | 149 | 0.00 | 227 | 0.00 | 305 | 0.00 | 383 | 0.00 |
| 72 | 0.00 | 150 | 0.00 | 228 | 0.00 | 306 | 0.00 | 384 | 0.00 |
| 73 | 0.00 | 151 | 0.00 | 229 | 0.00 | 307 | 0.00 | 385 | 0.00 |
| 74 | 0.00 | 152 | 0.00 | 230 | 0.00 | 308 | 0.00 | 386 | 0.00 |
| 75 | 0.00 | 153 | 0.00 | 231 | 0.00 | 309 | 0.00 | 387 | 0.00 |
| 76 | 0.00 | 154 | 0.00 | 232 | 0.00 | 310 | 0.00 | 388 | 0.00 |
| 77 | 0.00 | 155 | 0.00 | 233 | 0.00 | 311 | 0.00 | 389 | 0.00 |
| 78 | 0.00 | 156 | 0.00 | 234 | 0.00 | 312 | 0.00 | | |

**Interaction between residues of chain A and the ligand bound to chain B:**

| Residue No. | TNBIE (kcal/mol) | Residue No. | TNBIE (kcal/mol) | Residue No. | TNBIE (kcal/mol) | Residue No. | TNBIE (kcal/mol) | Residue No. | TNBIE (kcal/mol) |
|---|---|---|---|---|---|---|---|---|---|
| 1 | 0.00 | 79 | 0.00 | 157 | 0.00 | 235 | 0.00 | 313 | 0.00 |
| 2 | 0.00 | 80 | 0.00 | 158 | 0.00 | 236 | 0.00 | 314 | 0.00 |
| 3 | 0.00 | 81 | 0.00 | 159 | 0.00 | 237 | 0.00 | 315 | 0.00 |
| 4 | 0.00 | 82 | 0.00 | 160 | 0.00 | 238 | 0.00 | 316 | 0.00 |
| 5 | 0.00 | 83 | 0.00 | 161 | 0.00 | 239 | 0.00 | 317 | 0.00 |
| 6 | 0.00 | 84 | 0.00 | 162 | 0.00 | 240 | 0.00 | 318 | 0.00 |
| 7 | 0.00 | 85 | 0.00 | 163 | 0.00 | 241 | 0.00 | 319 | 0.00 |
| 8 | 0.00 | 86 | 0.03 | 164 | 0.00 | 242 | 0.00 | 320 | 0.00 |
| 9 | 0.00 | 87 | 0.07 | 165 | 0.00 | 243 | 0.00 | 321 | 0.00 |
| 10 | 0.00 | 88 | 0.04 | 166 | 0.00 | 244 | 0.00 | 322 | 0.00 |
| 11 | 0.00 | 89 | -0.04 | 167 | 0.00 | 245 | 0.00 | 323 | 0.00 |
| 12 | 0.00 | 90 | 0.00 | 168 | 0.00 | 246 | 0.00 | 324 | 0.00 |
| 13 | 0.00 | 91 | 0.00 | 169 | 0.00 | 247 | 0.00 | 325 | 0.00 |
| 14 | 0.00 | 92 | 0.00 | 170 | 0.00 | 248 | 0.00 | 326 | 0.00 |
| 15 | 0.00 | 93 | -0.02 | 171 | 0.00 | 249 | 0.00 | 327 | 0.00 |
| 16 | 0.00 | 94 | 0.00 | 172 | 0.00 | 250 | 0.00 | 328 | 0.00 |
| 17 | 0.00 | 95 | 0.00 | 173 | 0.00 | 251 | 0.00 | 329 | 0.00 |
| 18 | 0.00 | 96 | 0.00 | 174 | 0.00 | 252 | -0.02 | 330 | 0.00 |
| 19 | 0.00 | 97 | 0.07 | 175 | 0.00 | 253 | -0.03 | 331 | 0.00 |
| 20 | 0.00 | 98 | 0.00 | 176 | 0.00 | 254 | -0.18 | 332 | 0.00 |
| 21 | 0.00 | 99 | 0.00 | 177 | 0.00 | 255 | -0.32 | 333 | 0.00 |
| 22 | 0.00 | 100 | 0.01 | 178 | 0.00 | 256 | -0.82 | 334 | 0.00 |
| 23 | 0.00 | 101 | 0.00 | 179 | 0.00 | 257 | -0.70 | 335 | 0.00 |
| 24 | 0.00 | 102 | 0.00 | 180 | 0.00 | 258 | -0.49 | 336 | 0.00 |
| 25 | 0.00 | 103 | 0.00 | 181 | 0.00 | 259 | -2.17 | 337 | 0.00 |
| 26 | 0.00 | 104 | 0.00 | 182 | 0.00 | 260 | -5.29 | 338 | 0.00 |
| 27 | 0.00 | 105 | 0.00 | 183 | 0.00 | 261 | 0.28 | 339 | 0.00 |
| 28 | 0.00 | 106 | 0.00 | 184 | 0.00 | 262 | -1.34 | 340 | 0.00 |
| 29 | 0.00 | 107 | 0.00 | 185 | 0.00 | 263 | -0.97 | 341 | 0.00 |
| 30 | 0.00 | 108 | 0.00 | 186 | 0.00 | 264 | -0.22 | 342 | 0.00 |
| 31 | 0.00 | 109 | 0.00 | 187 | 0.00 | 265 | -0.01 | 343 | 0.00 |
| 32 | 0.00 | 110 | 0.00 | 188 | 0.00 | 266 | 0.05 | 344 | 0.00 |
| 33 | 0.00 | 111 | 0.00 | 189 | 0.00 | 267 | 0.03 | 345 | 0.00 |
| 34 | 0.00 | 112 | 0.00 | 190 | 0.00 | 268 | 0.01 | 346 | 0.00 |
| 35 | 0.00 | 113 | 0.00 | 191 | 0.00 | 269 | 0.00 | 347 | 0.00 |
| 36 | 0.00 | 114 | 0.00 | 192 | 0.00 | 270 | 0.01 | 348 | 0.00 |
| 37 | 0.00 | 115 | 0.00 | 193 | 0.00 | 271 | 0.00 | 349 | 0.00 |
| 38 | 0.00 | 116 | 0.00 | 194 | 0.00 | 272 | 0.00 | 350 | 0.00 |
| 39 | 0.00 | 117 | 0.00 | 195 | 0.00 | 273 | 0.00 | 351 | 0.00 |
| 40 | 0.00 | 118 | 0.00 | 196 | 0.00 | 274 | 0.00 | 352 | 0.00 |
| 41 | 0.00 | 119 | 0.00 | 197 | 0.00 | 275 | 0.00 | 353 | 0.00 |
| 42 | 0.00 | 120 | 0.00 | 198 | 0.00 | 276 | 0.00 | 354 | 0.00 |
| 43 | 0.00 | 121 | 0.00 | 199 | 0.00 | 277 | 0.00 | 355 | 0.00 |
| 44 | 0.00 | 122 | 0.00 | 200 | 0.00 | 278 | 0.00 | 356 | 0.00 |
| 45 | 0.00 | 123 | 0.00 | 201 | 0.00 | 279 | 0.00 | 357 | 0.00 |

| | | | | | | | | | |
|---|---|---|---|---|---|---|---|---|---|
| 46 | 0.00 | 124 | 0.00 | 202 | 0.00 | 280 | 0.00 | 358 | 0.00 |
| 47 | 0.00 | 125 | 0.00 | 203 | 0.00 | 281 | 0.00 | 359 | 0.00 |
| 48 | 0.00 | 126 | 0.00 | 204 | 0.00 | 282 | 0.00 | 360 | 0.00 |
| 49 | 0.00 | 127 | 0.00 | 205 | 0.00 | 283 | 0.00 | 361 | 0.00 |
| 50 | 0.00 | 128 | 0.00 | 206 | 0.00 | 284 | 0.00 | 362 | 0.00 |
| 51 | 0.00 | 129 | 0.00 | 207 | 0.00 | 285 | 0.00 | 363 | 0.00 |
| 52 | 0.00 | 130 | 0.00 | 208 | 0.00 | 286 | 0.00 | 364 | 0.00 |
| 53 | 0.00 | 131 | 0.00 | 209 | 0.00 | 287 | 0.00 | 365 | 0.00 |
| 54 | 0.00 | 132 | 0.00 | 210 | 0.00 | 288 | 0.00 | 366 | 0.00 |
| 55 | 0.00 | 133 | 0.00 | 211 | 0.00 | 289 | 0.00 | 367 | 0.00 |
| 56 | 0.00 | 134 | 0.00 | 212 | 0.00 | 290 | 0.00 | 368 | 0.00 |
| 57 | 0.00 | 135 | 0.00 | 213 | 0.00 | 291 | 0.00 | 369 | 0.00 |
| 58 | 0.00 | 136 | 0.00 | 214 | 0.00 | 292 | 0.00 | 370 | 0.00 |
| 59 | -0.03 | 137 | 0.00 | 215 | 0.00 | 293 | 0.00 | 371 | 0.00 |
| 60 | 0.00 | 138 | 0.00 | 216 | 0.00 | 294 | 0.00 | 372 | 0.00 |
| 61 | 0.00 | 139 | 0.00 | 217 | 0.00 | 295 | 0.00 | 373 | 0.00 |
| 62 | 0.00 | 140 | 0.00 | 218 | 0.00 | 296 | 0.00 | 374 | 0.00 |
| 63 | -0.09 | 141 | 0.00 | 219 | 0.00 | 297 | 0.00 | 375 | 0.00 |
| 64 | 0.00 | 142 | 0.00 | 220 | 0.00 | 298 | 0.00 | 376 | 0.00 |
| 65 | 0.00 | 143 | 0.00 | 221 | 0.00 | 299 | 0.00 | 377 | 0.00 |
| 66 | 0.00 | 144 | 0.00 | 222 | 0.00 | 300 | 0.00 | 378 | 0.00 |
| 67 | 0.00 | 145 | 0.00 | 223 | 0.00 | 301 | 0.00 | 379 | 0.00 |
| 68 | 0.00 | 146 | 0.00 | 224 | 0.00 | 302 | 0.00 | 380 | 0.00 |
| 69 | 0.00 | 147 | 0.00 | 225 | 0.00 | 303 | 0.00 | 381 | 0.00 |
| 70 | 0.00 | 148 | 0.00 | 226 | 0.00 | 304 | 0.00 | 382 | 0.00 |
| 71 | 0.00 | 149 | 0.00 | 227 | 0.00 | 305 | 0.00 | 383 | 0.00 |
| 72 | 0.00 | 150 | 0.00 | 228 | 0.00 | 306 | 0.00 | 384 | 0.00 |
| 73 | 0.00 | 151 | 0.00 | 229 | 0.00 | 307 | 0.00 | 385 | 0.00 |
| 74 | 0.00 | 152 | 0.00 | 230 | 0.00 | 308 | 0.00 | 386 | 0.00 |
| 75 | 0.00 | 153 | 0.00 | 231 | 0.00 | 309 | 0.00 | 387 | 0.00 |
| 76 | 0.00 | 154 | 0.00 | 232 | 0.00 | 310 | 0.00 | 388 | 0.00 |
| 77 | 0.00 | 155 | 0.00 | 233 | 0.00 | 311 | 0.00 | 389 | 0.00 |
| 78 | 0.00 | 156 | 0.00 | 234 | 0.00 | 312 | 0.00 | | |

**Interaction between residues of chain B and the ligand bound to chain B:**

| Residue No. | TNBIE (kcal/mol) | Residue No. | TNBIE (kcal/mol) | Residue No. | TNBIE (kcal/mol) | Residue No. | TNBIE (kcal/mol) | Residue No. | TNBIE (kcal/mol) |
|---|---|---|---|---|---|---|---|---|---|
| 1 | 0.00 | 79 | 0.00 | 157 | -0.01 | 235 | 0.00 | 313 | 0.00 |
| 2 | 0.00 | 80 | 0.00 | 158 | 0.22 | 236 | -0.03 | 314 | 0.00 |
| 3 | 0.00 | 81 | 0.00 | 159 | -0.37 | 237 | -0.04 | 315 | -0.01 |
| 4 | 0.00 | 82 | 0.00 | 160 | 0.55 | 238 | 1.34 | 316 | 0.01 |
| 5 | 0.00 | 83 | 0.00 | 161 | -0.39 | 239 | 0.01 | 317 | -0.27 |
| 6 | 0.00 | 84 | 0.00 | 162 | -0.51 | 240 | -4.64 | 318 | 0.12 |
| 7 | 0.00 | 85 | 0.00 | 163 | 0.06 | 241 | 0.01 | 319 | -0.23 |
| 8 | 0.01 | 86 | -0.01 | 164 | -0.41 | 242 | -0.07 | 320 | 0.12 |
| 9 | 0.02 | 87 | 0.00 | 165 | -0.01 | 243 | -0.12 | 321 | -0.03 |
| 10 | 0.00 | 88 | 0.00 | 166 | 0.04 | 244 | 0.05 | 322 | -0.01 |
| 11 | -0.01 | 89 | -0.07 | 167 | -0.04 | 245 | 0.00 | 323 | 0.00 |
| 12 | -1.36 | 90 | -0.50 | 168 | -0.01 | 246 | 0.00 | 324 | 0.00 |
| 13 | 0.02 | 91 | 0.04 | 169 | 0.00 | 247 | 0.00 | 325 | 0.00 |
| 14 | -0.76 | 92 | 0.19 | 170 | 0.00 | 248 | 0.00 | 326 | 0.00 |
| 15 | 0.08 | 93 | -0.02 | 171 | 0.00 | 249 | 0.00 | 327 | 0.00 |
| 16 | 0.12 | 94 | -0.62 | 172 | 0.00 | 250 | 0.00 | 328 | 0.00 |
| 17 | -1.05 | 95 | -5.07 | 173 | 0.00 | 251 | 0.00 | 329 | 0.00 |
| 18 | -0.04 | 96 | -0.45 | 174 | 0.00 | 252 | 0.00 | 330 | 0.00 |
| 19 | 0.08 | 97 | -0.31 | 175 | 0.00 | 253 | 0.00 | 331 | 0.00 |
| 20 | 0.32 | 98 | -0.35 | 176 | 0.00 | 254 | 0.00 | 332 | 0.00 |
| 21 | 0.19 | 99 | -0.15 | 177 | 0.00 | 255 | 0.00 | 333 | 0.00 |
| 22 | 0.01 | 100 | -0.03 | 178 | 0.00 | 256 | 0.00 | 334 | 0.00 |
| 23 | 0.04 | 101 | 0.00 | 179 | 0.00 | 257 | 0.00 | 335 | -0.01 |
| 24 | 0.00 | 102 | -0.03 | 180 | 0.00 | 258 | 0.00 | 336 | 0.00 |
| 25 | -0.01 | 103 | 0.00 | 181 | 0.00 | 259 | 0.00 | 337 | 0.00 |
| 26 | 0.20 | 104 | 0.00 | 182 | 0.00 | 260 | 0.00 | 338 | 0.00 |
| 27 | 0.16 | 105 | 0.00 | 183 | 0.00 | 261 | 0.00 | 339 | 0.00 |
| 28 | 0.41 | 106 | 0.00 | 184 | 0.00 | 262 | 0.00 | 340 | 0.00 |
| 29 | -0.06 | 107 | 0.00 | 185 | 0.00 | 263 | 0.00 | 341 | 0.00 |
| 30 | -1.06 | 108 | 0.00 | 186 | 0.00 | 264 | 0.00 | 342 | 0.00 |
| 31 | -2.27 | 109 | -0.04 | 187 | 0.00 | 265 | 0.00 | 343 | 0.00 |
| 32 | -0.03 | 110 | -0.01 | 188 | 0.00 | 266 | 0.00 | 344 | 0.14 |
| 33 | 0.02 | 111 | 0.04 | 189 | 0.00 | 267 | 0.00 | 345 | -0.16 |
| 34 | -0.01 | 112 | 0.08 | 190 | 0.00 | 268 | 0.00 | 346 | 0.07 |
| 35 | 0.00 | 113 | 0.00 | 191 | 0.00 | 269 | 0.00 | 347 | 0.03 |
| 36 | 0.00 | 114 | -0.12 | 192 | 0.00 | 270 | 0.00 | 348 | -0.02 |
| 37 | 0.00 | 115 | 0.11 | 193 | -0.02 | 271 | 0.00 | 349 | -0.01 |
| 38 | 0.00 | 116 | -0.04 | 194 | 0.13 | 272 | 0.00 | 350 | -0.11 |
| 39 | 0.00 | 117 | -0.29 | 195 | -29.86 | 273 | 0.00 | 351 | -0.22 |
| 40 | 0.00 | 118 | -3.11 | 196 | 0.28 | 274 | 0.00 | 352 | 0.00 |
| 41 | 0.00 | 119 | -0.11 | 197 | 2.01 | 275 | 0.00 | 353 | -0.01 |
| 42 | 0.00 | 120 | 0.15 | 198 | 0.54 | 276 | 0.00 | 354 | 0.00 |
| 43 | 0.00 | 121 | -0.83 | 199 | 0.13 | 277 | 0.00 | 355 | 0.00 |
| 44 | 0.00 | 122 | 0.16 | 200 | -0.13 | 278 | 0.00 | 356 | 0.00 |
| 45 | 0.00 | 123 | -0.01 | 201 | -0.18 | 279 | 0.00 | 357 | 0.00 |

| | | | | | | | | | |
|---|---|---|---|---|---|---|---|---|---|
| 46 | 0.00 | 124 | 0.01 | 202 | -0.02 | 280 | 0.00 | 358 | 0.00 |
| 47 | 0.00 | 125 | 0.51 | 203 | 0.00 | 281 | 0.00 | 359 | 0.00 |
| 48 | 0.00 | 126 | 0.01 | 204 | 0.00 | 282 | 0.00 | 360 | 0.00 |
| 49 | 0.00 | 127 | 0.00 | 205 | 0.00 | 283 | 0.00 | 361 | 0.00 |
| 50 | 0.00 | 128 | -0.01 | 206 | 0.00 | 284 | 0.00 | 362 | 0.00 |
| 51 | 0.00 | 129 | 0.00 | 207 | 0.00 | 285 | -0.01 | 363 | -0.04 |
| 52 | 0.00 | 130 | -0.02 | 208 | 0.00 | 286 | 0.00 | 364 | -11.79 |
| 53 | 0.00 | 131 | 0.00 | 209 | 0.00 | 287 | 0.00 | 365 | 0.07 |
| 54 | 0.00 | 132 | 0.00 | 210 | 0.06 | 288 | 0.00 | 366 | -0.22 |
| 55 | 0.00 | 133 | 0.00 | 211 | 0.00 | 289 | 0.00 | 367 | -0.07 |
| 56 | 0.00 | 134 | 0.00 | 212 | 0.00 | 290 | 0.00 | 368 | -0.12 |
| 57 | 0.00 | 135 | 0.00 | 213 | 0.00 | 291 | 0.00 | 369 | 0.10 |
| 58 | 0.00 | 136 | 0.00 | 214 | 0.00 | 292 | -0.02 | 370 | 0.00 |
| 59 | 0.00 | 137 | 0.00 | 215 | 0.00 | 293 | 0.00 | 371 | 0.00 |
| 60 | 0.00 | 138 | 0.00 | 216 | 0.00 | 294 | 0.00 | 372 | 0.00 |
| 61 | 0.00 | 139 | 0.00 | 217 | 0.00 | 295 | 0.00 | 373 | 0.00 |
| 62 | 0.00 | 140 | 0.00 | 218 | 0.00 | 296 | 0.00 | 374 | 0.00 |
| 63 | 0.00 | 141 | 0.00 | 219 | 0.00 | 297 | 0.00 | 375 | 0.00 |
| 64 | 0.00 | 142 | 0.00 | 220 | 0.00 | 298 | 0.00 | 376 | 0.00 |
| 65 | 0.00 | 143 | 0.00 | 221 | 0.00 | 299 | 0.00 | 377 | 0.00 |
| 66 | 0.00 | 144 | 0.00 | 222 | 0.00 | 300 | 0.00 | 378 | 0.00 |
| 67 | 0.00 | 145 | 0.00 | 223 | 0.00 | 301 | 0.00 | 379 | 0.00 |
| 68 | 0.00 | 146 | 0.00 | 224 | 0.00 | 302 | 0.00 | 380 | 0.00 |
| 69 | 0.00 | 147 | 0.00 | 225 | -0.16 | 303 | 0.00 | 381 | 0.00 |
| 70 | -0.01 | 148 | 0.00 | 226 | 0.30 | 304 | 0.00 | 382 | 0.00 |
| 71 | 0.00 | 149 | 0.00 | 227 | -0.15 | 305 | 0.00 | 383 | 0.00 |
| 72 | 0.00 | 150 | 0.00 | 228 | -0.20 | 306 | 0.00 | 384 | 0.00 |
| 73 | 0.00 | 151 | 0.00 | 229 | 3.51 | 307 | 0.00 | 385 | 0.00 |
| 74 | 0.00 | 152 | 0.00 | 230 | -0.27 | 308 | 0.00 | 386 | 0.00 |
| 75 | 0.00 | 153 | 0.00 | 231 | -0.60 | 309 | 0.00 | 387 | 0.00 |
| 76 | 0.00 | 154 | 0.00 | 232 | -12.79 | 310 | 0.00 | 388 | 0.00 |
| 77 | 0.00 | 155 | 0.00 | 233 | 0.21 | 311 | 0.00 | 389 | 0.00 |
| 78 | 0.00 | 156 | 0.00 | 234 | 0.03 | 312 | 0.00 | | |

**Figure S1: Agarose gel electrophoresis analysis of isolated genomic DNA.** Agarose gel image of isolated genomic DNA from the *Streptromyces rimofaciens* B-98891 strain (lane **1**), which was used as a template for cloning *milM* (M represents DNA marker).

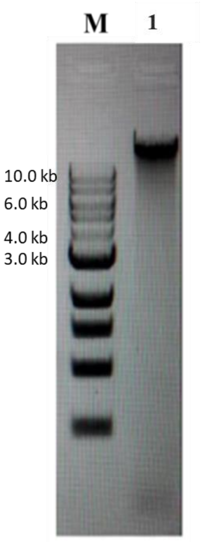

**Figure S2: SDS-PAGE analysis of the proteins used in this study.** Molecular weight of MilM-His$_6$-WT and its variants ~44.5 kDa (M represents protein marker).

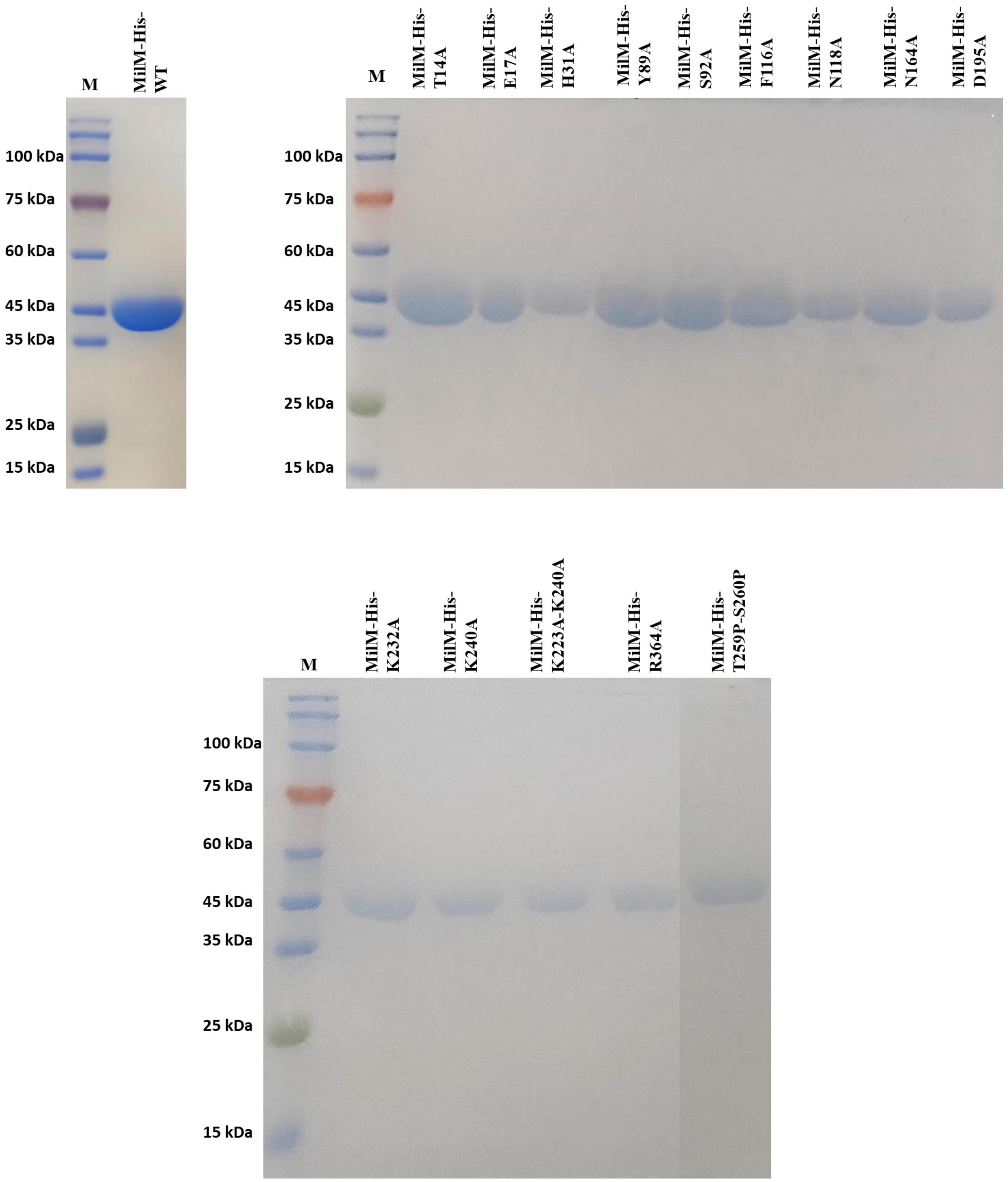

**Figure S3: Spectroscopic and chromatographic characterization of the PLP cofactor in MilM.**

**(A)** Image of the as isolated MilM-WT which is yellow in color indicating the presence of the PLP cofactor.

**(B)** UV-Vis absorption spectra of purified MilM (green), boiled MilM (red), and free PLP standard (blue) recorded in 50 mM HEPES buffer (pH 7.5). Native MilM exhibits a distinct absorption maximum at 420 nm, characteristic of the internal aldimine[2] (**17**) formed between the active-site Lys residue and the PLP cofactor (**16**). In contrast, the supernatant obtained from boiled MilM displays a peak near 392 nm, co-relating with the free PLP standard, and confirming loss of the bound PLP upon denaturation.

**(C)** HPLC chromatograms of PLP standard (blue) and supernatant obtained from boiled MilM (red) samples, monitored at 390 nm. The presence of a distinct PLP peak (Rt 10.8 min) in the boiled MilM sample, which co-elutes with an authentic PLP standard in HPLC further validates the presence of the PLP cofactor in MilM active as a covalent adduct.

**(D)** UV-Vis spectra of PLP standard with different concentrations (25-500 µM) and 100 µM boiled MilM (yellow) showing a quantitative analysis of PLP bound to MilM. The PLP:protein stoichiometry from this analysis was found to be 1:1, indicating that 100% MilM is bound with the cofactor PLP.

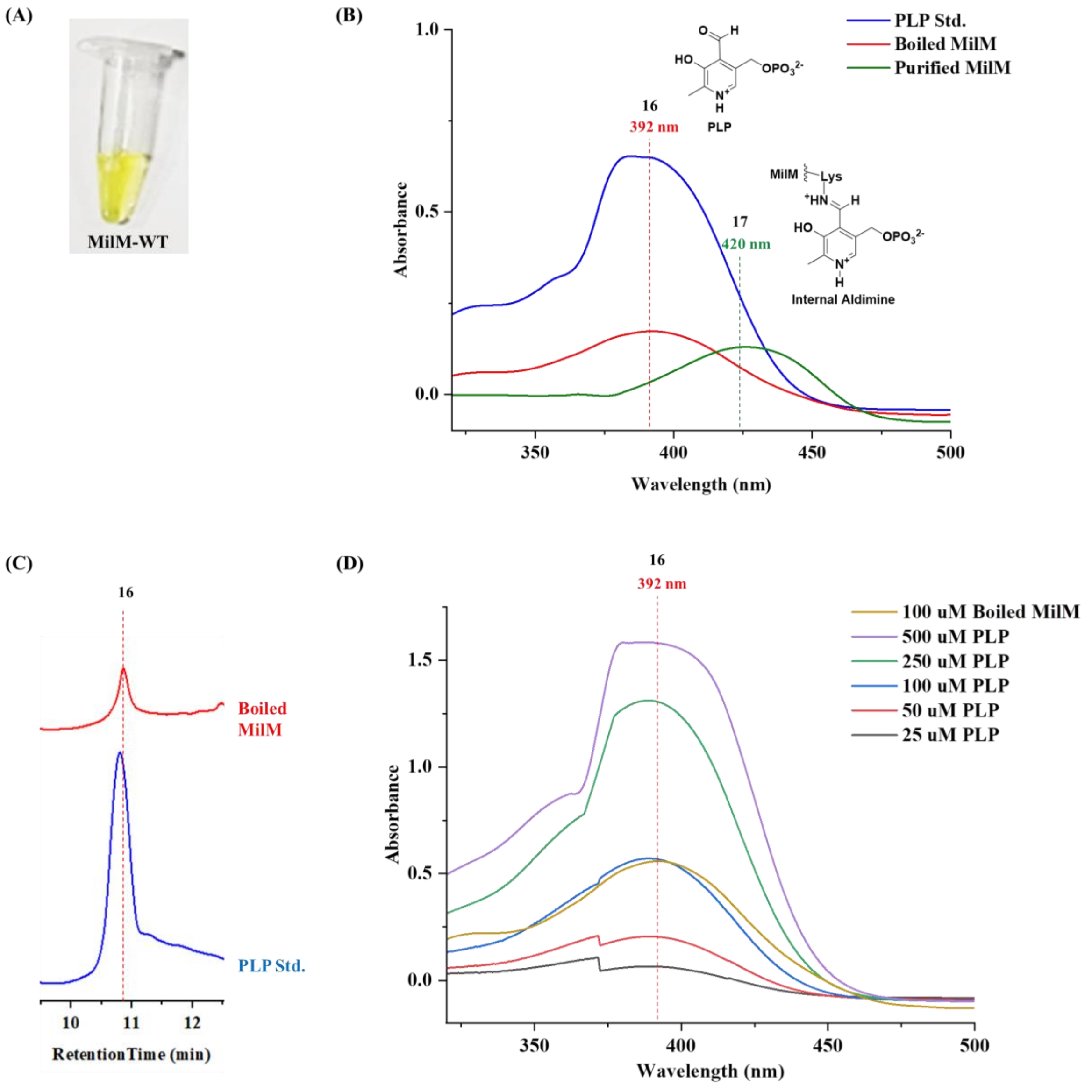

**Figure S4**: **Sequence alignment of MilM with homologous enzymes.** Three sequences (G8WNKC, $O_2$-, PLP-dependent L-Arginine hydroxylase RohP from *Streptantibioticus cattleyicolor*; A0A0D4BS17, Type I PLP-dependent aspartate aminotransferase-like Ind4 from *Streptomyces griseus subsp. griseus*; A0A0X1KHF5 PLP-dependent L-arginine hydroxylase MppP from *Streptomyces wadayamensis*) were obtained from NCBI BLAST analysis and the alignment was generated using Clustal Omega.[3] Conserved motifs in MilM include: PLP binding residues (in yellow) showing the catalytic Lys232 (in cyan) which forms the internal aldimine, L-Arg binding residues (in pink) with the catalytic His31 (in green) which is proposed to activate the water molecule for hydroxylation step and dimer interface residues (in grey). The numbing of amino acids in respect to MilM are mentioned in red.

```
                           14 17            31
MilM       ---MSDTLAHNRPLDLTQHEIAALRSEHNLADAHTHQYQSPAQQLIVDSLPALWHEAEKG        57
RohP       MHPQATPAPGAPLLDLTQHEIQALTMKYNLADAHTHQRQSASQQSIVSRLPQLWYEAEEG        60
MppP       -----MTTQPQLKENLTQWEYLALNSELNIADGHARQALSPGQQKIVNELPVLWAESEQR        55
Ind4       ---------MERFNNLTRYEHVGINQAINLADGHAHQGQNSTQQAIVADLPNIFEAAENA        51
                         :**: *  .:    *:**.*::*  .  ** **  ** ::  :*:

                                           89 92                    116
MilM       RQADFEQRFIEAFFRLHGQPTAIGL-DRTLLTYAASISTMIAGMFLKRRDARVTLVEPCF        116
RohP       LQATYEKRFTEAFFQLHRQPTALVK-NKTMLSYAASISTMVAGMFLKKERLAVTLIEPCF        119
MppP       PVQQIESEAHQAYFTLLGQHGYPAEPGRVLSCYSSSVSMEILARSLSASVDRVALVHPTF        115
Ind4       LQATTEREFQRAFYTLAGQHSAVDH-PRTMLCYSASLSTDLVATFLASRNLTVGLLQPCF        110
                *  .  .*:: *  *        :.:  *::*:*  : .  *      * *:.* *

           118                                           164
MilM       DNLPDLLVNLGVPLTALPEDALRDPARIHRELSRLVTTEALFLVDPNNPTGHSLFADGMR        176
RohP       DNLYDVLANMDVPLYPIDESVFYDVDRIYPELERRVRTDALFLVDPNNPTGFSLLRHGRK        179
MppP       DNIADLLRGNGLDLVPVEEDALHGAD-LSAE--LLSSVGCVFVTTPNNPTGRVLAE---E        169
Ind4       DNLATILRRRQVKVVPLGEEQFRPES-LDRT-FAELNTDAVFLTLPNNPTGFHLDQ---E        165
           **:  :*    : :  : *. :     :         . .:*:. ******  *     .

                            195                                  232
MilM       GFEEVVRFCRERGTVLVLDLCFAAFALGSGGPGRHDVYELLENSGVTYIAMEDTGKTWPV        236
RohP       GFEEVVRFCKDHDKLLLIDFCFASFTLFEPELARFDMYELLENSGVRYLAIEDTGKTWPV        239
MppP       RLRRLAEQCAEHGTVLALDTSFRGFDA----AAHYDHYAVLQEAGCRWVVIEDTGKLWPT        225
Ind4       QFRRVIDLCAEHQKILIVDCTFRFFAP----TPFWDQYAQLEESGISYVVVEDTGKTWPT        221
            :..:   * :: .:* :*  *  *          * *  *:::*  ::.:***** **.

             240                259
MilM       QDAKCALLTTSADIYPAVYNLHTSVLLNVSPFILNTLTRYIEDSRRDGFAS-VTDVLERN        295
RohP       QDAKCALITASDDIWETVYNLHTSVLLNVSPFVLNMLTQYVRDSAADRLAS-VREVLTRN        298
MppP       LDLKAGLLVFSEDIGLPVEKIYSDILLGVSPLILALIREFSRDAADGGLAD-LHAFILHN        284
Ind4       QDLKCSILAVSTDLHESVLELHNDILLNVSPFILQLLTAYLRDTERNGLQQTIWSVVRAN        281
            * *..::. * *:   * :::..:**.***::*  :  : .*:  . : . :  .:  *

MilM       RKSLRAATEGTV-LRAHEPDVPVSVAWFTIDDRGPDATQLQRDLSGHGIHVLPGTYFYWN        354
RohP       RECARKTLDGSI-LEYQEPVVKVSVAWFRVDHPELTATDVHRLLSADGVYVLPGRYFYWS        357
MppP       RSVVRRALAGVEGVSFPDPESRSSVE--RVAFAGRTGTEVWEELQRHHVFALPCRQFHWA        342
Ind4       RISLRKALEGTV-LVPAHPESTISVEWVRIDHPSLRSDDLVEMLAKSQVGILPGDHFYWD        340
           *   * :  *   :   .*    **    :      . :: . *    :  **   *:*

                   364
MilM       EPSRGERYVRVALARDPGEFDASMARLRTLLARYA-------- 389
RohP       EPSKGDAYVRMALAREPEMFADAMALTRQVLDRHGR------- 393
MppP       EPSDGDHMVRIALSRSTEPLEKSVQVLRTVLETR--------- 376
Ind4       DHETGSHFIRFALAREERWFATACGSLREALLARPELIEGAGS 383
           : . *.  :*.**:*.   :  :    *  *
```

**Figure S5: Spectroscopic and analytical characterization of MilM lysine mutants.**

**(A)** Images of purified MilM-K232A (upper) and MilM-K232A-K240A (lower). The yellow coloration observed for the K232A mutant indicates the presence of the PLP cofactor in its internal aldimine form, whereas the double mutant displays a loss of visible PLP color, consistent with disrupted cofactor binding.

**(B)** UV-Vis absorption spectra of the MilM-K232A (green) and MilM-K232A-K240A (blue) variants. MilM-K232A exhibits a strong absorbance band at ~420 nm, characteristic of the PLP internal aldimine, while this feature is absent in the double mutant, indicating the loss of bound PLP.

**(C)** LCMS analysis of the MilM-K232A reaction with L-Arg showing only the unreacted substrate mass (*m/z* = 175.1197 $[M + H]^+$, calc. 175.1190) after the reaction, indicating this mutant is catalytically inactive and suggesting that Lys232 is the residue involved in internal aldimine.

**(D)** LCMS analysis of the MilM-K232A-K240A reaction under identical conditions shows a similar reaction profile with only substrate mass at *m/z* = 175.1195 ($[M + H]^+$, calc. 175.1190), indicating that this MilM variant is catalytically inactive.

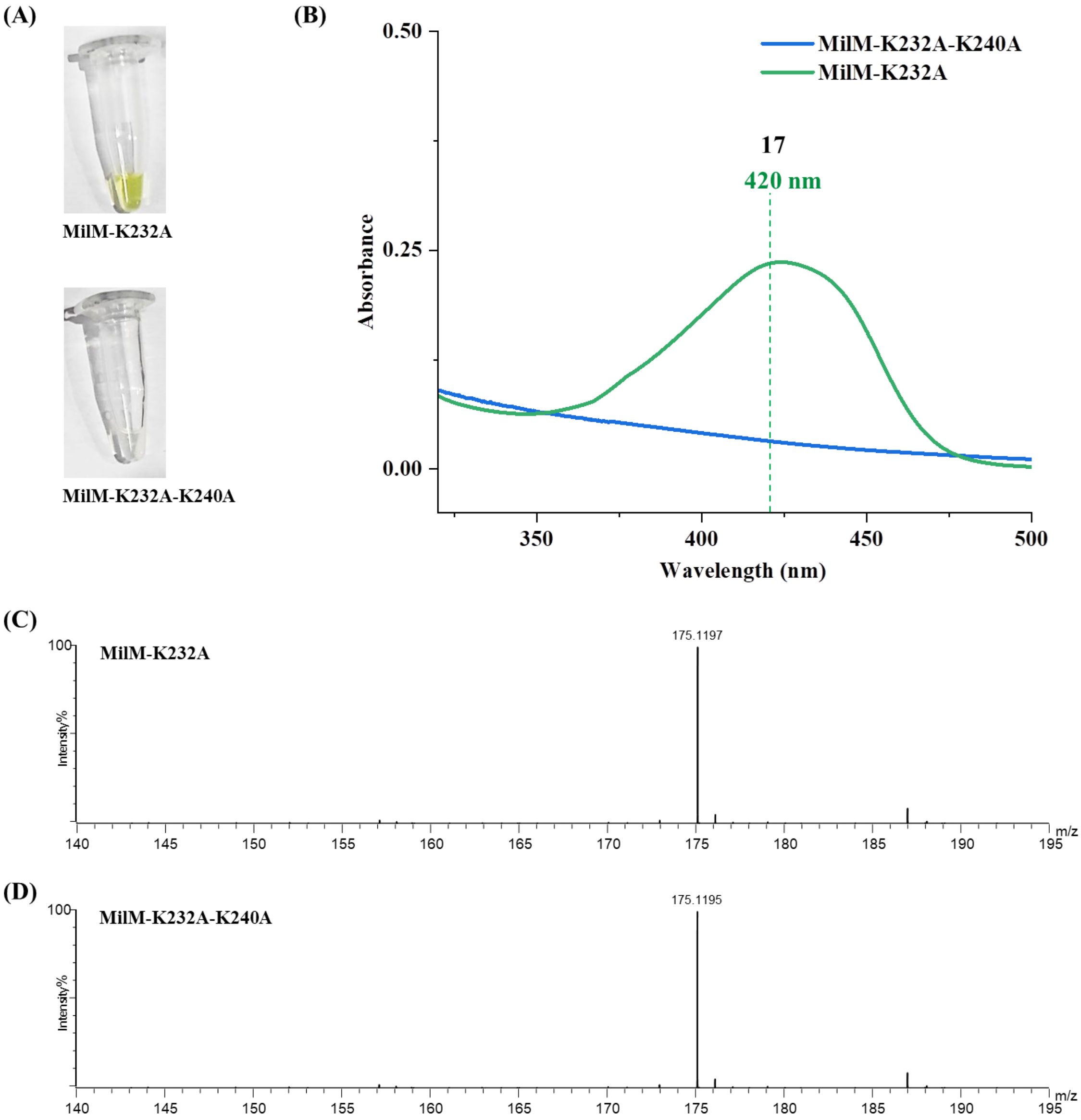

**Figure S6: DNS-Cl derivatization of L-Arg and analysis of MilM reaction mixtures.**

**(A)** Reaction scheme showing derivatization of L-Arg with dansyl chloride (DNS-Cl) to generate DNS-Arg.

**(B)** HPLC chromatograms of DNS-derivatized samples monitored at 254 nm: (i) Derivatized DNS-Arg showing a peak at retention time (Rt) ~3.5 min (black). (ii) MilM reaction (L-Arg + α-KG) derivatized with DNS-Cl post-reaction, showing complete disappearance of the DNS-Arg peak, indicating complete consumption of L-Arg (red). (iii) Enzyme control reaction where no MilM was added showing formation of DNS-Arg, indicating that no consumption of L-Arg occurs in the absence of MilM (blue). (iv) MilM reaction performed without α-KG, also showing loss of DNS-Arg peak (green). Together, these results demonstrate that MilM consumes L-Arg as substrate independently of α-KG, confirming that MilM does not function as an aminotransferase.

**(C)** LCMS spectrum of DNS-Arg showing a molecular ion peak at *m/z* 408.1710 ($[M+H]^+$, calc. 408.1700), confirming the expected derivatized Arg formation at Rt 3.5 min in HPLC chromatogram.

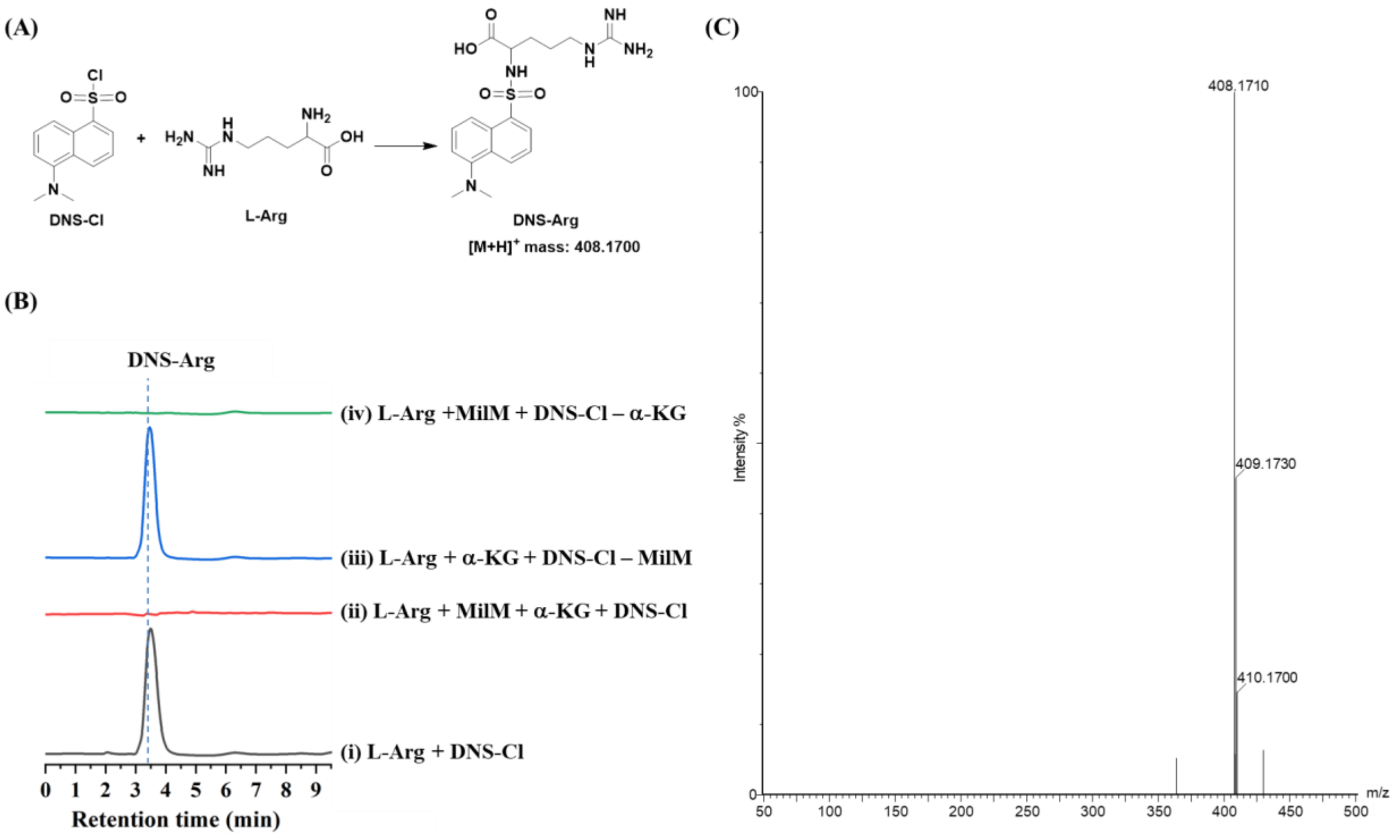

**Figure S7: LCMS analysis of the MilM-catalyzed reaction with L-Arg as substrate.**

**(A)** LC-MS profile of the MilM reaction with L-Arg (**9**) shows the formation of the oxo product **13** (*m/z* 174.0874 $[M+H]^+$, calc. *m/z* 174.0873) and the hydroxy product **10** (*m/z* 190.0825 $[M+H]^+$, calc. *m/z* 190.0822), along with the non-enzymatic products of **13** and **10**, **14** (*m/z* 146.0923 $[M+H]^+$, calc. *m/z* 146.0894) and **15** (*m/z* 162.0874 $[M+H]^+$, calc. *m/z* 162.0873), respectively.

**(B)** MilM catalyzes the oxidation of L-Arg using molecular oxygen and water to yield either 5-guanidino-2-oxovaleric acid (**13**) or 5-guanidino-4-hydroxy-2-oxovalric acid (**10**). Subsequent non-enzymatic reactions of these intermediates with hydrogen peroxide, a by-product released during catalysis, lead to products **14** and **15**, respectively (dashed box).[4]

**(C)** Mechanistic representation of $H_2O_2$ reaction with the products **13** and **10** to give **14** and **15**, respectively.

**(D)** In the presence of catalase, LCMS profile of the MilM reaction shows only the masses corresponding to **10** and **13**, indicating that catalase scavenges the $H_2O_2$ generated as a by-product during catalysis, which is responsible for the non-enzymatic conversion of **10** and **13** to produce **15** and **14**, respectively.

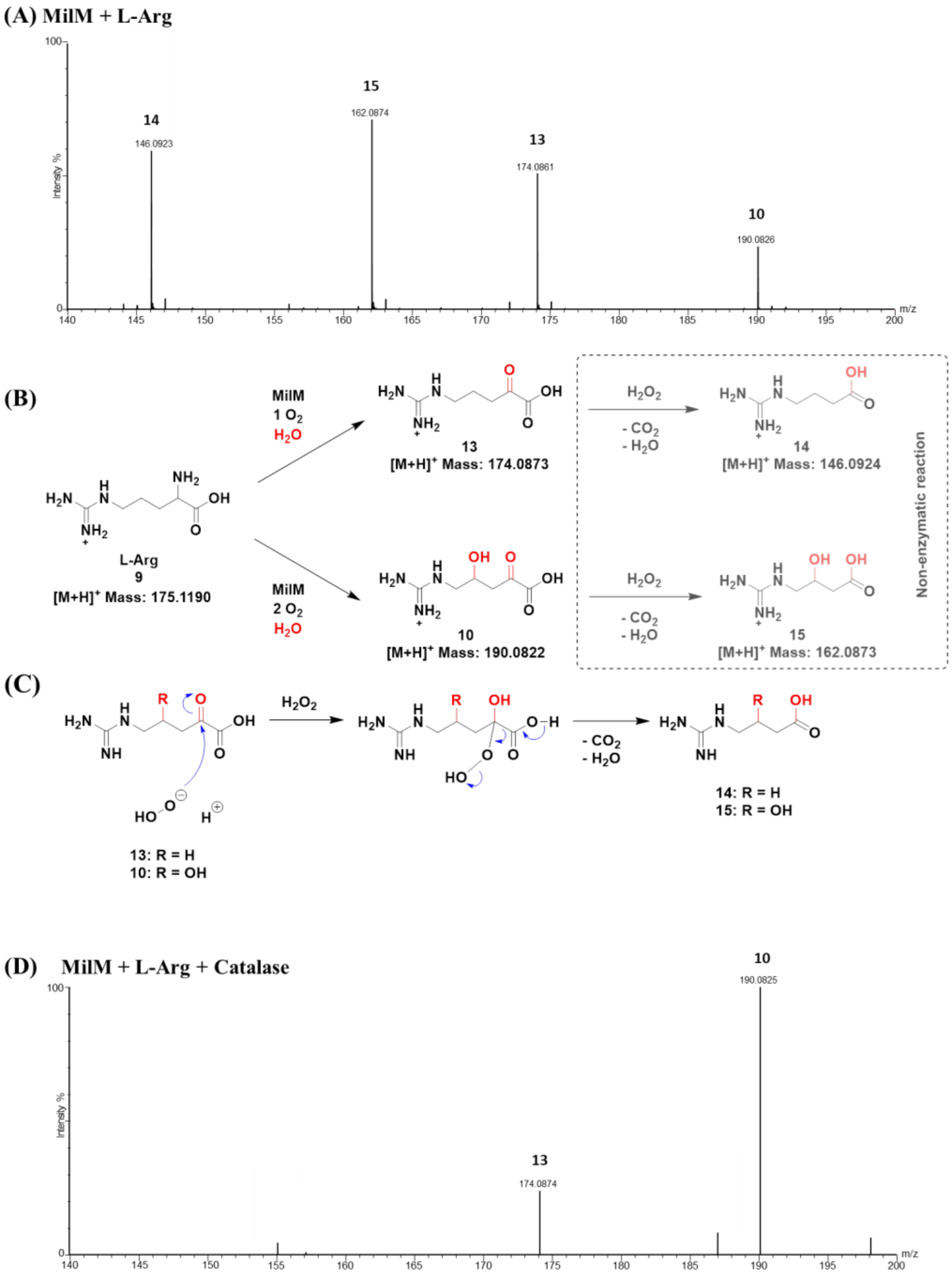

**Figure S8: $^1$H-NMR analysis of the MilM assay.** $^1$H-NMR spectra showing the conversion of L-Arg (green spectrum) to oxo- and hydroxy- products by MilM (blue spectrum). The reaction mixture of MilM with L-Arg (**9**) (without catalase, top spectrum) displays new resonance signals corresponding to hydroxylated (**10**) and oxo (**13**) products, which are absent in the L-Arg substrate (bottom spectrum). The expanded insets highlight the characteristic peaks for protons, $H^f$ (~ δ 3.95 - 4.03 ppm, multiplet, 1H) and $H^{f'}$ (~ δ 1.6 - 1.7 ppm, multiplet, 2H) associated with the hydroxylated (**10**) and oxo (**13**) products, respectively. The other unassigned peaks may be arising from the non-enzymatic products, **14** and **15** **(Figure S7A)** and the peaks denoted by * marks are due to glycerol contamination from the protein sample. Proton assignments for each metabolite are indicated alphabetically in red on respective molecular structures.

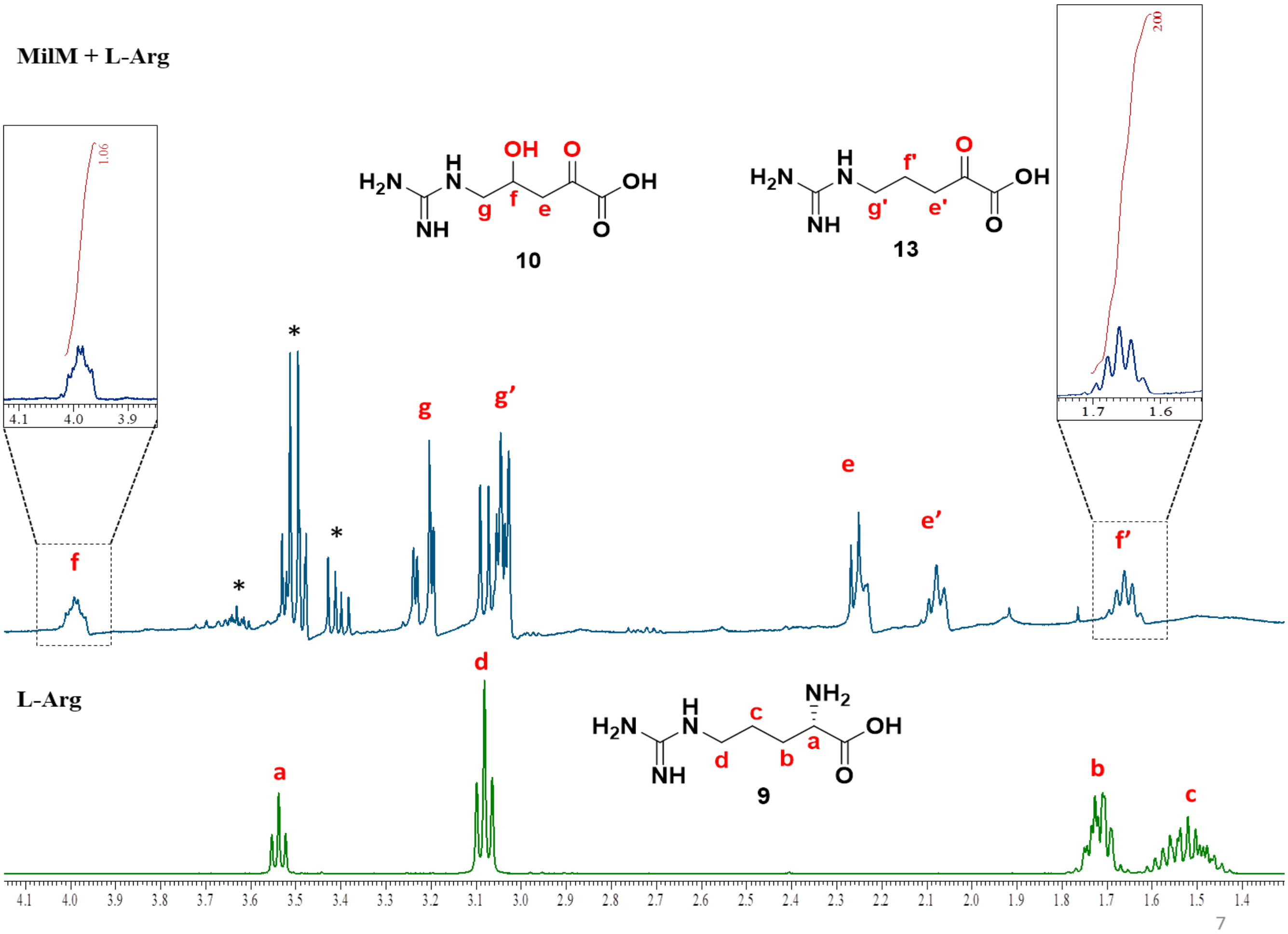

**Figure S9: UV-Vis spectroscopic investigation of the MilM reaction showing the time-dependent formation of quinonoid intermediates during the MilM-catalyzed oxidation of L-Arg.** The internal aldimine peak at 420 nm is exhibited by MilM in its resting state due to imine formation with PLP. But, upon substrate L-Arg addition, distinct absorption bands were observed at 425 nm (external aldimine with substrate L-Arg), 516 nm (quinonoid intermediate Q1), and 565 nm (quinonoid intermediate Q2), corresponding to characteristic PLP-dependent enzyme intermediates as per our proposed mechanism (**Figure 3**). The gradual emergence and decay of these peaks over time indicate sequential formation and interconversion of these PLP-substrate intermediates during the MilM catalytic cycle. Blank (in black) refers to the HEPES buffer used for this assay.

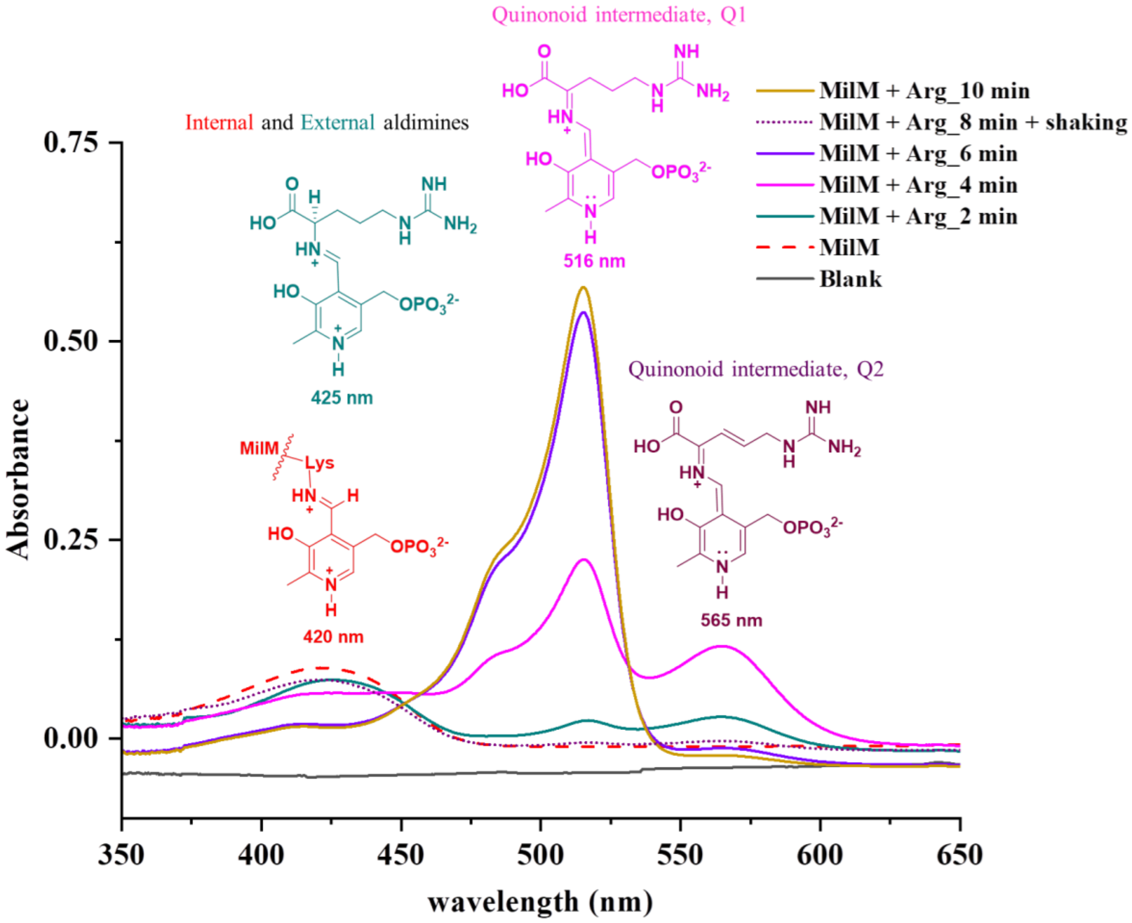

**Figure S10: MilM assay with L-Arg under anaerobic conditions.**

**(A)** UV-Vis spectra of MilM reaction recorded under anaerobic conditions in the presence of L-Arg at different time points (6, 10, and 15 min). The characteristic absorbance of the quinonoid intermediate (Q1, **19**) near 515 nm is retained over time, and no absorbance at 565 nm for Q2 (**24**) intermediate was observed, indicating that after the formation of first PLP-amino acid intermediate (Q1), the reaction does not proceed in the absence of oxygen (**Figure 3**).

**(B)** LC-MS analysis of the MilM reaction mixture under anaerobic conditions showing only the peak corresponding to unreacted L-Arg substrate (*m/z* = 175.1182 $[M + H]^+$, calc. 175.1190), confirming that MilM-mediated turnover requires molecular oxygen for subsequent product formation.

**(A)**

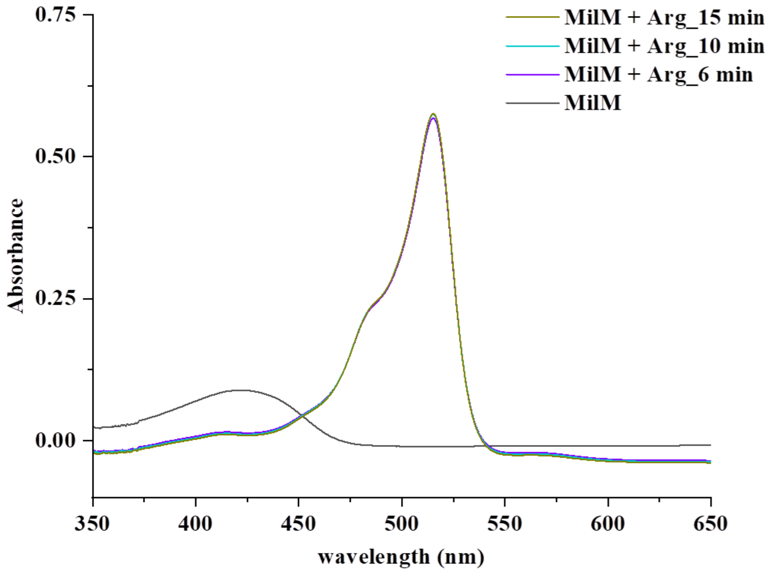


**(B) MilM + L-Arg in Anaerobic Chamber**

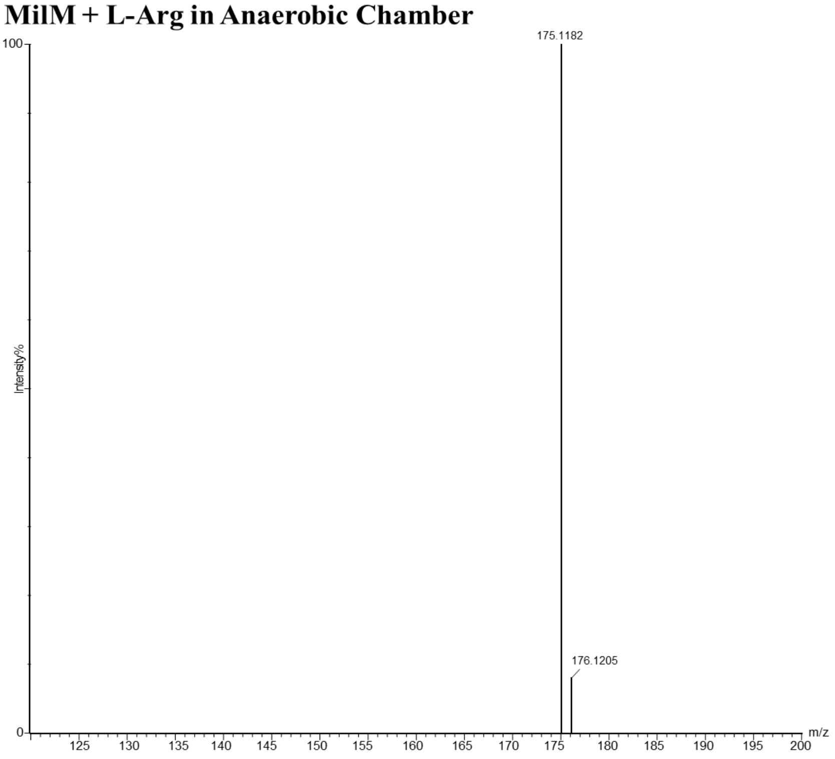

**Figure S11: MilM reaction in the presence of superoxide dismutase (SOD).**

**(A)** LC-MS spectra of MilM reaction mixtures containing L-Arg (**9**) and varying concentrations of SOD (0.5-10 mg/mL) in the presence of catalase. The characteristic product ion corresponding to **10** (*m/z* = 190.0824 [M + H]$^+$) and **13** (*m/z* = 174.0868 [M + H]$^+$) progressively diminishes with increasing SOD concentration, while the mass corresponding to unreacted L-Arg (*m/z* = 175.1186 [M + H]$^+$) increases. This is consistent with the fact that SOD quenches the superoxide radical anion (by converting it to $H_2O_2$ and $O_2$, panel **(B)**)[5], which is necessary for a critical hydrogen abstraction step in our proposed mechanism (**Figure 3**), leading to impairment of the reaction. These data support the possible intermediacy of the superoxide radical anion implicated in the catalytic mechanism of MilM.

**(A)**

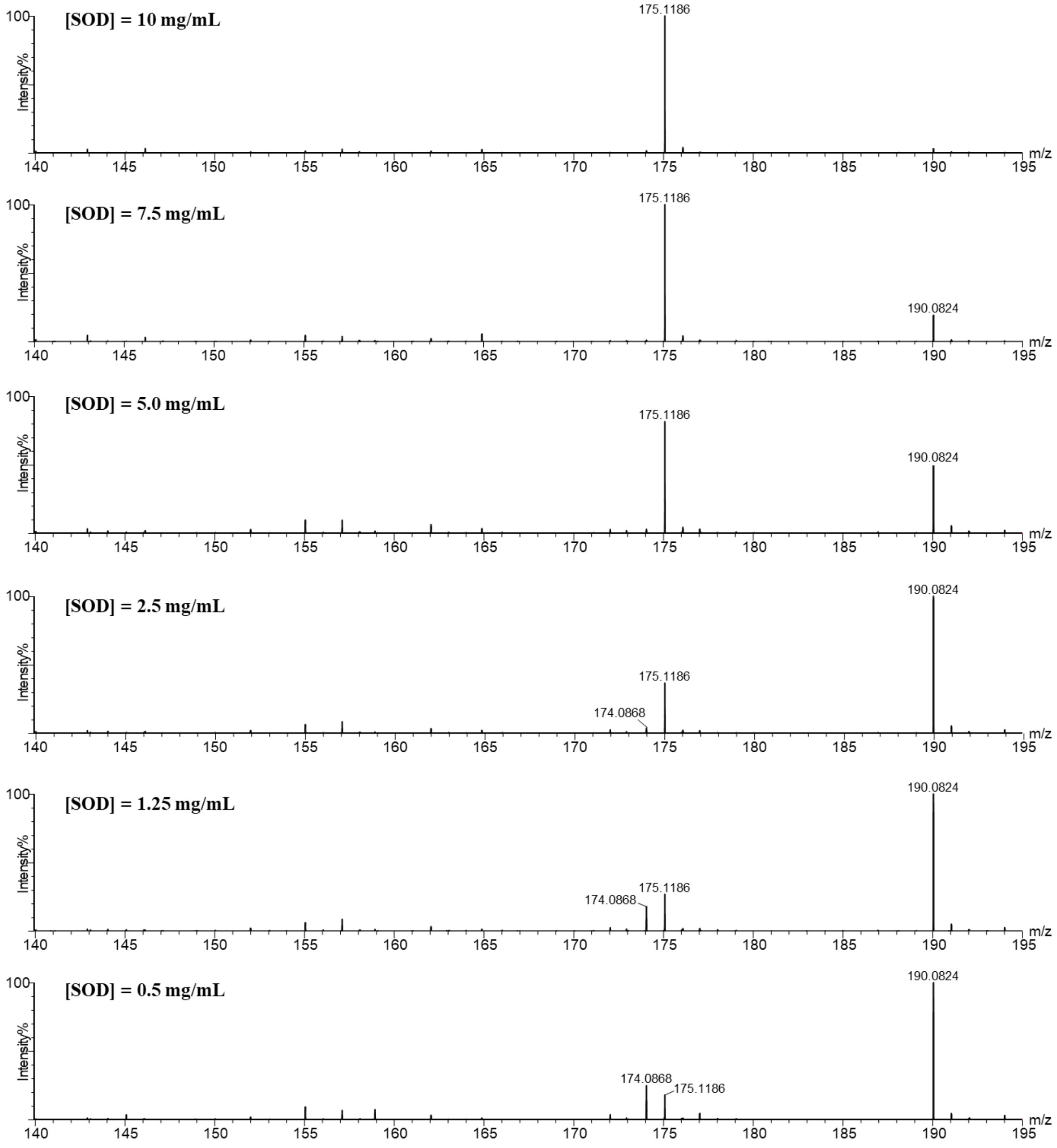


**(B)**

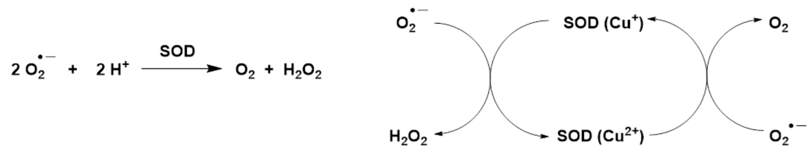

**Figure S12: Quantification of residual L-Arg in the MilM reaction in the presence of varying amounts of SOD using DNS-Cl derivatization.**
**(A)** HPLC chromatograms of DNS-derivatized MilM reaction mixtures in the presence varying concentrations of SOD (0.5-10 mg/mL), demonstrating reduced L-Arg consumption upon SOD addition, confirming the role of superoxide radical anion in the MilM-catalyzed reaction.
**(B)** HPLC chromatograms of standard L-Arg derivatization with DNS-Cl, showing formation of DNS-Arg standard at Rt = 3.5 min.
**(C)** Standard calibration curve for DNS-Arg used for quantification.

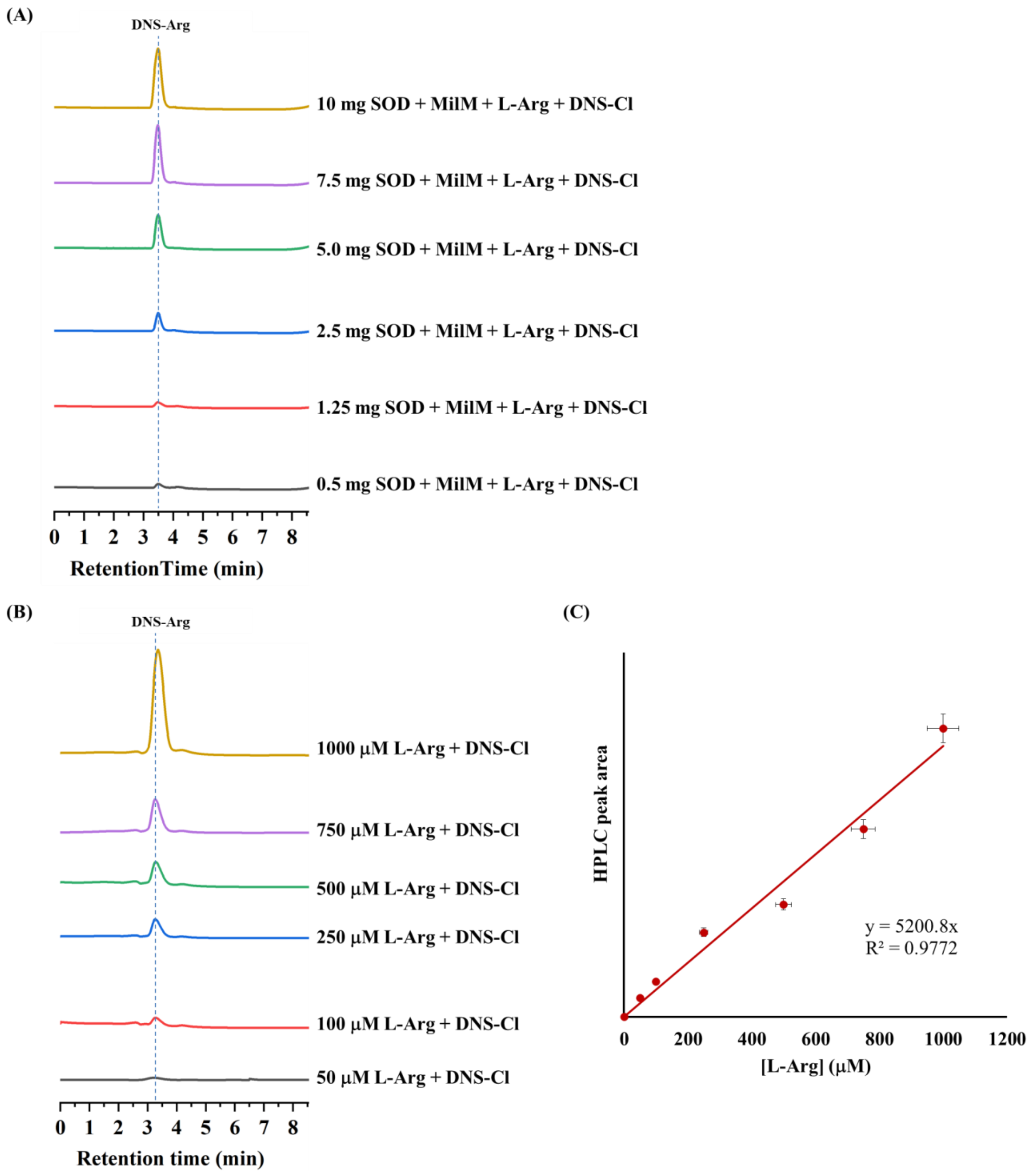

**Figure S13: pH-dependent product formation catalyzed by MilM.** LCMS analysis of MilM-catalyzed reactions performed in the presence of catalase across a pH range of 5.5-9.0 (50 mM HEPES buffer) displayed the hydroxylated product (**10**, *m/z* = 190.0825 [M + H]$^+$) decreases over pH 8.0 demonstrating the importance of protonation state of the catalytic base, His31 on hydroxylation (**Figure 3**).

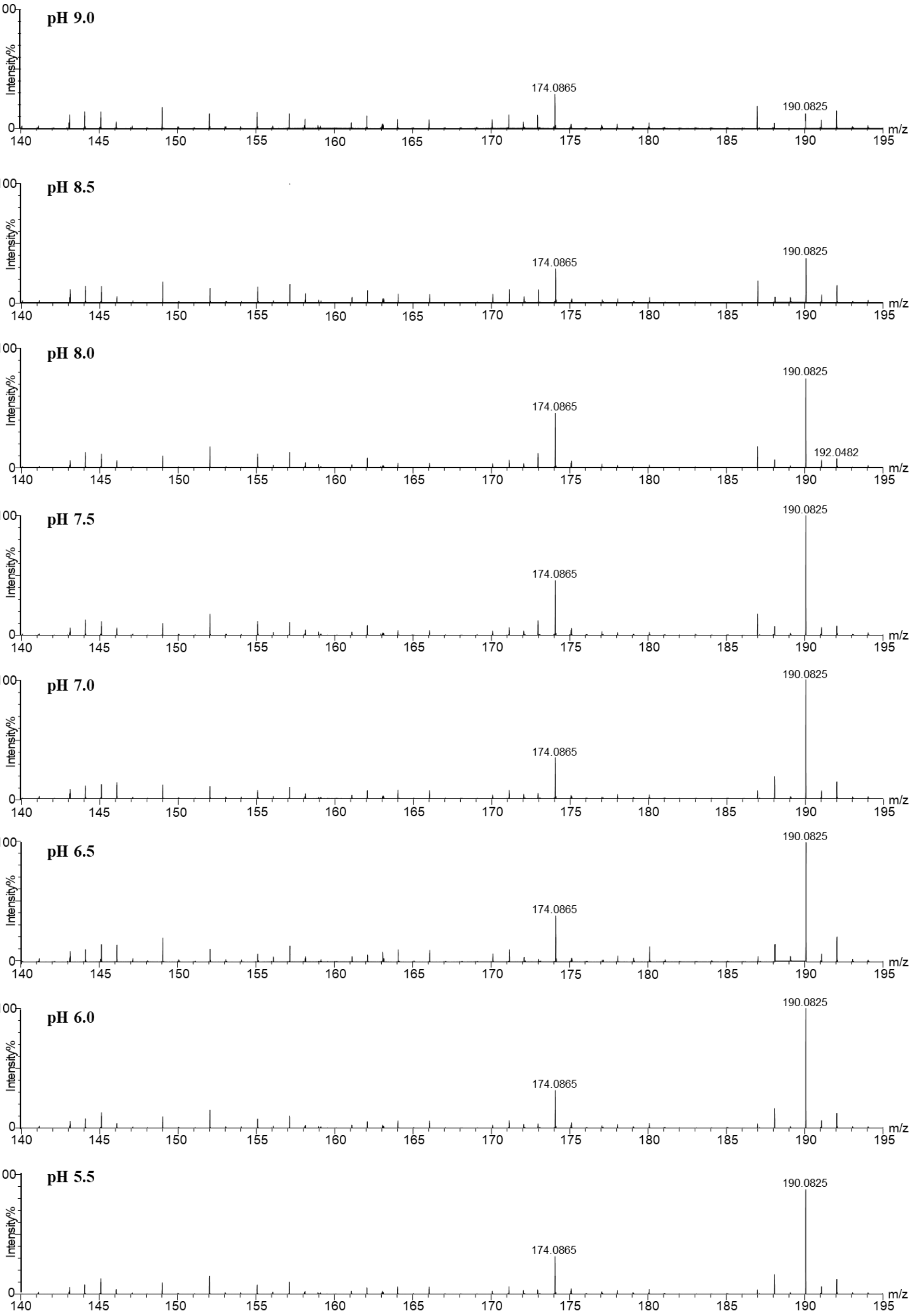

**Figure S14: LCMS analysis of MilM-His31Ala variant.**

(**A**) LCMS chromatogram of as-isolated MilM-H31A showing this variant is catalytically inactive as no product formation was observed and only the unreacted substrate L-Arg (**9**) mass (*m/z* = 175.1193 $[M + H]^+$) was detected.

(**B**) LCMS chromatogram of the reaction mixture of MilM-H31A variant reconstituted in the presence of externally added PLP showing a little formation of only the oxo product (**13**, *m/z* = 174.0862 $[M + H]^+$, ~10% product conversion) and no formation of the hydroxylated product (**10**), which confirms that His31 is a critical residue for hydroxylation.

**(A)**

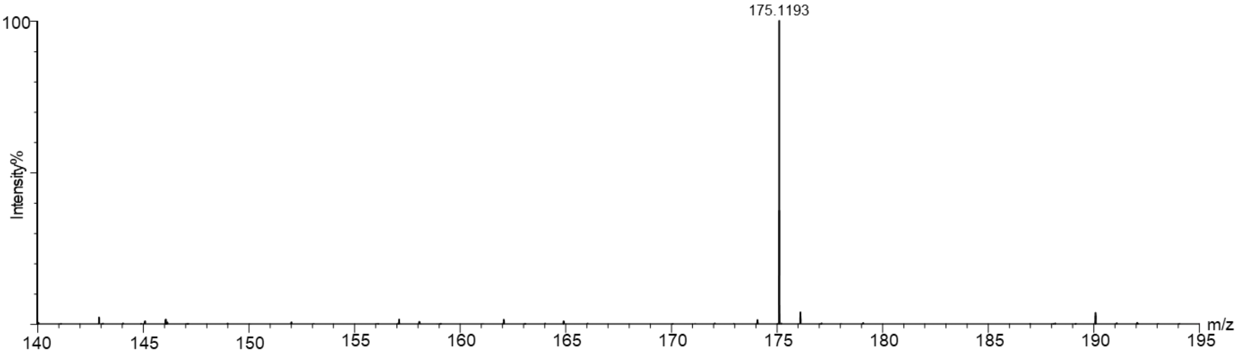


**(B)**

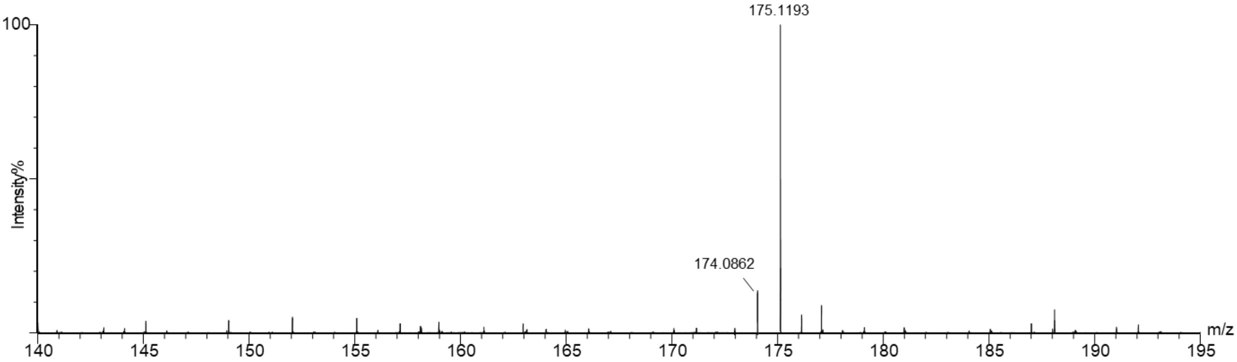

**Figure S15: Detection of $H_2O_2$ and $NH_3$ as by-products in MilM-catalyzed reaction.**

**(A) ABTS-HRP assay to detect $H_2O_2$:** Schematic representation of the oxidation of ABTS (colorless / light green, $\lambda_{max}$ = 340 nm) to its radical cation ABTS•$^{+}$ (bluish green, $\lambda_{max}$ = 418 nm) by $H_2O_2$ in the presence of HRP.[6]

**(B)** 96-well plate image of ABTS-HRP assay in 50 mM HEPES pH 7.5 having 10 mM ABTS and 10 ng HRP, where **1** contain different L-Arg concentrations (1000-10 μM) with MilM-WT.

**(D) Glutamate dehydrogenase (GDH) assay to detect $NH_3$:** Schematic representation of conversion of α-KG to L-Glu by $NH_3$ in the presence of GDH and NADH ($\lambda_{max}$ = 340 nm).[7]

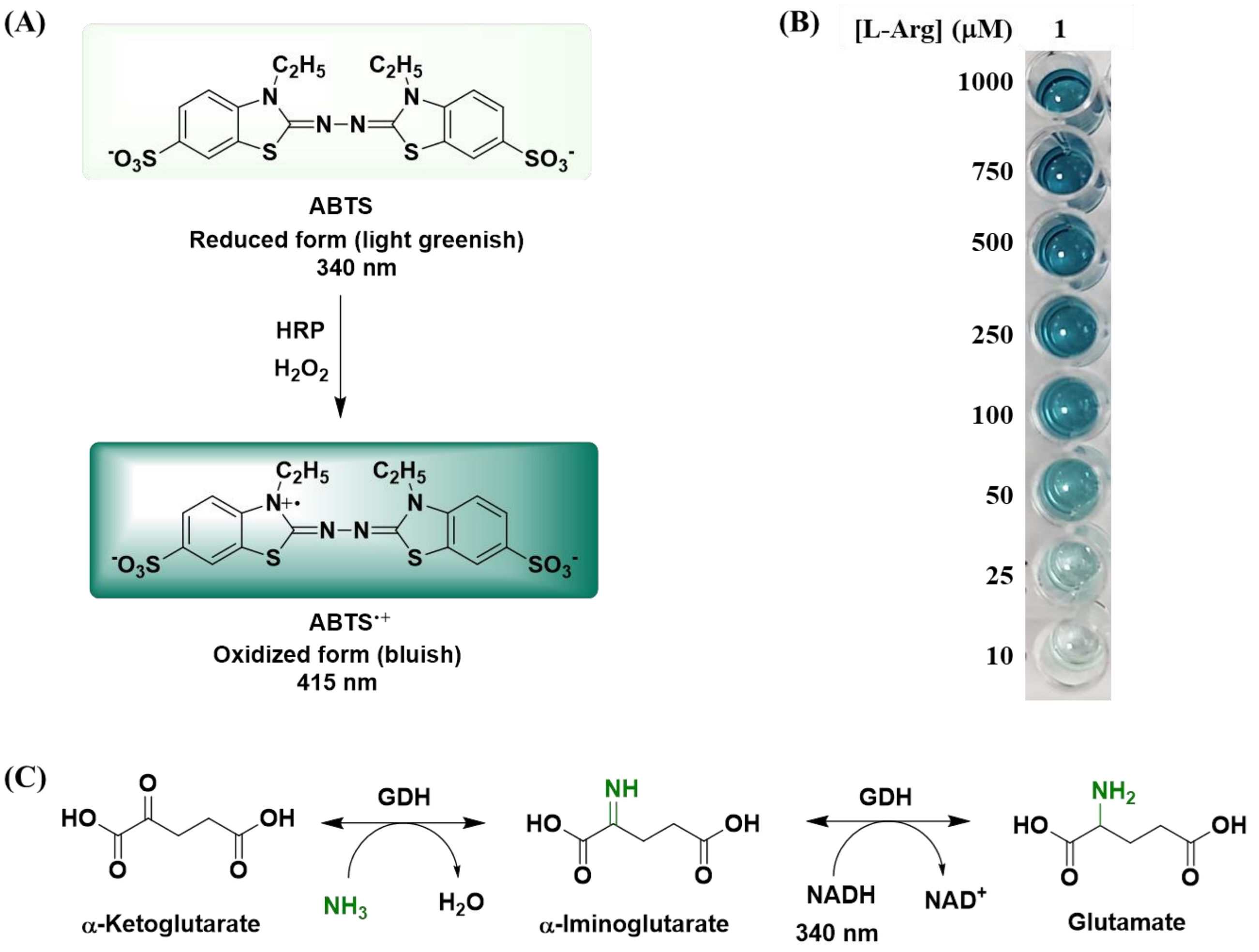

**Figure S16: DimPlot analysis of residues stabilizing the MilM dimer interface.** Carbon atoms of interacting residues from chain A and chain B are labeled in green and red, respectively. The interface is stabilized by a network of hydrophobic contacts, hydrogen bonds, and polar (hydrophilic) interactions between the two subunits.

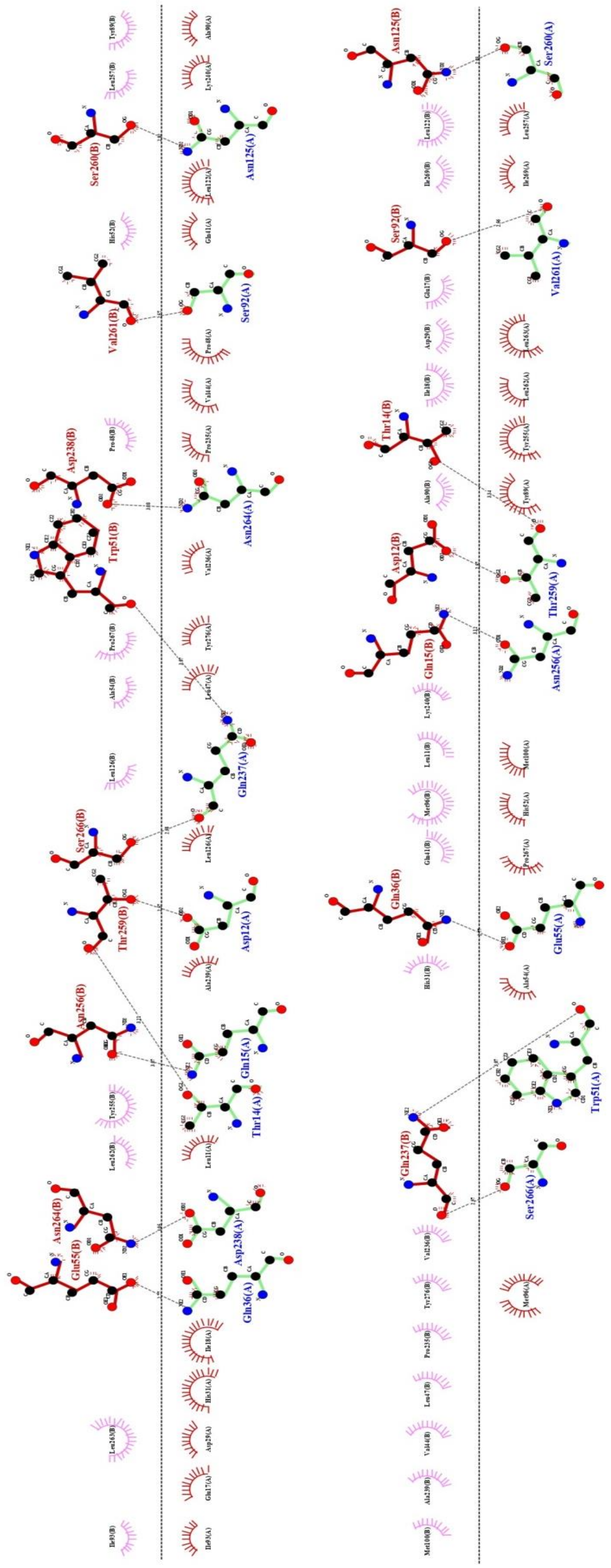

**Figure S17: Topology of MilM.**

**(A)** Secondary structure prediction of MilM generated using the PDBsum server.

**(B)** Topology diagram of the MilM AlphaFold model, illustrating the central 13-stranded β-sheet (pink) surrounded by α-helices (red), generated using TopDraw based on PDBsum-derived topology analysis.

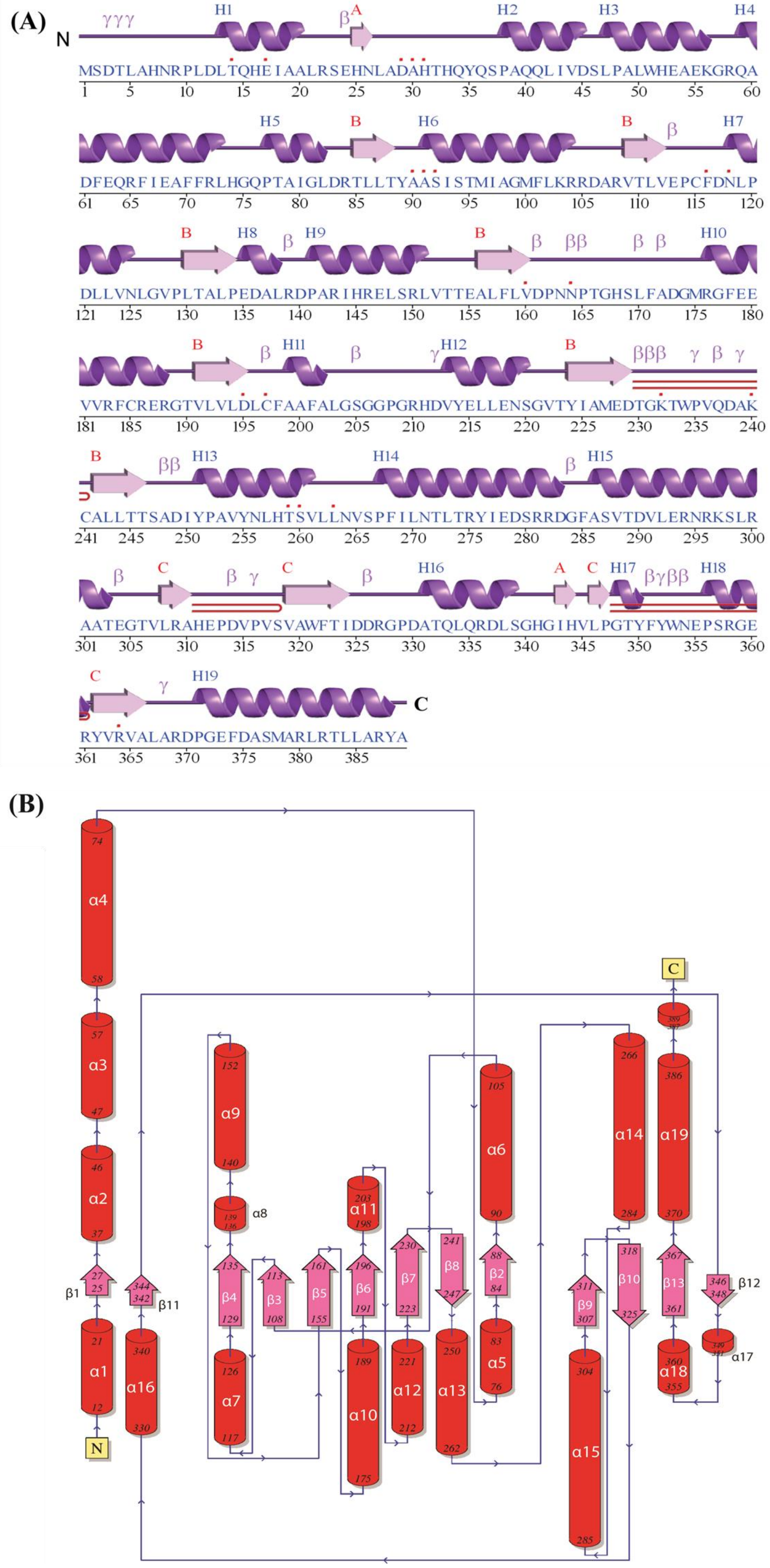

**Figure S18: Sequence alignment of MilM homologs showing conservation of residues at the dimer interface.** Eleven sequences were obtained from NCBI BLAST analysis[8] using *S. rimofaciens* as the input sequence and the alignment was generated using Clustal Omega.[3] The details of all the sequences are given below. Conserved motifs for dimer interface residues (in yellow) are highlighted in reference to **Table S3**. The numbing of amino acids in respect to MilM are mentioned in red.

- MilM: AFD20753.1 aspartate/tyrosine/aromatic aminotransferase [*Streptomyces rimofaciens*]
- RohP: G8WNKC $O_2$-, PLP-dependent L-Arginine hydroxylase [*Streptantibioticus cattleyicolor*]
- MppP: A0A0X1KHF5 PLP-dependent L-arginine hydroxylase [*Streptomyces wadayamensis*]
- Ind4: A0A0D4BS17 type I PLP-dependent aspartate aminotransferase-like [*Streptomyces griseus subsp. griseus*]
- WP_399337027.1 aminotransferase class I/II-fold pyridoxal phosphate-dependent enzyme [*Streptomyces* sp. NPDC050560]
- WP_073229495.1 MULTISPECIES: aminotransferase class I/II-fold pyridoxal phosphate-dependent enzyme [unclassified *Streptomyces*]
- WP_384049659.1 aminotransferase class I/II-fold pyridoxal phosphate-dependent enzyme [*Streptomyces* sp. NPDC057565]
- WP_149851715.1 aminotransferase class I/II-fold pyridoxal phosphate-dependent enzyme [*Solihabitans fulvus*]
- WP_051969389.1 pyridoxal phosphate-dependent aminotransferase [*Kitasatospora azatica*]
- WP_116214779.1 aminotransferase class I/II-fold pyridoxal phosphate-dependent enzyme [*Streptomyces olivoreticuli*]
- WP_229328800.1 aminotransferase class I/II-fold pyridoxal phosphate-dependent enzyme [*Streptomyces* sp. UNOC14_S4]
- MGH3938013.1 pyridoxal phosphate-dependent aminotransferase [*Pseudonocardiaceae bacterium*]
- MGB6162345.1 pyridoxal phosphate-dependent aminotransferase [*Pseudonocardiaceae bacterium*]

```
                                                  12 15                   36
MilM                -------------------MSDTLAHNRPLDLTQHEIAALRSEHNLADAHTHQYQSPAQQ     41
RohP                ----------------MHPQATPAPGAPLLDLTQHEIQALTMKYNLADAHTHQRQSASQQ     44
MppP                ---------------------MTTQPQLKENLTQWEYLALNSELNIADGHARQALSPGQQ     39
Ind4                -------------------------MERFNNLTRYEHVGINQAINLADGHAHQGQNSTQQ     35
WP_399337027.1      -------------------MSPSALSLSAADLTQHEIRALRSEYNLADAHTHQAQSPSQR     41
WP_073229495.1      -------------------MPGTTLTVSAADLTQHEIRALRSEHNLADAHTHQSQSPGQR     41
WP_384049659.1      -------------------MSGTALSVSAVDLTQHEIRALRSEHNLADAHTHQSQSSGQR     41
WP_149851715.1      MHGEADTRENPLLTASGQDGAVIPDVAGFVDLTQHEIQALKTRFNLADAHTHQHQSASQD     60
WP_051969389.1      MNPNQS---------QTAIDSAVDPAVDLLDLTQHEIEALRTRYNLADAHTHQQQSPRQR     51
WP_116214779.1      MNHS-G---------TRTSSPENLTDGALVDLTQHEIQALRTEFNLADAHTHQRQSASQR     50
WP_229328800.1      MTTS-S---------IRTGSPETRTGGALVDLTQHEIEALRTEFNLADAHTHQRQSASQQ     50
MGH3938013.1        MYRP-S---------LTASSSFNFDRETLVDLTQHEIQALRTEFNLADAHTHQRQSLSQR     50
MGB6162345.1        MRYS-S---------IAIEPMLGLDRGTLVDLTQHEIQALRTEFNLADAHTHQRQSSSQR     50
                                                  :**: *  .:    *:**.*::*  .  *

                             51  55                                     92
MilM                LIVDSLPALWHEAEKGRQADFEQRFIEAFFRLHGQPTAIG-LDRTLLTYAASISTMIAGM     100
RohP                SIVSRLPQLWYEAEEGLQATYEKRFTEAFFQLHRQPTALV-KNKTMLSYAASISTMVAGM     103
MppP                KIVNELPVLWAESEQRPVQQIESEAHQAYFTLLGQHGYPAEPGRVLSCYSSSVSMEILAR     99
Ind4                AIVADLPNIFEAAENALQATTEREFQRAFYTLAGQHSAVD-HPRTMLCYSASLSTDLVAT     94
WP_399337027.1      GIVESLPRLWYESEAGRQGEFERRFLDAFFRFRGQRTAPG-LGRTLLTYAASVSTVIAGM     100
WP_073229495.1      QIVASLPQLWYEAEAGKQSAFEERFIDTFFRFRGQHTALG-LNRTLLTYAASVSTVVAGM     100
WP_384049659.1      QIVASLPQLWYESEAGKQSTFEQRFIDTFFRFRGQHTALG-LDRTLLTYAASISTVIAGM     100
```

```
WP_149851715.1      KIVSKLPDLWYEAENGQQAYYEQRFIKAFFRLHEQPTAAG-LNKTMLSYAASISTMVAAM      119
WP_051969389.1      EIVASLPKLWYQAEEGLQAHYEQRFTEAFFRLHRQPTALA-RNKTMLSYAASISTMVAGM      110
WP_116214779.1      QIVERLPQLWYEAEEGLQETYEQRFVDAFFRLHRQPTALA-RKKTMLSYAASISTMVVGM      109
WP_229328800.1      HIVARLPQLWYEAEEGLQETYEQRFTDAFFRLHRQPTALA-RKKTMLSYAASISTMVVGM      109
MGH3938013.1        KIVEQLPKLWYEAEEGLQETYEHRFTEAFFRLHGQPTALV-RGKTLLSYAASISTMVAAM      109
MGB6162345.1        KIIEQLPMLWYEAEQGLQETYEHRFTEAFFRLRGQPTVLA-KGKTLLSYAASISTMVAAM      109
                     *:  ** ::  :*       * .   ::: :  *        :.:  *::*:*  : .

                      104           118    125
MilM                FLKRRDARVTLVEPCFDNLPDLLVNLGVPLTALPEDALRDPARIHRELSRLVTTEALFLV      160
RohP                FLKKERLAVTLIEPCFDNLYDVLANMDVPLYPIDESVFYDVDRIYPELERRVRTDALFLV      163
MppP                SLSASVDRVALVHPTFDNIADLLRGNGLDLVPVEEDALHGADLSAELL---SSVGCVFVT      156
Ind4                FLASRNLTVGLLQPCFDNLATILRRRQVKVVPLGEEQFRPES--LDRTFAELNTDAVFLT      152
WP_399337027.1      YLKRQGARVSLIEPCFDNLRDLLVNLDVELVPLPEEALASADTVYDRLARQPRTEALFLV      160
WP_073229495.1      YLKRRGARVSLIEPCFDNLRDLLVNLDIELVSLPEQVLASADTVYDQLAQQPATEAVFLV      160
WP_384049659.1      YLKRRGARVSLIEPCFDNLRDLLVNLDIELVPLPEQVLSSADTVYDRLAQQPRTEVVFLV      160
WP_149851715.1      FLSQRNMSVTLVEPCFDNLVDLLRNMQVELNAVDESHLADPASIYDNLVREVKTDALFLV      179
WP_051969389.1      FLKQRRMAVTLIEPCFDNLHDVLANLGVAAYPLDESALHDPDRIYSELKRRVRTDALFLV      170
WP_116214779.1      YLKRRRMAVTLIEPCFDNLYDVLANMGVPLHPVDESAFADPDRIYQELKRRVRTDALFLV      169
WP_229328800.1      YLKQQRMAVTLIEPCFDNLYDVLANMGVPLYPIDESALADPDRIYDELRRRVRTDALFLV      169
MGH3938013.1        FLKQQRMAVTLIEPCFDNLHDVLANMGVPLYPIDESVFTDPDLIYSELKRRVRTDALFLV      169
MGB6162345.1        FLKQHRMAVTLVEPCFDNLHDVLANVGVPTYPIGESALTNPNMIYSELERHVRTDALFLV      169
                     *      * *:.* ***:  :*    :    : *. :               .  :*:.

MilM                DPNNPTGHSLFADGMRGFEEVVRFCRERGTVLVLDLCFAAFALGSGGPGRHDVYELLENS      220
RohP                DPNNPTGFSLLRHGRKGFEEVVRFCKDHDKLLLIDFCFASFTLFEPELARFDMYELLENS      223
MppP                TPNNPTGRVLA---EERLRRLAEQCAEHGTVLALDTSFRGFDA----AAHYDHYAVLQEA      209
Ind4                LPNNPTGFHLD---QEQFRRVIDLCAEHQKILIVDCTFRFFAP----TPFWDQYAQLEES      205
WP_399337027.1      DPNNPTGHSLLAGGRGTFEEVVRFCRDNGRLLVLDLCFAAFALNSAG-GRTDVYEILEAS      219
WP_073229495.1      DPNNPTGHSLLTGGRRAFEEVVRFCRENNRLLVLDLCFAAFALDSPG-GRIDVYEILETS      219
WP_384049659.1      DPNNPTGHSLLTGGRRIFEEVVRFCRENDRLLVLDLCFAAFALGSPG-GRIDVYEILEAS      219
WP_149851715.1      DPNNPTGFSLLKDGRRGFEEVVRFCVDHNKVLVIDQCFAAFALADRAVGRFDIYQLLESS      239
WP_051969389.1      DPNNPTGFSLLQFGRRGFEEVVRFCKDHHKVLVIDFCFASFTLFDSEATRFDVYELLESS      230
WP_116214779.1      DPNNPTGFSLLKHGRKGFEEVVRFCKDHGKLLVIDFCFASFTLFDPDVARFDMYELLENS      229
WP_229328800.1      DPNNPTGFSLLKHGRRGFEEVVRFCKDHDKLLVIDFCFASFTLFDPAVARFDMYELLENS      229
MGH3938013.1        DPNNPTGYSLLAHGRKGFEEVVRFCRDHGKLLVIDFCFAAFTLFDSKHARFDIYELLENS      229
MGB6162345.1        DPNNPTGYSLLAHGRKGFEEVVRFCRDHGKLLLIDFCFAAFTLFDSKLARFDVYELLENS      229
                     ******  *       :..:   * :.  :* :*  *  *          * *  *: :

                                   237               256 259  264 266
MilM                GVTYIAMEDTGKTWPVQDAKCALLTTSADIYPAVYNLHTSVLLNVSPFILNTLTRYIEDS      280
RohP                GVRYLAIEDTGKTWPVQDAKCALITASDDIWETVYNLHTSVLLNVSPFVLNMLTQYVRDS      283
MppP                GCRWVVIEDTGKLWPTLDLKAGLLVFSEDIGLPVEKIYSDILLGVSPLILALIREFSRDA      269
Ind4                GISYVVVEDTGKTWPTQDLKCSILAVSTDLHESVLELHNDILLNVSPFILQLLTAYLRDT      265
WP_399337027.1      GVSYIAMEDTGKTWPVQDAKCALITASADLYPHLYNIHTSVLLNVSPFTLNVLTHYVEDS      279
WP_073229495.1      GVSYIAMEDTGKTWPVQDAKCALITASADIYPDLYNVHTSVLLNVSPFTLNVLTRYIEDS      279
WP_384049659.1      GVSYIAMEDTGKTWPVQDAKCALITASADIYPELYNIHTSVLLNVSPFTLNVLTQYIKDS      279
WP_149851715.1      GVTYMAMEDTGKTWPVQDAKCAMITASDDIQGEVYNLHTSVLLNVSPFVLNFLAHYIEDS      299
WP_051969389.1      GVTYLAIEDTGKTWPVQDAKCAMITSSDDIREAVYNLHTSVLLNVSPFTLNMLAHYLEDA      290
WP_116214779.1      GVSYLAIEDTGKTWPVQDAKCAMLTVSDDIHEAVYNLHTSVLLNVSPFVLNMLAEYIEDS      289
WP_229328800.1      GVRYMAIEDTGKTWPVQDAKCAMLTVSDDIHEAVYNLHTSVLLNVSPFVLNMLTEYIEDS      289
MGH3938013.1        GVTYMAIEDTGKTWPVQDAKCAMITVSDDIRKAIYNIHTSVLLNVSPFVLNMLTRYIEDS      289
MGB6162345.1        GVTYMAIEDTGKTWPVQDAKCAMITVSDDIREMVYNIHTSVLLNVSPFVLNMLTRYIDDS      289
                    *  ::.:***** **. * *..::. * *:   : :::..:**.***: *  :  :  *:
```

```
MilM                RRDGFA-SVTDVLERNRKSLRAATEGTV-LRAHEPDVPVSVAWFTIDDRGPDATQLQRDL      338
RohP                AADRLA-SVREVLTRNRECARKTLDGSI-LEYQEPVVKVSVAWFRVDHPELTATDVHRLL      341
MppP                ADGGLA-DLHAFILHNRSVVRRALAGVEGVSFPDPESRSSVE--RVAFAGRTGTEVWEEL      326
Ind4                ERNGLQQTIWSVVRANRISLRKALEGTV-LVPAHPESTISVEWVRIDHPSLRSDDLVEML      324
WP_399337027.1      LADGFA-SVADVLSANRELARELTKGTV-LRHQEPAVPVSVAWFELDDRAPTATELQREL      337
WP_073229495.1      MADGFA-SVADVLSENRSVARELTRGTV-LQHQEPAVPVSVAWFRIADGAPRATELQQEL      337
WP_384049659.1      VADGFA-SVSDVLSTNRALARELTRGTV-LHHQEPAVPVSVAWFELADSAPSATELQQEL      337
WP_149851715.1      IEDGFA-SVRDVITENRKLCRESLDGDV-LEYQEPSVNVSVAWFRIKPAGLTATKLQEEL      357
WP_051969389.1      RHEQLA-SVRDVLNRNRDQVRKTLDGTI-LQYQEPAVKVSVAWAKVNHPLLTATDLQRLL      348
WP_116214779.1      IEDDLS-SVRDVLTRNRETARKALDGTI-LEYQEPVVNVSVAWFRIDHPELTATELQREL      347
WP_229328800.1      IQDDLA-SVRDVLTRNRETARKALDGTV-LEYQEPVVGVSVAWFRIAHPELTASGLQREL      347
MGH3938013.1        LDDDFC-SIRDVLMHNRETARRMLDGSI-LEYQEPVVPVSVAWFRINDPELTASELQSDL      347
MGB6162345.1        IDDDFC-SIRDVLMRNREIARNTLDGSI-LEYQEPVVPVSVAWFRIDHPKLTASELQSDL      347
                        :   :  .:  **   *    *   :   .*    **    :      .  :   *

MilM                SGHGIHVLPGTYFYWNEPSRGERYVRVALARDPGEFDASMARLRTLLARYA--------       389
RohP                SADGVYVLPGRYFYWSEPSKGDAYVRMALAREPEMFADAMALTRQVLDRHGR-------       393
MppP                QRHHVFALPCRQFHWAEPSDGDHMVRIALSRSTEPLEKSVQVLRTVLETR---------       376
Ind4                AKSQVGILPGDHFYWDDHETGSHFIRFALAREERWFATACGSLREALLARPELIEGAGS       383
WP_399337027.1      ADADVHVLPGTYFFWSRPERGERWMRVALARDPEEFAQSMRLLRKVVGRHG--------       388
WP_073229495.1      ADEGVHVLPGTFFYWSRPEQGERWIRVALARDPEDFAASMHRLREVVSRHG--------       388
WP_384049659.1      ADAGVHVLPGTFFFWSRPELGERWIRVALARDPEEFAASMRRLREVVLRHG--------       388
WP_149851715.1      LQNEVYVLPGTYFYWNDKSRGESFIRLALARDPQEFAGAMNAMREILARHER-------       409
WP_051969389.1      YVEGVYVLPGRYFYWSEPSRGDAYIRFALARDPEMFAEAMDITRQVLDRNVA-------       400
WP_116214779.1      TADGLYLLPGRYFYWSEPGKGESYLRIALARDPQMFSTAMTRLAAALDRHAR-------       399
WP_229328800.1      TGDGLYLLPGRYFYWSQPGKGESHLRVALARDPKMFSAAMSRLAQALDRHAR-------       399
MGH3938013.1        FGEGVYVLPGRYFYWSQPSKGDGFVRIALARDPEMFATAMNRLRRVLDCHGR-------       399
MGB6162345.1        LGEKVYVLPGRYFYWSQPSKGDSFVRIALARDPEMFATAMTRLRQVLDCHVW-------       399
                        :  **   *.*     *.  :*.**:*.   :  :       :
```

**Figure S19: Structural comparison of MilM.**
**(A)** Superposition of the AlphaFold model of MilM (grey) with RohP (PDB ID: 6C3C, orange), shown as ribbon representations.
**(B)** Close-up view of the PLP-binding pocket shown in cartoon representation, highlighting conserved residues (grey) from chains A and B interacting with PLP and L-Arg (cyan) and structured water molecules (violet).

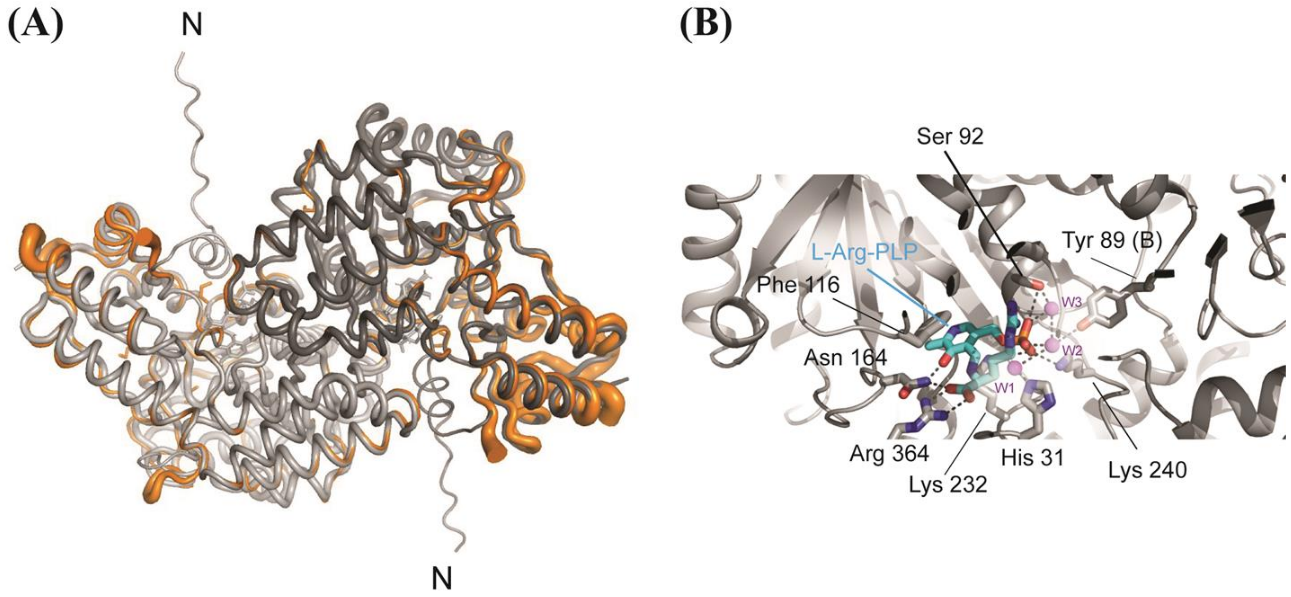

**Figure S20: LCMS analysis of the reactions of PLP-binding MilM variants with L-Arg substrate.** PLP-coordinating residues of MilM (**Figure 5C**) were mutated to Ala to construct the MilM variants including MilM-S92A, MilM-F116A, MilM-N164A, MilM-D195A, and MilM-K240A to evaluate the effects of these mutations on the MilM reaction. These variants were reacted with L-Arg under identical conditions as that of MilM-WT. LCMS analysis of the reaction mixtures of these variants showed no formation of expected products (**10** or **13**), while only the unreacted substrate L-Arg (**9**) mass (*m/z* = 175.1197 $[M + H]^+$) was detected. This indicates that these residues are important for the binding and anchoring of the PLP cofactor in the enzyme active site, which upon mutation leads to complete abolishment of the MilM activity.

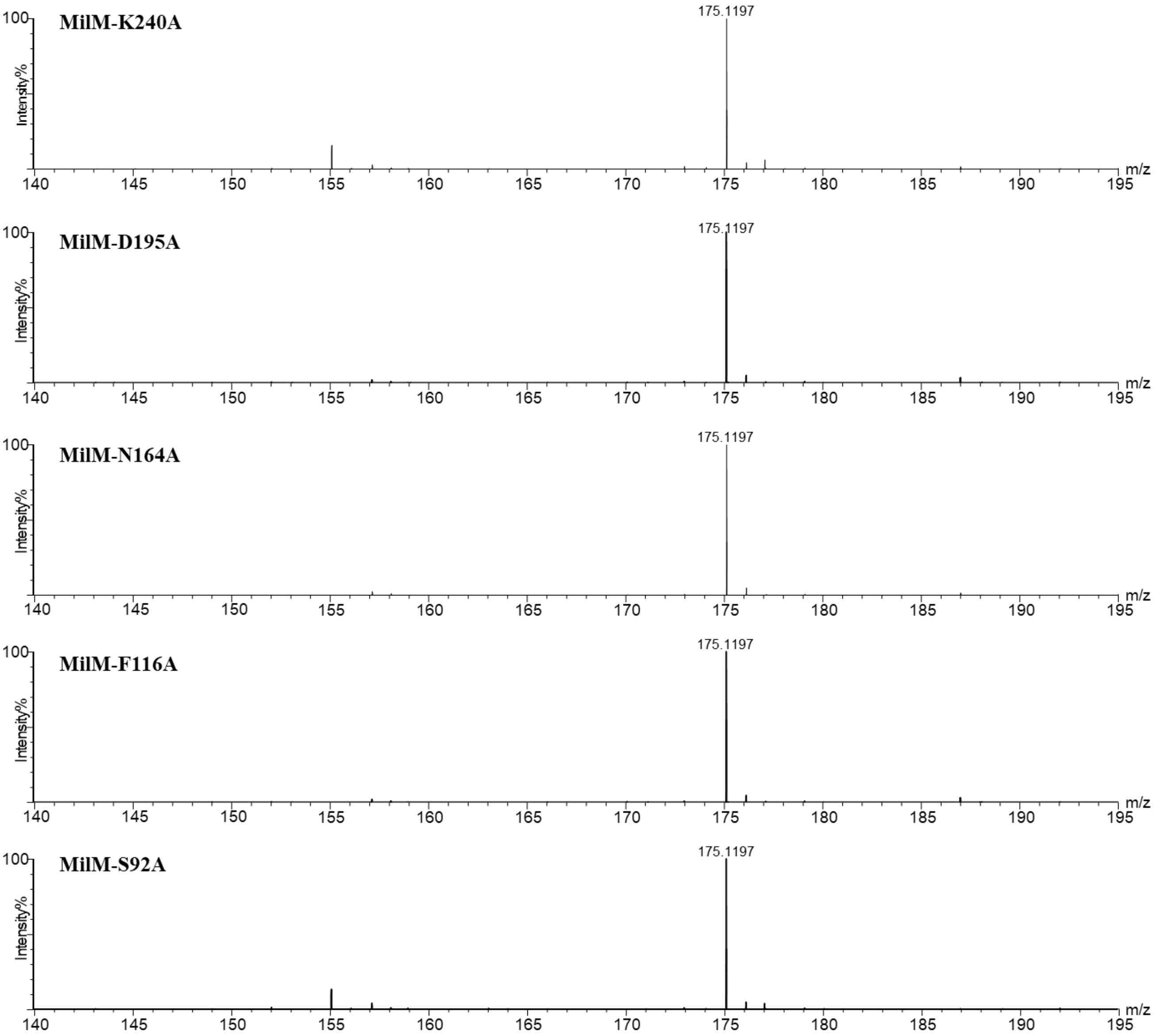

**Figure S21: LCMS analysis of the reactions of L-Arg binding MilM variants.** L-Arg-coordinating residues of MilM (**Figure 5C**) were mutated to Ala to construct the MilM variants including MilM-T14A, MilM-E17A, MilM-N118A and MilM-R364A to evaluate the effects of these Ala substitutions on MilM activity. These variants were reacted with L-Arg under identical conditions as that of MilM-WT. MilM-R364A variant showed no formation of products (**10** or **13**), while unreacted L-Arg (**9**) mass (*m/z* = 175.1197 $[M + H]^+$) was observed, indicating this residue is essential for proper positioning of L-Arg in MilM active site for further catalysis. On the other hand, MilM-T14A, MilM-E17A and MilM-N118A showed formation of only the oxo product (**13**, *m/z* = 174.0861 $[M + H]^+$), indicating these residues are indispensable for the formation of the hydroxylated product (**10**).

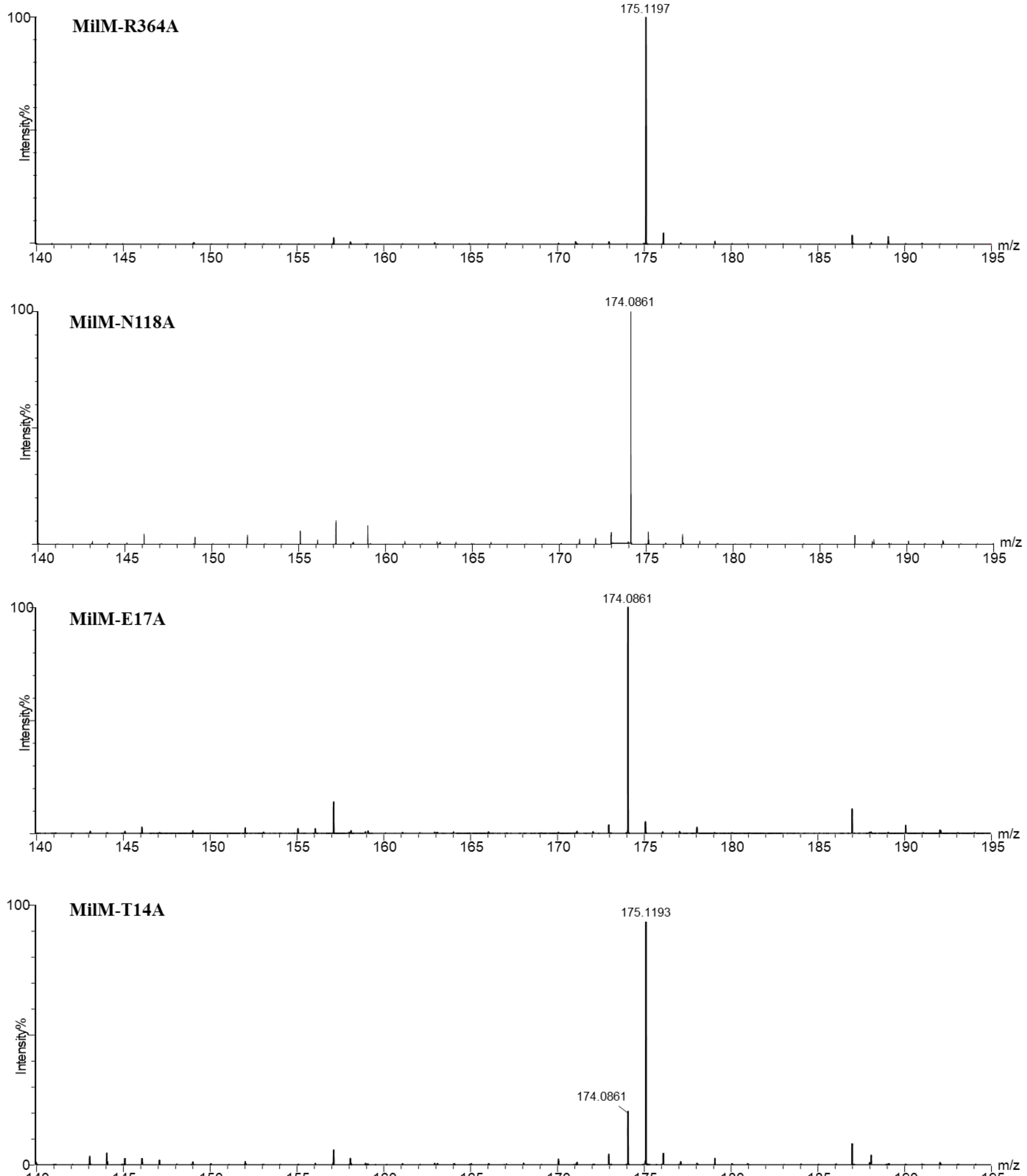

**Figure S22: LCMS analysis of the reactions of MilM variants from chain B binding with L-Arg-PLP in chain A.** Residue of chain B of MilM dimer which coordinate with the PLP cofactor bound in chain A was mutated to Ala to construct MilM-Y89A; and residues of chain B interacting with L-Arg in chain A ( through amide bonds) were mutated to Pro to construct MilM-T259P-S260P double variant to evaluate the effects of these mutations on the MilM reaction. These variants were reacted with L-Arg under identical conditions as that of MilM-WT. LCMS analysis of the reaction mixtures of MilM-Y89A and MilM-T259P-S260P showed no formation of products (**10** or **13**), while only the unreacted substrate L-Arg (**9**) mass (*m/z* = 175.1194 $[M + H]^+$) was detected. This indicates that these residues from one chain are important not only for the binding and anchoring of the PLP or L-Arg bound in the other chain but also necessary for dimer formation, which upon mutation led to abolishment of the MilM activity.

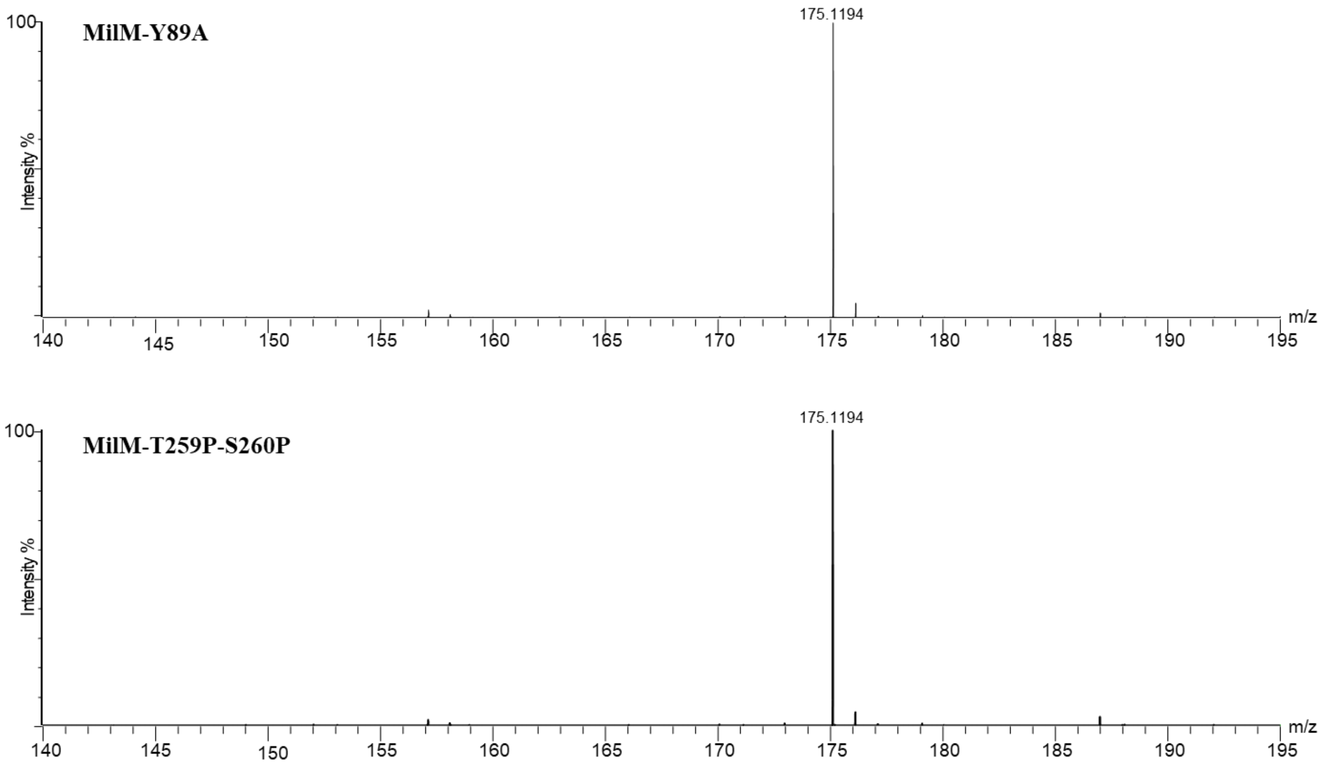

**Figure S23: Steady-state kinetic analysis of MilM-WT and its variants.**

**(A) Determination of $H_2O_2$ concentration using the ABTS-HRP assay:** Standard calibration curve of $H_2O_2$ (0 - 250 μM) with HRP (10 ng) and ABTS (10 mM), obtained by measuring absorbance at 418 nm, which was further used for quantifying hydrogen peroxide formation in MilM-catalyzed reactions.

Michaelis-Menten plots showing the catalytic parameters of: **(B)** MilM-WT, **(C)** MilM-T14A, **(D)** MilM-E17A and **(E)** MilM-N118A using L-Arg substrate. Reaction rates were determined by quantifying $H_2O_2$ formation using the ABTS-HRP assay.[4,6] Data were plotted and parameters were calculated using the KaleidaGraph software (**Table 1**).

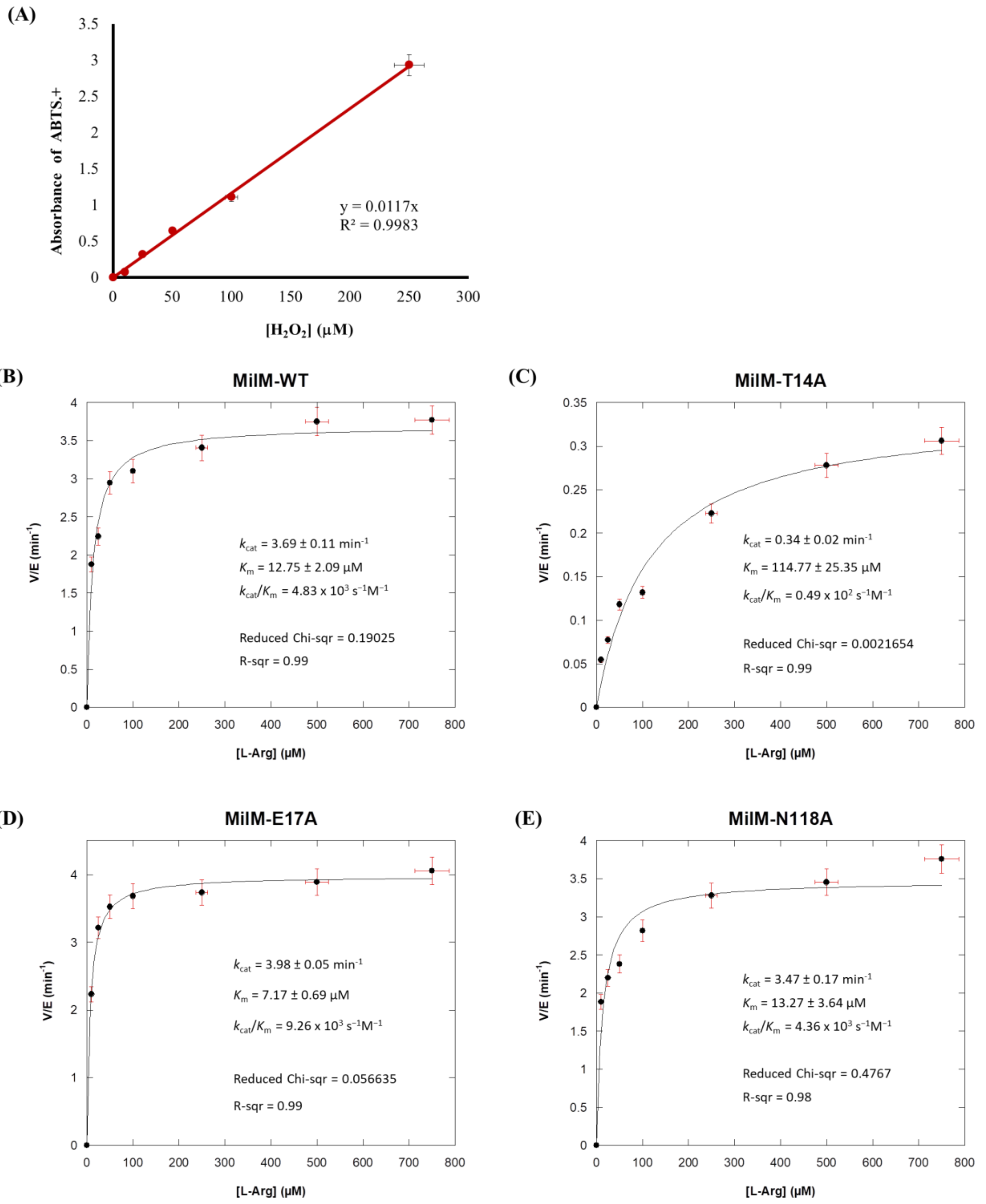

**Figure S24: LCMS analysis of the MilM assay using D-Arg as the substrate.** LCMS spectrum of the MilM reaction with D-Arg (enantiomer of the native substrate L-Arg) indicated that it was not accepted as a substrate by MilM. As observed, only the mass corresponding to unreacted D-Arg (*m/z* = 175.1187 $[M + H]^+$) was observed, while no masses corresponding to the expected products were detected. This experiment highlights the strict stereochemical requirement and substrate specificity of MilM (**Figure 7C**).

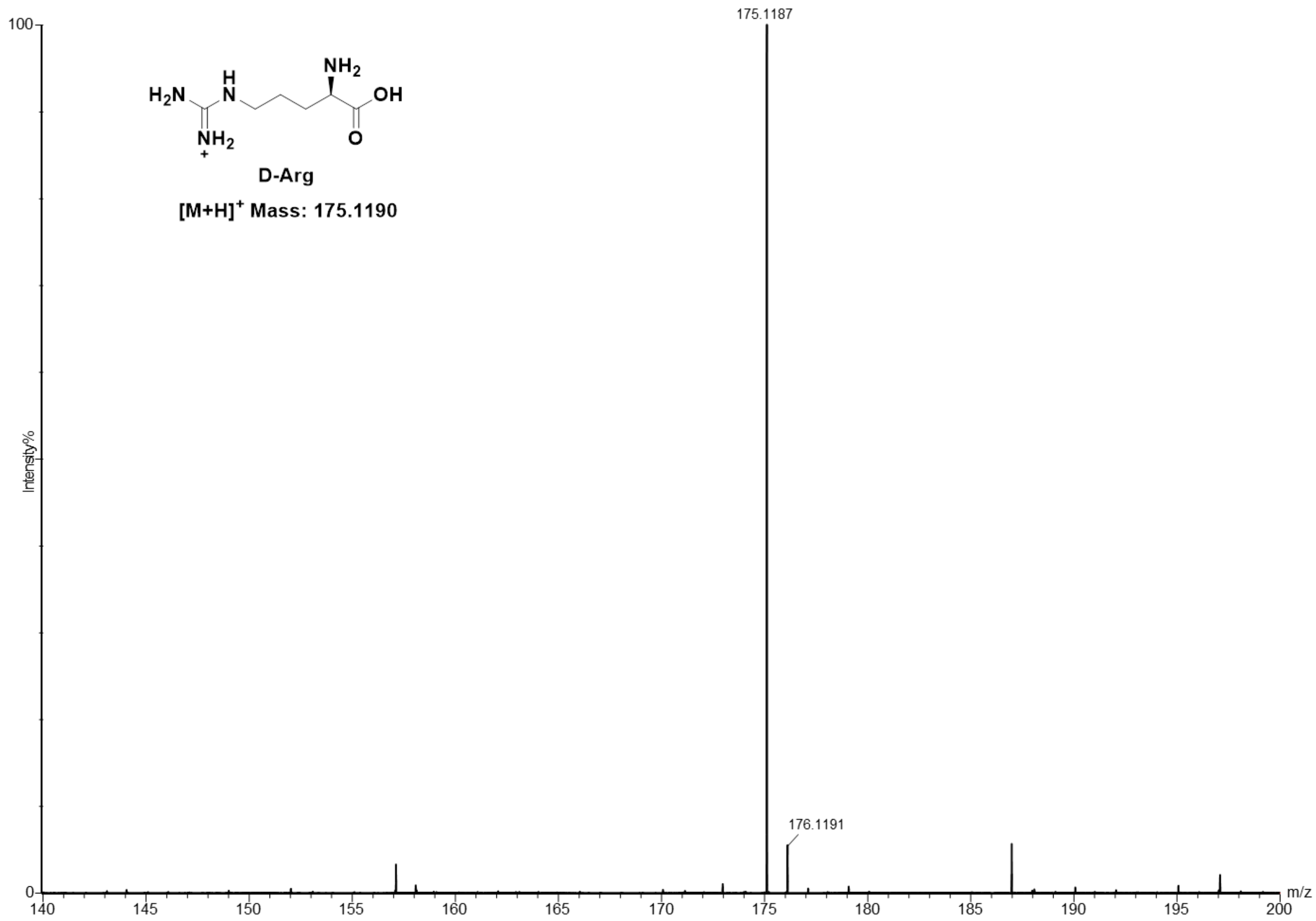

**Figure S25: Superimposition of MilM model structure with MppP and RohP crystal structures.**

**(A)** The crystal structure of MppP-D-Arg complex (PDB ID: 5DJ3, purple)[9] was superimposed on MilM model (grey) complexed with L-Arg-PLP (cyan).

**(B)** The product **10** from RohP-4-hydroxy-2-ketoarginine (HKA, orange) complex (PDB ID: 6C3A)[4] was superimposed with our L-Arg-PLP structure (cyan) from MilM model to predict the probable configuration of the hydroxyl group in product **10** (**Figure 3**).

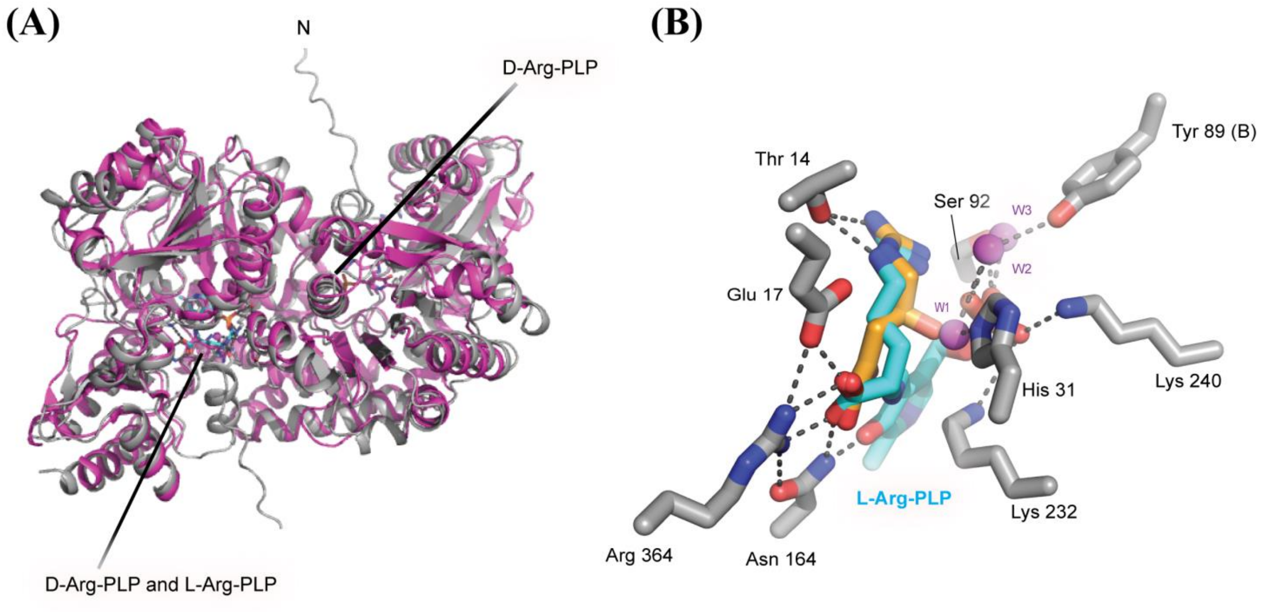

**Figure S26: The characteristics of the major principal components (PCs)**, including **(A)** the eigenvalue for each component, and **(B)** the individual and cumulative fractions of variance associated with each component. The top two principal components, PC1 and PC2, possess the highest eigenvalues and the greatest individual proportions of variance.

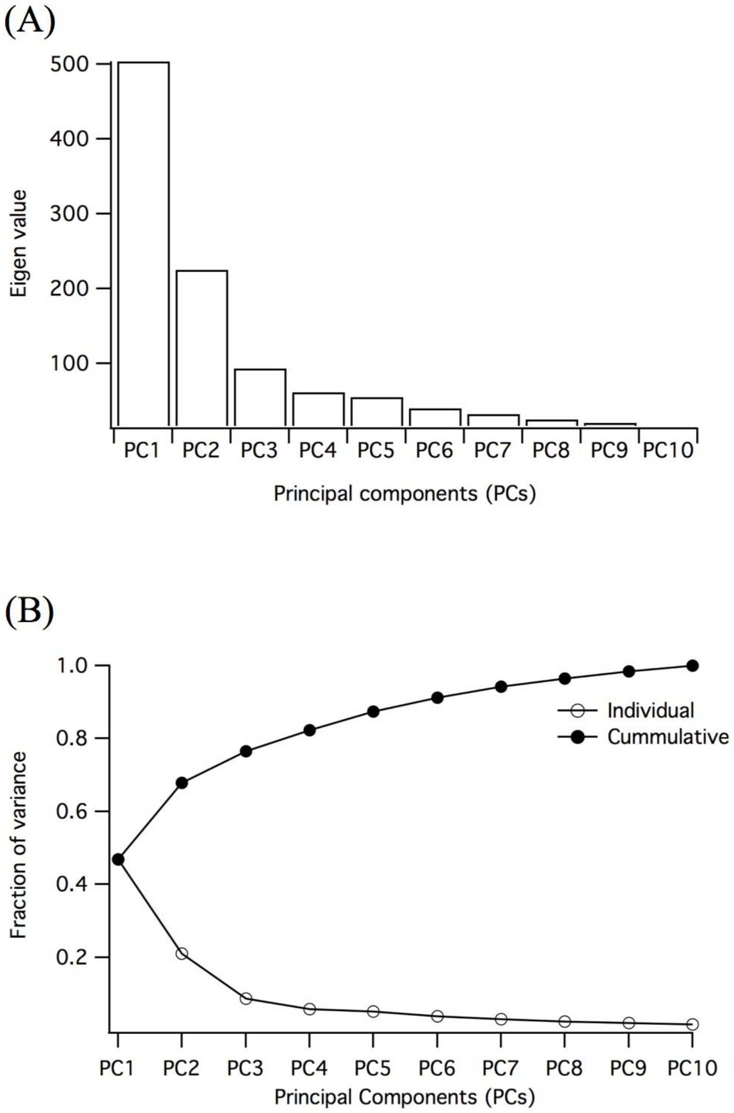

**Figure S27: Residue cross-correlation map based on RMSD of protein Cα atoms.**

**(A)** The cross-correlation for the complete dimer protein, consisting of residues of chain A (residues 1 to 389), and chain B (residues 390 to 778).

**(B)** The cross-correlation for only chain A residues of the dimer.

**(C)** The cross-correlation for solely chain B residues of the dimer.

**(D)** The cross-correlation between residues of chain A and chain B.

In the panels **A-D**, a positive value represents a positive correlation of motion between the residue pairs, a negative value represents negative correlation of motion, and zero represents a complete lack of correlation.

**(E)** Representative MD snapshot highlighting the dimer interface helices (chain A in cyan and chain B in blue), whose motions are positively correlated. Here, the red spheres and roman numbers (I to VI) on the interface helices represent the Cα atoms of the residues, identical to PCA: Ser23 (I) of chain A, Gln41 (II) of chain A, Arg58 (III) of chain B, Ser23 (IV) of chain B, Gln41 (V) of chain B, and Arg58 (VI) of chain A.

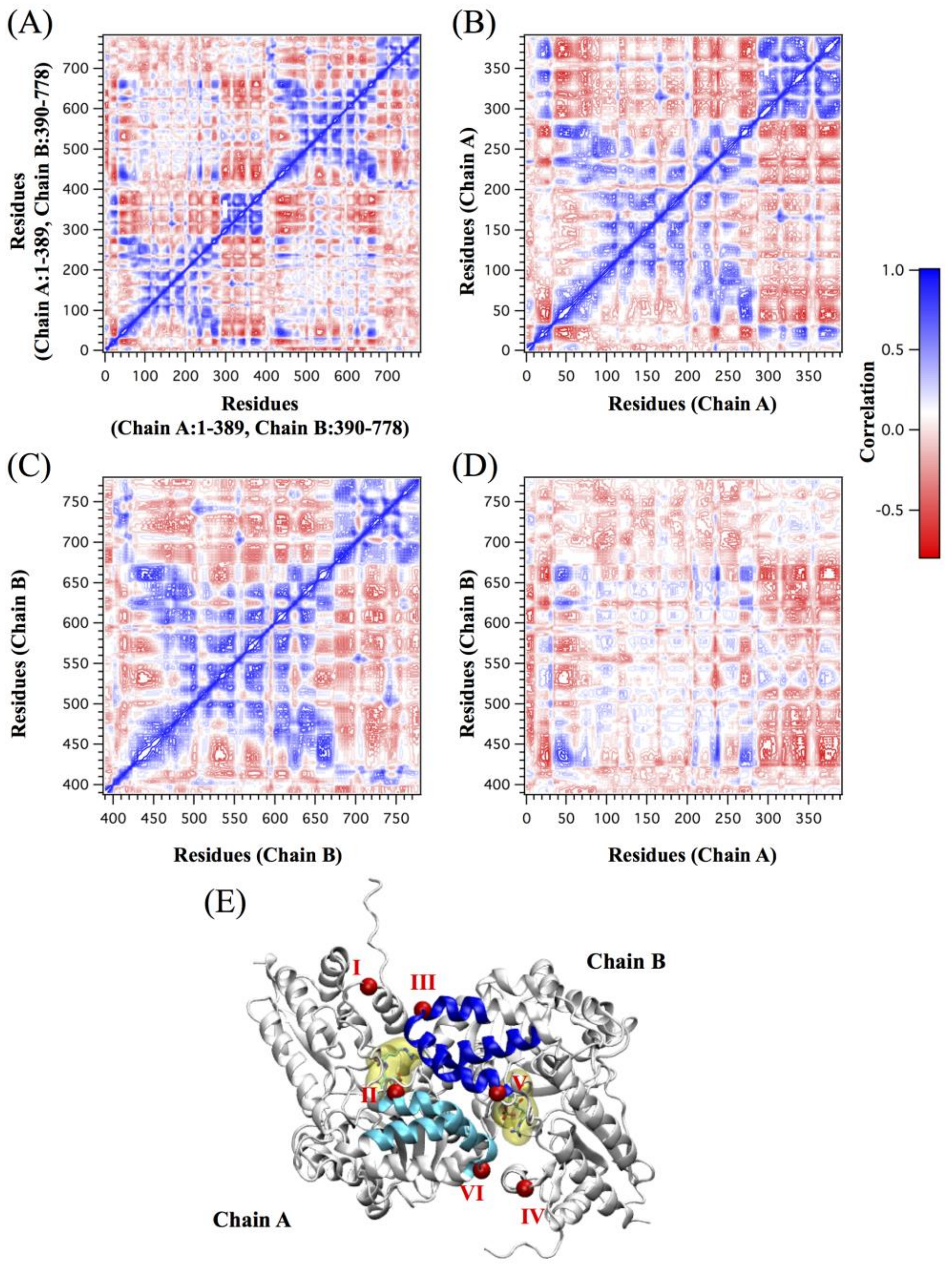

**Figure S28: 3D model structure of holo MilM enzyme.**
**(A)** MilM-PLP complex model was created based on RohP-PLP complex (PDB ID 6C3B).[4] This MilM structure bound with PLP represents a possible closed-lid like conformation.
**(B)** Superimposition of MilM-PLP complex (dark gray), RhoP-PLP complex (light pink) and RhoP-L-Arg-PLP complex (orange).

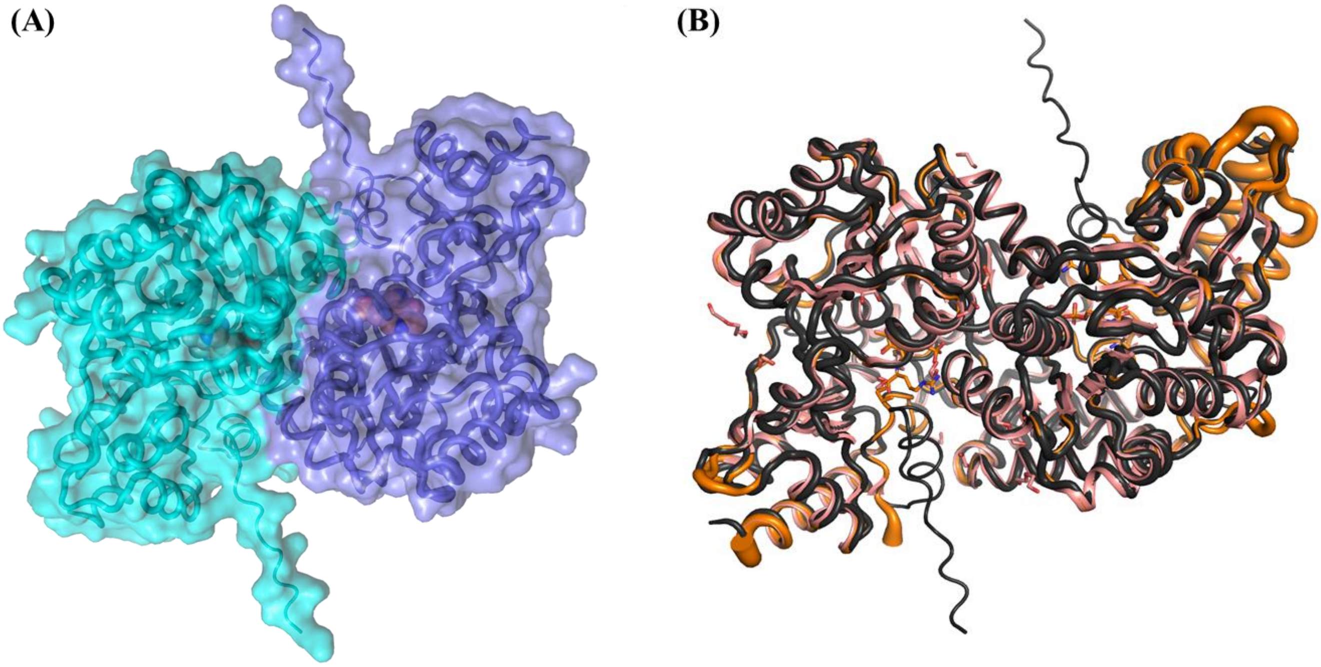
